# A Reproducibility Focused Meta-Analysis Method for Single-Cell Transcriptomic Case-Control Studies Uncovers Robust Differentially Expressed Genes

**DOI:** 10.1101/2024.10.15.618577

**Authors:** Nathan Nakatsuka, Drew Adler, Longda Jiang, Austin Hartman, Evan Cheng, Eric Klann, Rahul Satija

## Abstract

We assessed the reproducibility of differentially expressed genes (DEGs) in previously published Alzheimer’s (AD), Parkinson’s (PD), Huntington’s (HD), Schizophrenia (SCZ), and COVID-19 scRNA-seq studies. While transcriptional scores from DEGs of individual PD, HD, and COVID-19 datasets had moderate predictive power for case-control status of other datasets, genes from individual AD and SCZ datasets had poor predictive power. We developed a non-parametric meta-analysis method, SumRank, based on reproducibility of relative differential expression ranks across datasets, and found DEGs with improved predictive power. By multiple metrics, specificity and sensitivity of these genes were substantially higher than those discovered by dataset merging and inverse variance weighted p-value aggregation methods and had significant enrichment in snATAC-seq peaks and human disease gene associations. The DEGs revealed known and novel biological pathways, such as up-regulation of chaperone-mediated protein processing in PD glia and lipid transport in AD and PD microglia, and down-regulation of glutamatergic processes in AD astrocytes and glutamatergic neurons and synaptic processing and neuron projection genes in HD FOXP2 neurons. We find 56 DEGs shared amongst AD, PD, and HD, and validate *BCAT1* as down-regulated in AD mouse oligodendrocytes. Lastly, we evaluate factors influencing reproducibility of individual studies as a prospective guide for experimental design.

## Introduction

As single cell RNA-sequencing (scRNA-seq) technologies mature to process clinical samples, an increasing number of studies are profiling tissue from a multitude of disease states to identify cell type specific transcriptional alterations associated with pathophysiology and general development. scRNA-seq case-control studies have generated data on a multitude of neuropsychiatric diseases, such as multiple sclerosis^1–3^, schizophrenia (SCZ)^4–6^, major depressive disorder^7^, autism^8,9^, Parkinson’s disease (PD)^10–15^, alcohol use disorder^16,17^, Rett Syndrome^18^, vascular dementia^19^, and Huntington’s disease^20–23^, though all with relatively few individuals per study and often not in the same brain region. For Alzheimer’s Disease (AD) and COVID-19, however, scRNA-seq studies now have sample sizes in the hundreds^24–27^. These studies have uncovered known and novel biological pathways perturbed in these conditions that represent potential therapeutic targets.

Nevertheless, there has been concern for possible false positive results in these studies^28^, and thus the statistical methodology required to perform case-control studies across multiple cell types remains an area of active interest^29^. Initial studies implemented case-control analyses by performing differential-expression testing on individual cells. This approach treats each cell as an independent replicate, which fails to account for correlations across cells from the same individual and can lead to a large false-positive bias. Subsequent studies have dealt with these issues by using mixed models with individuals as a fixed or random effect^26^ or alternative regression models previously developed for bulk RNA-seq^30^ that can be used after pseudobulking clusters of single cells. Many of these methods can adequately control false positive rate and yet are sufficiently powered in analyses of simulated differentially expressed genes (DEGs). Nevertheless, there still has been substantial worry about potential false positives in DEG results due to technical artifacts or simply biological variation present in only small numbers of individuals (particularly for studies with smaller sample sizes). This issue is likely of particular relevance for many neuropsychiatric diseases due to the high transcriptomic heterogeneity of the brain at baseline^31^ and GWAS evidence for etiological diversity in many of these diseases^32^.

The field of human genetics, particularly genome-wide association studies (GWAS), can provide a model for the single-cell field in its high reproducibility^33^ and well-established meta-analysis methods for combining information across multiple datasets^34,35^. The typical GWAS meta-analysis usually applies an inverse variance weighting to aggregate the effect sizes and standard errors derived from each study to obtain final effect sizes and p-values for each genetic locus^36^. It is standard for new studies to have a separate test dataset to assess the reproducibility of significant genes found in the general analysis, testing for effect size and at least ensuring the same direction of effect in the test dataset. Now that many large-scale case-control scRNA-seq studies have been undertaken for several diseases, the field is in a strong position to develop standardized meta-analysis methods that combine information across multiple datasets with the goal of finding genes with transcriptional expression (and later other epigenetic loci) robustly associated with disease.

In this study we provide a systematic approach in this direction by first examining the reproducibility of 17 AD, 6 PD studies, 3 SCZ, 4 HD single-nucleus RNA-seq (snRNA-seq) studies and, as a positive control comparison due to its known strong transcriptional response, 16 single cell RNA-sequencing (scRNA-seq) COVID-19 studies. We find by several measures that a large fraction of the genes found to be differentially expressed in single AD and SCZ datasets do not reproduce in other AD and SCZ datasets, while genes found in PD, HD, and COVID-19 datasets have moderate reproducibility. To address this challenge, we introduce a new procedure for large-scale meta-analysis of scRNA-seq called SumRank that prioritizes the identification of DEGs that exhibit reproducible signals across multiple datasets and demonstrate that this approach substantially outperforms existing meta-analysis techniques in sensitivity and specificity of discovered DEGs. We demonstrate that SumRank identifies DEGs with high predictive power and reveals known and new biology. We use a mouse model of AD to validate a gene of particular interest and demonstrate for the first time that *BCAT1* is down-regulated specifically in oligodendrocytes, pointing to diminished branched chain amino acid metabolism in this cell type. We also show that SumRank DEGs are significantly enriched in genes associated with differentially accessible snATAC-seq peaks from a previous AD study. We find 56 DEGs shared amongst AD, PD, and HD. Moreover, we adapt SumRank to identify sex-specific DEGs. Finally, we assess factors that influence the reproducibility of an individual study’s results as a prospective guide for experimental design. Our work demonstrates the importance and potential for large-scale meta-analyses to draw robust biological conclusions, especially for neuropsychiatric disorders.

## Results

### Reproducibility of DEGs in individual datasets is poor in AD and SCZ and moderate in PD and COVID-19

We first compiled data from 17 snRNA-seq studies of AD prefrontal cortex (Supplementary Data File 1). We performed standard quality control measures on each dataset (Methods) and then determined cell types by mapping them to an established snRNA-seq reference of human cortical tissue (motor cortex) from the Allen Brain Atlas^37^ using the Azimuth toolkit^38^, which returns consistent cell type annotations for all datasets at multiple levels of resolution (Figure 1). We then performed pseudobulk analyses for broad cell types, obtaining transcriptome-wide gene expression means or aggregate sums for each gene within each of the 7 cell types within each individual (aggregate sums were used for DESeq2^30^ analyses while means were used for all other analyses). We used these values to identify celltype-specific DEGs for AD vs. control samples in downstream analyses. Leveraging pseudobulk values removes the inherent lack of independence that characterizes multiple cells from the same individual, which would otherwise lead to substantial false positives for standard single-cell differential expression workflows. We also performed the same pipeline for 6 snRNA-seq studies of PD midbrain and 4 studies of HD caudate, determining cell types by mapping to the highest quality dataset (because there are no midbrain or caudate Azimuth atlases), and 3 snRNA-seq studies of SCZ prefrontal cortex. As a control experiment for a disease phenotype with a well-described and strong transcriptional response, we repeated this process for 16 scRNA-seq studies from PBMC samples from COVID-19 patients and healthy controls (Supplementary Data File 1 contains information about all datasets).

**Figure 1.**
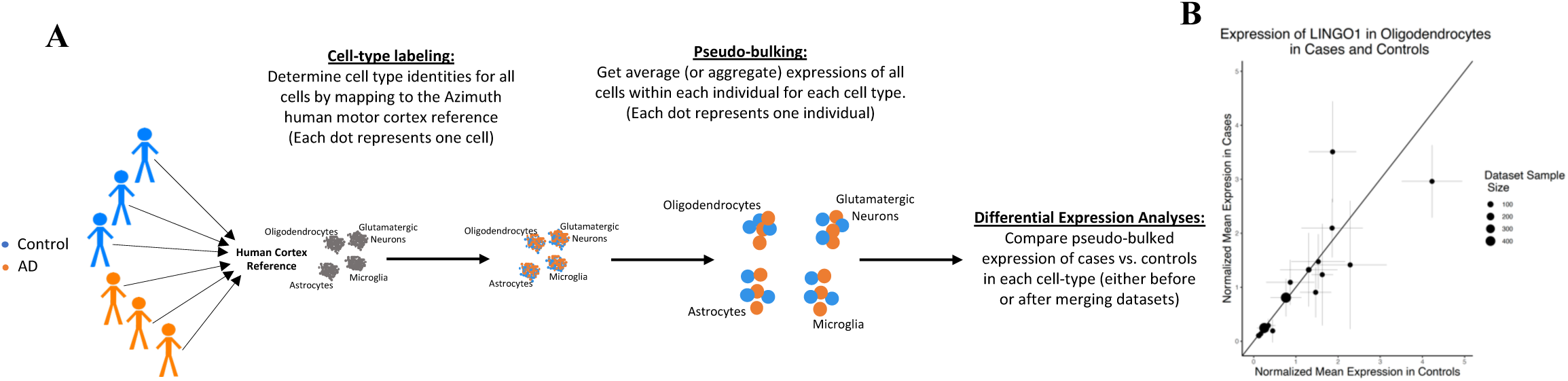
Schematic of the procedure for obtaining differentially expressed genes. **A)** Schematic of mapping cells to determine cell types, pseudobulking, and obtaining cell type specific differential expression (some cell types are removed for clarity). Orange represents AD individuals or cells, and blue represents controls. The first two sets of dots represent cells while the third set of dots represent individuals (the sum or mean expression across all cells in a particular cell type for that individual). **B)** Example of a gene, *LINGO1*, previously highlighted as up-regulated in oligodendrocytes that was shown to not be up-regulated in most datasets. Values above the line (intercept=0, slope=1) are up-regulated, while values below the line are down-regulated. Error bars are standard deviations in all plots. Violin plots of the expression of *LINGO1* in each individual across all datasets is shown in Supplementary Figure 1.

We evaluated the reproducibility of DEGs between diseased and control samples by calculating DEGs based on pseudobulked values for each cell type and utilized the DESeq2^30^ package for DEG detection using a q-value based FDR cutoff of 0.05, because DESeq2 with pseudo-bulking has been shown to have good performance in terms of specificity and sensitivity relative to other methods^39^. Strikingly, when using this criterion over 85% of the AD DEGs we detected in one individual dataset failed to reproduce in any of the 16 others (Supplementary Table 1). Few genes (<0.1%) were consistently identified as DEGs in more than three of the 17 AD studies, and none were reproduced in over six studies. While we observed improved reproducibility in PD, HD, and COVID-19 datasets, we still failed to observe a single gene that was independently detected as exhibiting consistent cell type-specific differential expression in more than 4 of the 6 PD, 10 of 16 COVID-19, or 1 of the 3 SCZ studies (Supplementary Tables 2-5; note: the SCZ low overlap here was driven by having extremely few DEGs with this criteria, see Supplementary Note).

We frequently observed that genes that were identified as DEGs in multiple studies tended to rank highly even in studies where they failed to pass the required threshold. For example, when we instead looked at the reproducibility of the top 200 genes for each cell type (ranked by p-values), some genes were found in up to 9 of 17 AD, 6 of the 6 PD, 11 of 16 COVID-19, and 3 of the 3 SCZ datasets (Supplementary Tables 6-10). This suggests that at least some of the variability in DEG identification is driven by a lack of statistical power for any individual study. This further highlights the limitation of depending solely on one study to reliably identify DEGs that will reproduce in other studies, especially in intricate diseases such as AD. Illustrating this, we examined the gene *LINGO1*, a negative regulator of myelination previously spotlighted as a crucial oligodendrocyte DEG in a recent AD review^40^. While we reproduced this finding in a few individual datasets, our broader analysis suggests that *LINGO1* was not consistently up-regulated in oligodendrocytes in the majority of datasets and was even down-regulated in several studies (Figure 1 and Supplementary Figure 1), highlighting challenges associated with identifying bona-fide and reproducible DEGs.

We also tested reproducibility by assessing the ability of the DEG sets from individual studies to differentiate between cases and controls in other studies. To standardize cross-dataset comparisons, we identified the same number of top-ranked DEGs (ranked by p-value without requiring an explicit FDR cutoff) and derived a transcriptional disease score for each cell type in each individual. We obtained these by leveraging the UCell score^41^—a method that determines the relative rank of genes compared to others in a dataset. Our findings revealed that the DEGs identified by any individual AD dataset were not highly effective in predicting case-control status in other AD datasets (mean AUC of 0.68) or SCZ datasets (mean AUC of 0.55), though we observed improved power for PD, HD, and COVID-19 studies (mean AUCs of 0.77, 0.85, and 0.75, respectively) (Extended Data Tables 1-3, Table 1, Supplementary Table 11-12). Using a fixed FDR cutoff as an alternative for deriving transcriptional disease scores generally led to even poorer results (Supplementary Tables 13-15). However, we observed that DEGs identified by the 3 AD studies with a large number of individuals (>150 cases and controls each) exhibited superior predictive performance in alternative datasets (AUCs of 0.75 to 0.80) (Extended Data Table 1).

**Table 1.** Comparisons of individual datasets and different meta-analysis methods in their predictive performances. For all analyses here the DEG lists included the same number of top genes (based on the number of SumRank genes with -log10(p-value) at a cutoff identified in the main text). RCA Gene List is the list of genes ranked by their individual ability to distinguish cases from controls in all datasets (see text and Methods for more details). Relative Classification Accuracy is the mean AUC of individual genes in their ability to distinguish diagnosis status in each dataset, normalized within each disease. Mean absolute log2fc were from comparisons of cases and controls in each dataset. * indicates that the RCA Gene List is likely less reliable in SCZ due to the low number of datasets.

| Disease | Gene Set Type | Mean AUC when using DEGs as a Group to Predict Diagnoses of Left-Out Datasets | Specificity: Percentage of DEGs in Top 10% of RCA Gene List | Mean Relative Classification Accuracy of Individual DEGs | Mean absolute log2fc of individual genes between cases and controls in each dataset |
| --- | --- | --- | --- | --- | --- |
| AD | Mean of Individual Datasets | 0.67 | 34 | 43.4 | 0.15 |
| AD | SumRank | 0.78 | 73 | 64.4 | 0.33 |
| AD | Merge | 0.78 | 41 | 55.6 | 0.32 |
| AD | Inverse Variance | 0.74 | 21 | 43.6 | 0.20 |
| COVID-19 | Mean of Individual Datasets | 0.75 | 58 | 40.4 | 0.37 |
| COVID-19 | SumRank | 0.91 | 78 | 58.6 | 0.79 |
| COVID-19 | Merge | 0.90 | 72 | 57.0 | 0.97 |
| COVID-19 | Inverse Variance | 0.88 | 42 | 46.5 | 0.72 |
| PD | Mean of Individual Datasets | 0.77 | 57 | 53.0 | 0.31 |
| PD | SumRank | 0.88 | 87 | 71.0 | 0.52 |
| PD | Merge | 0.84 | 68 | 63.2 | 0.63 |
| PD | Inverse Variance | 0.85 | 57 | 57.6 | 0.41 |
| SCZ | Mean of Individual Datasets | 0.55 | 37* | 44.3* | 0.24 |
| SCZ | SumRank | 0.62 | 51* | 53.4* | 0.35 |
| SCZ | Merge | 0.52 | 23* | 43.8* | 0.26 |
| SCZ | Inverse Variance | 0.56 | 21* | 38.4* | 0.29 |
| HD | Mean of Individual Datasets | 0.85 | 48 | 50.2 | 0.58 |
| HD | SumRank | 0.84 | 68 | 62.3 | 0.96 |
| HD | Merge | 0.85 | 48 | 56.3 | 1.18 |
| HD | Inverse Variance | 0.83 | 18 | 37.9 | 0.57 |

We wanted to evaluate reproducibility on a per gene level rather than at only a combined gene set level, so we also tested the ability of individual DEGs to classify disease status for all samples across all studies. While the expected classification power for a single gene is expected to be low, we reasoned that the relative ranking of the genes could serve as an informative metric for evaluating different DEG sets. We therefore developed a single-gene metric of classification power (‘Relative Classification Accuracy’), which was the normalized AUC of an individual gene for predicting case-control status (see Methods for more details); we ranked the genes by this metric and named the ranked list ‘RCA Gene List’. We identified the top 10% of genes in the RCA Gene List (1,520, 1,780, 1,107, 1,843, and 1,742 for AD, PD, COVID-19, HD, and SCZ, respectively), reasoning that bona fide DEGs should generally fall within this set. However, when returning to the sets of DEGs identified by individual datasets, we observed poor overlap within this list (mean of 34%, 57%, 58%, 48%, and 37% for AD, PD, COVID-19, HD, and SCZ). Even when examining the three largest AD datasets, we still observed poor performance for individual genes (37-51% in the top 10% of the RCA Gene List). Taken together, we conclude that analysis of individual datasets often fails to identify DEGs between cases and controls that reproduce in additional studies, and that this problem is exacerbated for diseases with more subtle or more heterogeneous transcriptional phenotypes such as AD. We therefore sought to explore approaches for meta-analysis that would leverage datasets from multiple studies to identify robust DEGs.

### A non-parametric meta-analysis uncovers DEGs with strong reproducibility across datasets

We tested two standard meta-analysis strategies. As one approach, we merged pseudobulk profiles together from all datasets and then conducted a differential expression analysis using DESeq2 while including the dataset ID as a batch covariate. As an alternative approach, we incorporated an inverse variance meta-analysis, a conventional approach for amalgamating GWAS summary statistics. For this, we fused the effect sizes and standard errors from each dataset’s DESeq2 results using metagen^42^. We used both approaches to calculate consensus DEG sets.

We found that the DEG sets identified by the merge and inverse variance strategies outperformed the DEG sets identified from individual dataset analyses. As an example, both methods correctly failed to identify significant differential expression for *LINGO1*. More broadly, the DEG gene sets had improved predictions of case control status in omitted datasets with mean AUCs of 0.78 and 0.74, respectively, for AD and similar improvements for PD and COVID-19. Yet, even with enhanced AUCs, numerous genes identified by the meta-analyses showcased limited specificity, with less than 42% ranking within the top 10% of the RCA Gene list for AD (Table 1; Figure 2). When examining the reason for this low specificity, we found an inherent weakness with these approaches: if a gene was highly significant in a small minority of datasets it would often pass significance thresholds after meta-analysis, even if no signal was observed in the remainder of the studies. We conclude that meta-analysis can improve the robustness of DEG identification, but existing methods remain prone to false positive identification.

**Figure 2.**
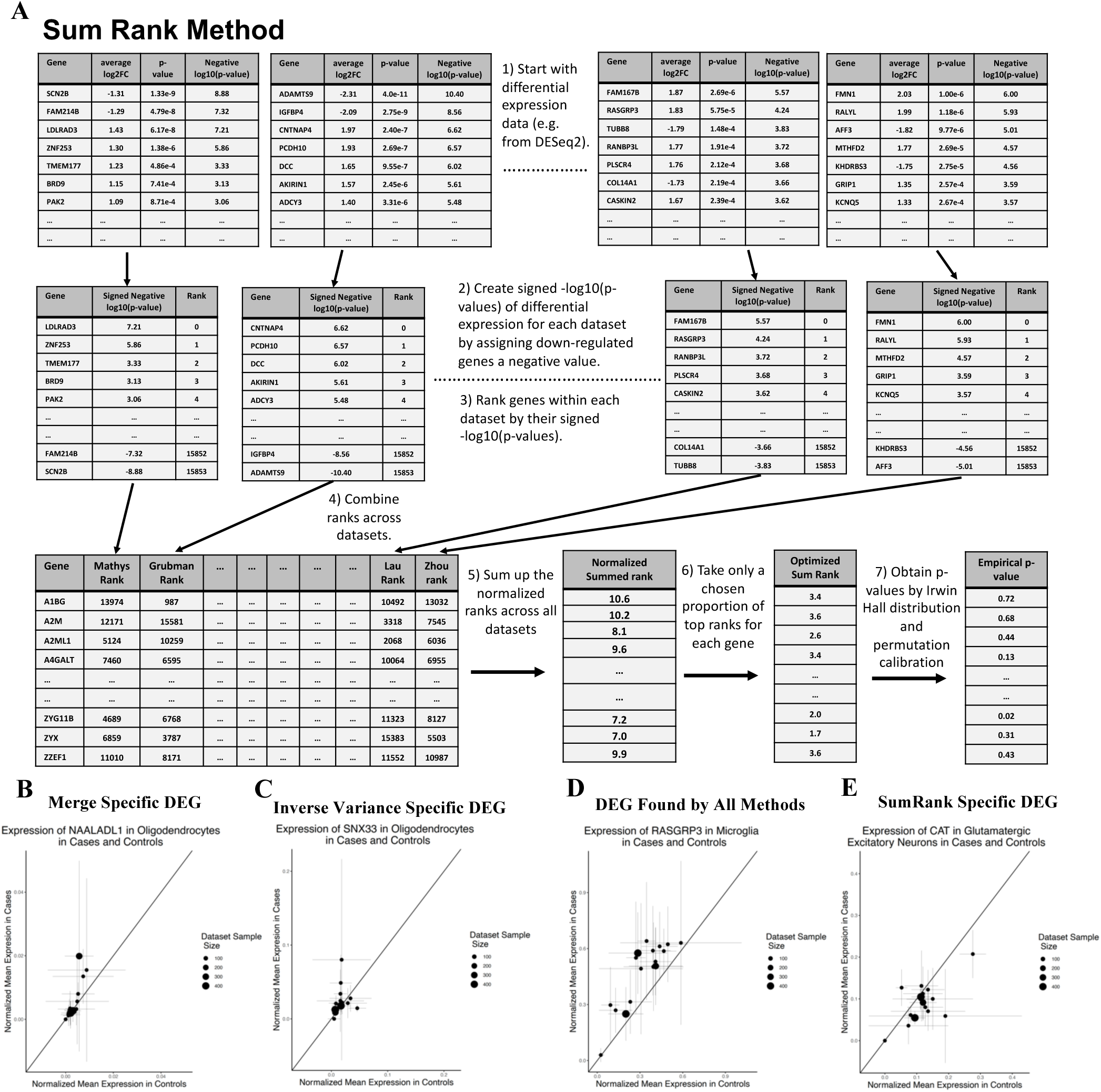
Schematic and results of the SumRank method. **A)** Cartoon of the SumRank method: scoring each gene based on the sum of their ranks across all datasets (see text and Methods for more details). **B)** Example of a gene (*NAALADL1*) putatively up-regulated in AD oligodendrocytes based on the Merge method that is likely a false positive (very low expression and high variance). **C)** Example of a gene (*SNX33*) putatively up-regulated in AD oligodendrocytes based on the Inverse Variance method that is likely a false positive. **D)** Example of a gene (*RASGRP3*) up-regulated in AD microglia based on all methods. **E)** Example of a gene (*CAT*) down-regulated in AD glutamatergic neurons based on the SumRank method that was not discovered by the Merge or Inverse Variance methods. Values above the line (intercept=0, slope=1) are up-regulated, while values below the line are down-regulated. Error bars are standard deviations in all plots. Violin plots of the expression of *RASGRP3* in each individual across all datasets are shown in Supplementary Figure 2.

To address the issue of genes found with low reproducibility across datasets we developed a novel, non-parametric meta-analysis method, which we call SumRank, that explicitly prioritizes reproducibility across multiple studies yet does not impose strict statistical cutoffs for any individual study (Figure 2). This method takes the results of dataset-specific DE analysis, calculates ranks (p-value based) for each gene in each dataset, and sums these ranks together across datasets. The resulting sum reflects a statistic that prioritizes genes that consistently exhibit evidence of differential expression across datasets. Given that requiring strong signals across all datasets can be overly strict—especially with large dataset numbers—we adjusted the SumRank statistic to consider only the ranks from a percentage of datasets. We set this percentage to 100% for meta-analyses based on fewer numbers of studies (PD, HD, and SCZ). For larger meta-analyses, we set this percentage based on cross-validation (65% and 55%, for AD and COVID-19, respectively), but found that our results remained consistent regardless of the exact threshold selected (Supplementary Data File 2). While the theoretical distribution of the SumRank statistic follows the Irwin-Hall distribution (see Methods), using only a subset of datasets causes deviations from this distribution. To address this, we empirically modeled the distribution by performing 10,000 random permutations of case-control status. This allowed us to apply the identical differential expression and meta-analysis process to create a null distribution of SumRank statistics, which we used to compute empirical p-values.

When we applied a Benjamini-Hochberg FDR cutoff of 0.05, we obtained 521 genes (394 up- and 127 down-regulated across 7 cell types) as significant in AD, 1,597 genes in PD (1,540 up- and 57 down-regulated across 8 cell types), 1026 genes in HD (628 up- and 398 down-regulated across 15 cell types), and 1,638 genes (1,432 up- and 206 down-regulated across 8 cell types) in COVID-19, but 0 genes in SCZ (Supplementary Data Files 3-7; Supplementary Figures 3-4). With this cutoff some cell types had no DEGs, so we looked for uniform -log10(p-value) cutoffs that led to gene sets that maximized the ability to predict case-control status in left out datasets. We found that for AD a -log10(p-value) cutoff of 3.65 produced 814 genes (502 up- and 312 down-regulated) with an AUC of 0.78, for PD a cutoff of 3.35 produced 1,527 genes (1,232 up- and 295 down-regulated) with an AUC of 0.88, for HD a cutoff of 3.30 produced 1,555 genes (740 up- and 815 down-regulated) with an AUC of 0.84, for COVID-19 a cutoff of 3.90 produced 937 genes (730 up- and 207 down-regulated) with an AUC of 0.91, and for SCZ a cutoff of 3.40 produced 98 genes (50 up- and 48 down-regulated) with an AUC of 0.62, all higher AUCs than those from individual datasets or either of the previously tested meta-analysis procedures. Most encouragingly, we found that more than 73% of the AD DEGs fell within the top 10% of the RCA gene list, suggesting high specificity for individually identified genes. For standardization, we used the same number of genes from the SumRank meta-analyses (814, 1,527, 1,723, 937, and 98) for all other analyses reported in this paper. When thresholds based on corrected p-values of the meta-analysis outputs were used (either through Bonferroni or q-value based FDR), it was not possible to find uniform p-value cutoffs that allowed reasonable comparisons between the meta-analysis methods (in Extended Data Figure 1 we show plots with the q-value based FDR thresholds for AD).

To assess whether clinical covariates affected reproducibility, we performed both DESeq2 and a logistic regression while regressing out all relevant covariates available for each dataset (sex, age, PMI, RIN, education level, ethnicity, language, age at death, batch, fixation interval, nCount_RNA, and nFeature_RNA). We did not observe any improvement in reproducibility with these analyses (Supplementary Table 16), suggesting that the datasets were generally well-controlled experiments with no systematic biases between cases and controls. We also used a newly developed single-cell differential expression method (Memento^43^) to assess whether this might improve reproducibility relative to using DESeq2 with pseudo-bulk data. We found that this also had slightly lower reproducibility relative to DESeq2, though SumRank combination of the Memento results improved reproducibility relative to individual Memento results (Supplementary Table 17). We also performed analyses at an increased cell resolution, looking at more fine-grained subsets of the cortical neurons. We found 1,611 significant (FDR<0.05) DEGs (155 up-regulated and 1,456 down-regulated) across the 14 neural cell types and 1,408 at a -log10p-value cutoff of 3.65 (330 up-regulated and 1078 down-regulated; Supplementary Data File 3). The genes found at the broader neuron types were found repeatedly across the more specific types (e.g. *ADAMTS2*, *SCGN*, *HES4*, *CIRBP*, *PDE10A*, *VGF*), but the genes only found in the higher resolution types could represent true cell type-specific DEGs. However, when we used the more specific DEGs together with the glial genes we obtained slightly decreased reproducibility (AUC=0.77 for AD and 0.59 for SCZ). We believe this is potentially due to the predictive signal now being diluted across more cell types (increased model parameters), less accurate cell type mapping, or increasing missingness in the datasets at the higher cell resolution. We thus continued our subsequent analyses at the broader cell resolution.

To more carefully benchmark SumRank against alternative methods for meta-analysis, we compared the AD DEG gene sets for each method. We first focused on the 81 genes found across all three methods (SumRank, merge, Inverse Variance), reasoning that this represented a gold-standard DEG set (example in Figure 2D and Supplementary Figure 2). Consistent with this, we found that these genes tended to exhibit high Relative Classification Accuracy (Figure 3). They also exhibited medium-high levels of expression (suggesting that they could be accurately quantified in individual datasets), and high mean absolute log2(fold-change) in comparisons of case vs control status in each dataset. We next examined genes that were identified by only a subset of methods. For example, we examined the genes that were identified by either the merge or inverse-variance methods (or both), but not by the SumRank method. In contrast to our gold-standard gene set, these genes exhibited low RCA and reduced log2(fold-change) (Figure 3). They also tended to be lowly expressed. Taken together, these results suggest that many of these genes likely represent false positives, and that the SumRank method correctly failed to identify them as DEGs. In contrast, the genes identified by SumRank (either exclusively or with one of the other meta-analysis methods) closely resembled the gold standard gene set. We conclude that the SumRank method exhibits superior performance by avoiding both false-positives and false-negatives, excluding genes that do not reproduce across multiple datasets but also sensitively identifying genes whose aggregate signal across multiple datasets is reliably supportive of differential expression between cases and controls.

**Figure 3.**
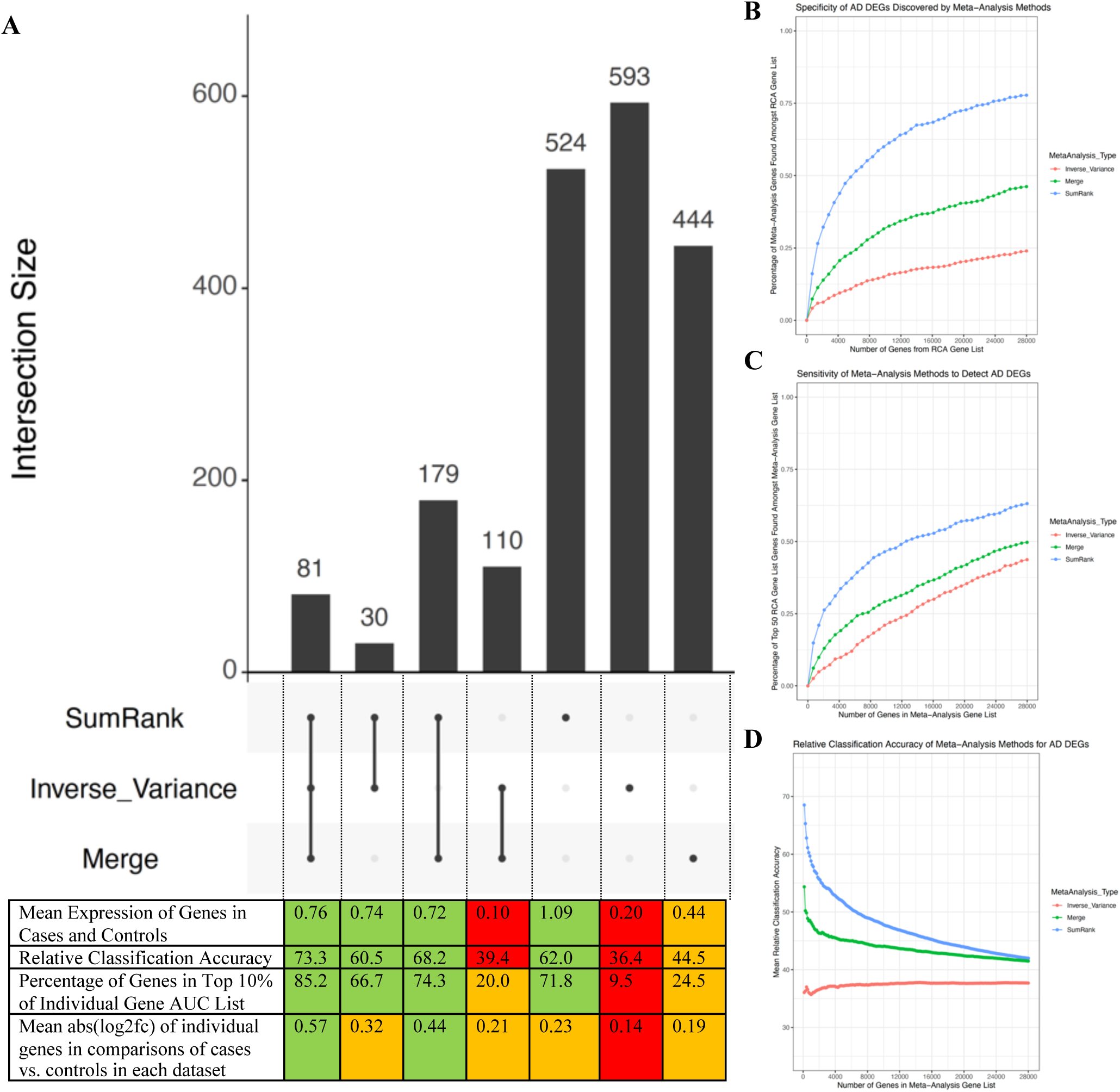
Sensitivity and Specificity of SumRank meta-analysis is better than merge and inverse variance methods. **A)** UpSet R plot^44^ showing intersection of AD genes discovered between the meta-analysis methods, the mean expression of the genes, relative classification accuracy (the normalized mean AUC of the individual genes in ability to predict diagnoses in all datasets), percentage of genes in top 10% of RCA Gene List, and mean abs(log2fc) from comparisons of cases vs. controls in each dataset. Color coding is based on the relative quality of the value, with green indicating the best values, orange indicating moderate, and red indicating poor. Comparisons of meta-analysis methods in their **B)** specificity, as measured by the percentage of their genes that intersect with the RCA Gene List (at different thresholds) with the same number of genes used in all meta-analyses (based on the 814 SumRank genes with -log10(p-value)>3.65), **C)** sensitivity, as measured by the percentage of the top 50 RCA Gene List genes found amongst the meta-analysis DEGs at different thresholds, and **D)** Relative Classification Accuracy, the mean AUC of individual genes in their ability to distinguish diagnosis status in each dataset (in this case averaged over all genes in the gene set). On the x-axes of B-D, the number of genes are spread evenly across up and down-regulated and all the different cell types. Similar plots for COVID-19 are shown in Extended Data Figure 6.

Examining the AD SumRank gene sets, we found that microglia, oligodendrocytes, GABA-ergic neurons, and astrocytes exhibited a greater number of up-regulated genes compared to down-regulated ones. In contrast, glutamatergic neurons demonstrated more down-regulated genes than up-regulated, consistent with earlier findings^45,46^ (Figure 4, Extended Data Figures 2-3, and Supplementary Figure 5). For AD, we detected the highest number of up-regulated genes in astrocytes. In contrast, for PD the highest number of up-regulated genes were in oligodendrocytes, while for HD, FOXP2 neurons had a substantially higher number of down-regulated genes than other cell types. For all diseases, over 75% of the DEGs were restricted to a single cell type (Supplementary Figure 6). When examining the correlations of -log(p-value)s for each cell type, we observed that cell types with greater similarities showed higher correlation (Supplementary Figure 7). Furthermore, using the SumRank genes, we identified some predictive capacity for disease specificity (Braak score) within AD patients (r=0.32) when compared to separate datasets (mean r=0.12) (Supplementary Data File 3). However, we found no predictive ability related to COVID-19 severity (r=0.03) (Supplementary Data File 5). This was anticipated, as the severity of COVID-19 has minimal relation to transcriptional response^47^.

**Figure 4.**
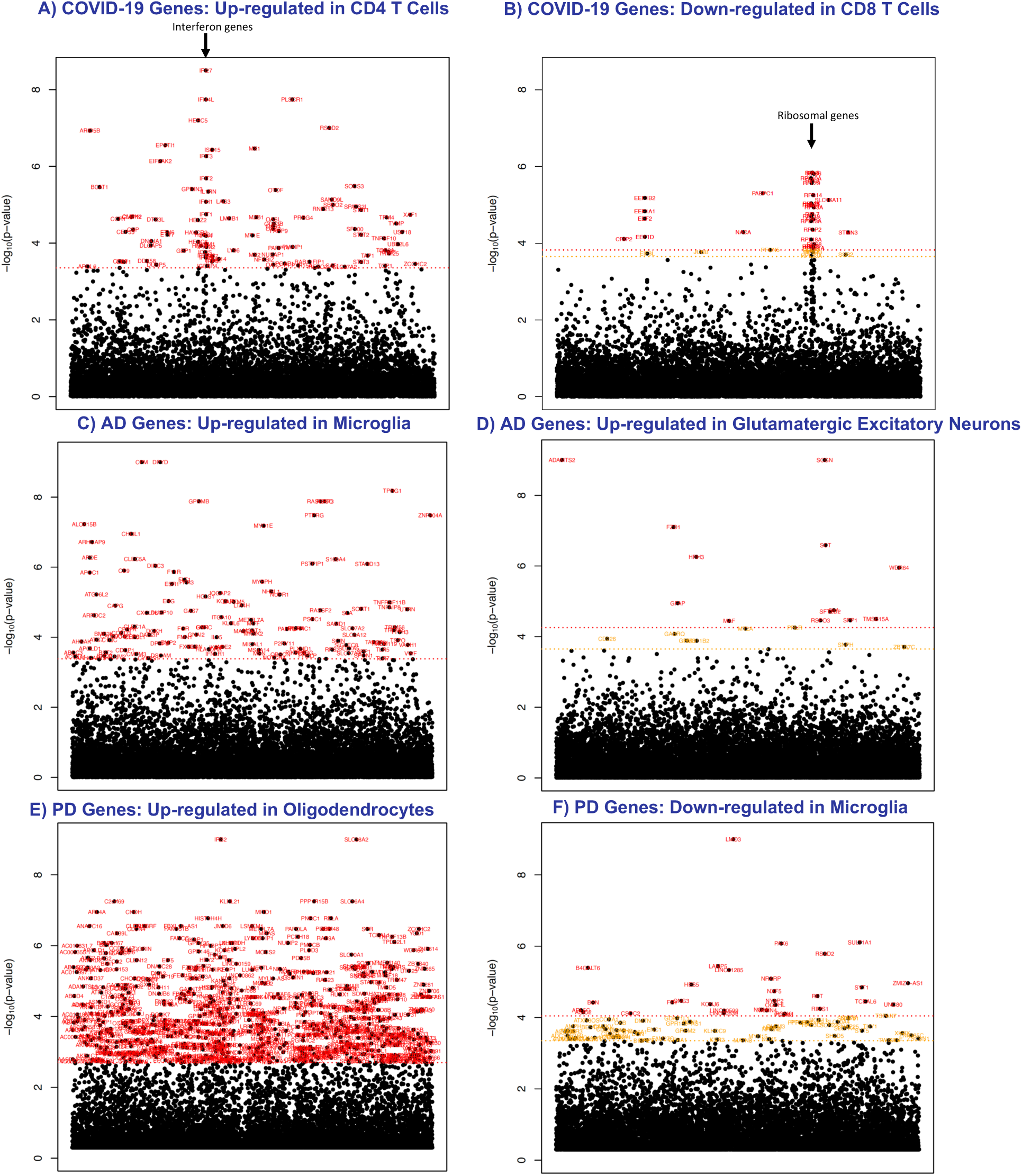
Manhattan plots of differentially expressed genes in AD, COVID-19, and PD. Significance threshold is in red with 0.05 FDR cutoff (Benjamini-Hochberg). In orange is a -log10(p-value) cutoff that maximizes AUC (3.65 for AD, 3.90 for COVID-19, 3.35 for PD; not shown if it is higher than the FDR cutoff red line). The x-axis are genes arranged in alphabetical order. Additional similar plots (including with SCZ) are found in Extended Data Figures 2-7 and Supplementary Figures 3-4. Supplementary Data Files 3-7 show all genes with their p-values.

### Determining factors affecting reproducibility across diseases and datasets

The SumRank approach outperformed other methods in the context of PD and COVID-19, as shown in Table 1 and Supplementary Figure 8. However, the margin of superiority was not as pronounced, likely due to the baseline increased reproducibility of PD and COVID-19 relative to AD. We thus sought to identify the factors underlying the differences in reproducibility between diseases. We restricted all AD datasets such that cases were only those with Braak scores of 5 or 6 and controls were only those with Braak scores of 0-2 to determine if patient selection was a major factor to reproducibility. The AUC with these selection criteria was 0.82, which, though higher than without these criteria, still was much lower than that of PD and COVID-19 (with Braak scores of 3-4, the AUC was 0.69). Given Braak scores are an imperfect measure of disease severity (since some individuals without dementia can have high Braak scores), it is possible that other metrics could decrease patient heterogeneity and increase DEG reproducibility, but alternatively, this might point to a general principle that AD might have more biological heterogeneity than PD and COVID-19, with potentially more factors contributing to the final phenotype clinically diagnosed as AD. Most strikingly, SCZ had a substantially lower reproducibility than all other diseases (Supplementary Note), which could represent substantial heterogeneity in the brains of patient’s with SCZ^4^ due to inherent biology or different life experiences (e.g. more heterogeneous drug/medication use).

We next examined transcriptional effect size to assess its role in reproducibility (Supplementary Figure 9). We found a significant (p=0.0001) positive correlation (Pearson’s r=0.72) between effect size (abs(log_2_(fold-change))) and reproducibility (average AUC for ability to predict case-control status in all datasets) for up-regulated genes, meaning that genes with more differentiation between cases and controls are discovered more regularly across datasets (though for unclear reasons we find no significant relation (r=0.04, p=0.86) for down-regulated genes). Consistent with this, PD and COVID-19, the most reproducible diseases, elicited the strongest transcriptional response, with mean abs(log_2_(fold-change))s of 0.93 (0.97 for up-regulated genes and 0.77 for down-regulated genes) and 0.86 (0.92 for up-regulated genes and 0.39 for down-regulated genes), respectively. In contrast, AD genes had a mean abs(log_2_(fold-change)) of 0.49 (0.55 for up-regulated genes and 0.40 for down-regulated ones) and SCZ genes had a mean abs(log_2_(fold-change)) of 0.25 (0.16 for up-regulated genes and 0.35 for down-regulated ones). We examined the relationship of variance (normalized to effect size by dividing by log2fc) to reproducibility and found a small inverse correlation (r=-0.40; p=0.07) between variance/log2fc and average AUC for up-regulated genes (with down-regulated genes r=-0.03, p=0.89), providing suggestive evidence that reproducibility potentially increases with decreased variance.

We then attempted to identify experimental design factors that increased the performance and reproducibility of DEGs within the same disease. We down-sampled the individuals in the Fujita, MathysCell, and Hoffman datasets to see how varying sample numbers influenced reproducibility measures. We did not discover any clear saturation point, suggesting that reproducibility might continue to increase with even more individuals (Supplementary Figure 10). This is consistent with our observation that for AD datasets there is a positive correlation of Relative Classification Accuracy with sample size (r=0.65, p=0.005; Extended Data Table 1). In contrast, when we down-sampled the Stephenson COVID-19 dataset, reproducibility began to saturate at 70 individuals, and for the other COVID-19 datasets, sample sizes of only 7 cases and controls each had similar reproducibility as those with larger sample sizes (Extended Data Table 4). During this analysis we performed multiple random iterations of the same number of samples and observed that even at 160 samples (80 cases and 80 controls), there was substantial variability in reproducibility, showing the large impact of biological variability to reproducibility (Supplementary Figure 10). We also subsampled all AD datasets with sufficient sample size to 6 cases and 6 controls each and show that reproducibility is highly variable even at the same sample number (Supplementary Table 18). We then down-sampled the cell numbers of the AD datasets to assess its effect on reproducibility and found that reproducibility began to saturate around 0.05 to 0.1 (Supplementary Figure 11). This suggests that particularly when doing analyses involving pseudo-bulking of broader cell types, single-cell experiments should generally prioritize sequencing more individuals rather than more cells per individual.

In addition to sample size, we noted that different studies used different phenotyping criteria to categorize diseased and control individuals. For example, the Hoffman study^26^ carefully selected AD individuals as those fulfilling a combination of neuropathological and clinical criteria. In contrast, the Fujita and MathysCell studies^48,49^ intentionally encompassed a broader range of intermediate phenotypes amongst their cases, likely reducing DEG detection power even with increased sample number. As a result, we found that the Hoffman dataset displayed the highest AUC of all individual AD datasets, driven not only by a large number of individuals, but also likely by the pronounced phenotypic contrasts that separate cases and controls.

We down-sampled AD datasets starting from either the most or least reproducible and found that adding datasets with even low reproducibility continues to increase or maintain the same overall reproducibility of the meta-analysis DEGs, and even down to 3 datasets, the reproducibility of the meta-analysis DEGs are higher than those of the individual datasets (Supplementary Tables 19-20) and higher than the reproducibility of the 3 SCZ datasets. Consistent with this, when we only analyzed the 11 AD datasets with at least 10 cases each the meta-analysis DEGs were not more reproducible than when all 17 datasets were analyzed (Supplementary Table 13). We performed a linear regression analysis of Braak Score on gene expression (while regressing out relevant covariates) to determine if reproducibility would improve with consideration of disease severity. Unfortunately, this did not improve reproducibility (Supplementary Table 16), potentially due to Braak scores being an imperfect correlate of disease severity. We used Gene Set Enrichment Analysis (GSEA)^50^, a threshold-free method to look at pathway enrichment that is generally more robust to power differences, and looked at overlap of pathways. We found that there was a substantial decline in pathway overlap when down-sampling datasets (Supplementary Tables 19-20) or individuals (Supplementary Figure 10), demonstrating that power differences only account for some of the reproducibility differences, while biological variability (captured by the increased number of datasets and individuals) also is important. We also looked at gene set consistency at different thresholds by analyzing the top 25 genes from SumRank on all datasets and assessed the ranks of these genes after down-sampling datasets or individuals (Supplementary Figure 12). We found that these genes were usually in the top 500 genes, though their average rank decreased as datasets or individuals decreased, demonstrating how both power and biological variability impact reproducibility. We also performed SumRank after mapping AD datasets to an AD-only reference (prefrontal cortex of AD patients from Mathys *et al*., Nature 2024^51^) to assess for possible AD-specific cell types/states. We found a slight decrease in reproducibility when only using the broader cell types (AUC=0.76) as well as when using the more specific cell types (AUC=0.65), which could be due to loss of cell type identity in AD^52^, leading to worse cell mapping. However, the DEGs were mostly similar and demonstrate some potential sub-type specific AD DEGs (Supplementary Data File 8).

### DEGs found in meta-analyses reveal known and novel biology

We explored the biological pathways associated with the genes identified in our meta-analyses, initially utilizing gene ontology (GO) via ClusterProfiler^53^. In the context of COVID-19, there was an up-regulation of many interferon genes in CD4 and CD8 T cells, dendritic cells, monocytes, and natural killer cells (Figure 4 and Extended Data Figure 6). This was mirrored in the GO pathways which highlighted processes like "response to virus", interferon response, and other related biological pathways (Supplementary Data File 9). We used gene sets generated from a new stimulation-based Perturb-seq experiment that provided more specific pathways than those generated by gene ontologies^54^ and found that the interferon-beta pathway in particular was up-regulated in COVID-19 cell types more than the interferon-gamma, TNF-alpha, or TGF-beta1 pathways (Supplementary Data File 10). Natural killer cells displayed up-regulated pathways linked to nuclear division and chromosome segregation, stemming from the activation of cell cycle genes during cell proliferation (Extended Data Figure 6; Supplementary Data File 9). B cells showcased elevated endoplasmic reticulum, protein folding, and protein modification pathways, which can be tied to the antibody production process. Across other cell types, there was a noticeable down-regulation of many ribosomal genes, captured under the "cytoplasmic translation" pathway, potentially as a measure to thwart viral RNA translation (Extended Data Figure 7).

For PD, the biological pathways up-regulated were protein localization to the nucleus or mitochondria in oligodendrocytes and oligodendrocyte precursor cells (OPCs) and chaperone mediated protein folding in oligodendrocytes, OPCs, endothelial cells, and astrocytes (Supplementary Data File 9; Extended Data Figures 4-5), particularly due to upregulation of chaperonin genes, including CCT3 and MAPT, which also harbor GWAS variants in PD^55^. This is consistent with the known mechanism of Parkinson’s disease as the misfolding of alpha-synuclein, leading to aggregation of Lewy bodies and the subsequent destruction of dopaminergic neurons^56^. Interestingly, a top down-regulated microglia gene is PAK6 (Figure 4), a PD therapeutics target due to its role phosphorylating LRRK2, a gene mutated in sporadic and inherited PD that activates substantia nigra microglia triggering dopaminergic neuron death^57^.

For HD, we find up-regulation of protein hydroxylation (*PLOD1*, *PLOD2*, *PLOD3*, *P4HA1*, and *P4HA2* genes) in ciliary ependymal cells, which is necessary for CSF transport. Interestingly, we find dramatic down-regulation of synapse organization and neuron projection development genes in FOXP2 neurons (Supplementary Figure 5E and Supplementary Data File 9), pointing to their dysfunction. This is consistent with reports that FOXP2 is degraded in HD due to its interaction with the mutant huntingtin protein, diminishing FOXP2 neuron function^58^. Indeed, FOXP2 overexpression rescues HD behaviors in mice^59^, while downregulation of FOXP2 worsens HD symptoms^60^.

For AD microglia, cytokine production and immune response pathways were up-regulated, likely promoting neurodegeneration and AD progression^61^. In endothelial cells, negative regulation of growth was up-regulated, and in astrocytes amino acid catabolism was downregulated (Supplementary Data File 9). The pathways, however, were not consistent and were mixed with many other pathways of unclear relevance. Given this, we used STRING^62^ to construct protein-protein interaction (PPI) networks using AD DEGs, searching for dense sub-networks that could improve our power to detect relevant biological pathways (Supplementary Data Files 3-4 and 6). This yielded more clear signals. For example, the most densely connected network of up-regulated microglia genes was enriched for pathways in regulation of lipid transport, driven by genes such as APOE, TREM2, and SORT1, which are all AD GWAS genes^63–65^ (Supplementary Figure 13). In GABAergic neurons, chemokine driven cellular response, cell fate specification, and wide pore channel activity (driven by the connexins GJA3, GJA4, GJB2, and GJB4) were enriched in the most connected up-regulated network. For the most connected down-regulated networks, signaling by neurotrophic tyrosine kinase receptors, such as BDNF, DUSP4, DUSP6, and VGF, was enriched in glutamatergic neurons, while in astrocytes glutamate metabolism was enriched, together showing how AD inhibits glutamatergic excitatory neuron growth and function. When we applied this approach to PD, oligodendrocytes, astrocytes, and OPCs show up-regulation of networks enriched for chaperone-mediated protein folding and protein localization to mitochondria (Supplementary Figure 14), while PD microglia show up-regulation of lipid transport, revealing a common mechanism with AD microglia. Similarly, HD mural cells showed clear up-regulation of chaperone folding genes as the most densely connected network, showing a possible common mechanism with PD glia.

In addition to these pathways, SumRank pointed to many genes with very clear reproducibility across a large majority of datasets that had not previously been highlighted by other AD papers in a cell type specific manner. For example, *PDE10A* was down-regulated in excitatory and inhibitory neurons (Supplementary Data File 3). PDE inhibitors have long been proposed for AD^66^, and PDE10A inhibitors have shown some improvement in AD symptoms^67^. We also observed downregulation of *HES4* in inhibitory and excitatory neurons, *HES5* in OPCs, *VGF* in inhibitory and excitatory neurons, and microglia, and *VEGFA* in OPCs, all of which are involved in neuron^68–70^ and endothelial growth^71^. Similarly, *SPP1*, a gene associated with synapse loss^72^, was up-regulated in endothelial cells and glutamatergic neurons, while *ADAMTS2*, a gene that breaks down extracellular matrix in the brain^73^, was up-regulated in glutamatergic neurons. Together, this suggests that AD pathophysiology might involve inhibition of growth pathways, which could decrease synaptic plasticity and contribute to the cognitive dysfunction in AD; thus, therapeutics aimed at increasing these factors might be useful^56^. The importance of G protein mediated signaling and amino acid and nucleotide metabolism dysregulation in AD was demonstrated by the fact that *RASGRP3* and *DPYD* were up-regulated in microglia and *SLC38A2* was upregulated in oligodendrocytes, while *ARRDC3* was down-regulated in astrocytes and *BCAT1* was down-regulated in oligodendrocytes. Lastly, we observed that the *CAT* gene was down-regulated specifically in glutamatergic neurons (in the SumRank analyses but not in the merge or inverse variance analyses; Figure 2E). Catalase activity had previously been shown to be decreased in AD due to amyloid-beta^74^, and a catalase derivative has been proposed as a possible therapeutic for AD to decrease oxidative stress from free radicals^75^. These analyses suggest that *CAT* is specifically down-regulated in glutamatergic neurons and not GABAergic inhibitory neurons or other cell types, consistent with the observation that excitatory neurons have increased oxidative stress and die at higher rates in AD.

Our approach of focusing on reproducible genes and predicting phenotypes in leave one out analyses provides some internal validation for our genes, but we wanted to compare to an independent system of AD. We thus performed experimental validation of one of the SumRank DEGs using the 5xFAD mouse line, which is a well-known model of late-onset AD^76^ that overexpresses a mutant human amyloid-beta precursor protein, harbors multiple AD-associated mutations in human presenilin 1, and has been shown to have many phenotypic similarities to humans with AD, including amyloidosis and behavioral impairment. We looked to test a gene that was significant in the SumRank but not merge or inverse variance methods and that had potential therapeutic relevance but with no prior known cell type-specific data. We thus chose the *BCAT1* gene, which we found only by SumRank (not merge or inverse variance) to be down-regulated in AD oligodendrocytes and is a cytosolic amino acid transaminase in both humans and mice. We performed multiplexed immunohistochemistry (IHC) staining on slices of the medial prefrontal cortex for BCAT1 and measured the degree of staining in CC1 SOX10 double-positive, mature oligodendrocytes. We found that the 5xFAD mice had significantly lower *BCAT1* expression in oligodendrocytes (Figure 5), demonstrating for the first time in both humans and mice that *BCAT1* has oligodendrocyte-specific decreased expression in AD. BCAT1 facilitates transamination of branched chain amino acids (BCAAs) to produce glutamate and GABA, which are essential for cognition, while elevated BCAAs can lead to neurotoxicity^77^; thus, decreased *BCAT1* expression could lead to AD progression through elevated BCAAs or decreased glutamate. Our findings also point to oligodendrocyte-specific manipulation of BCAA metabolism as a potential therapeutic for AD^78^. Notably, we also find BCAT1 specifically down-regulated in oligodendrocytes in HD.

**Figure 5.**
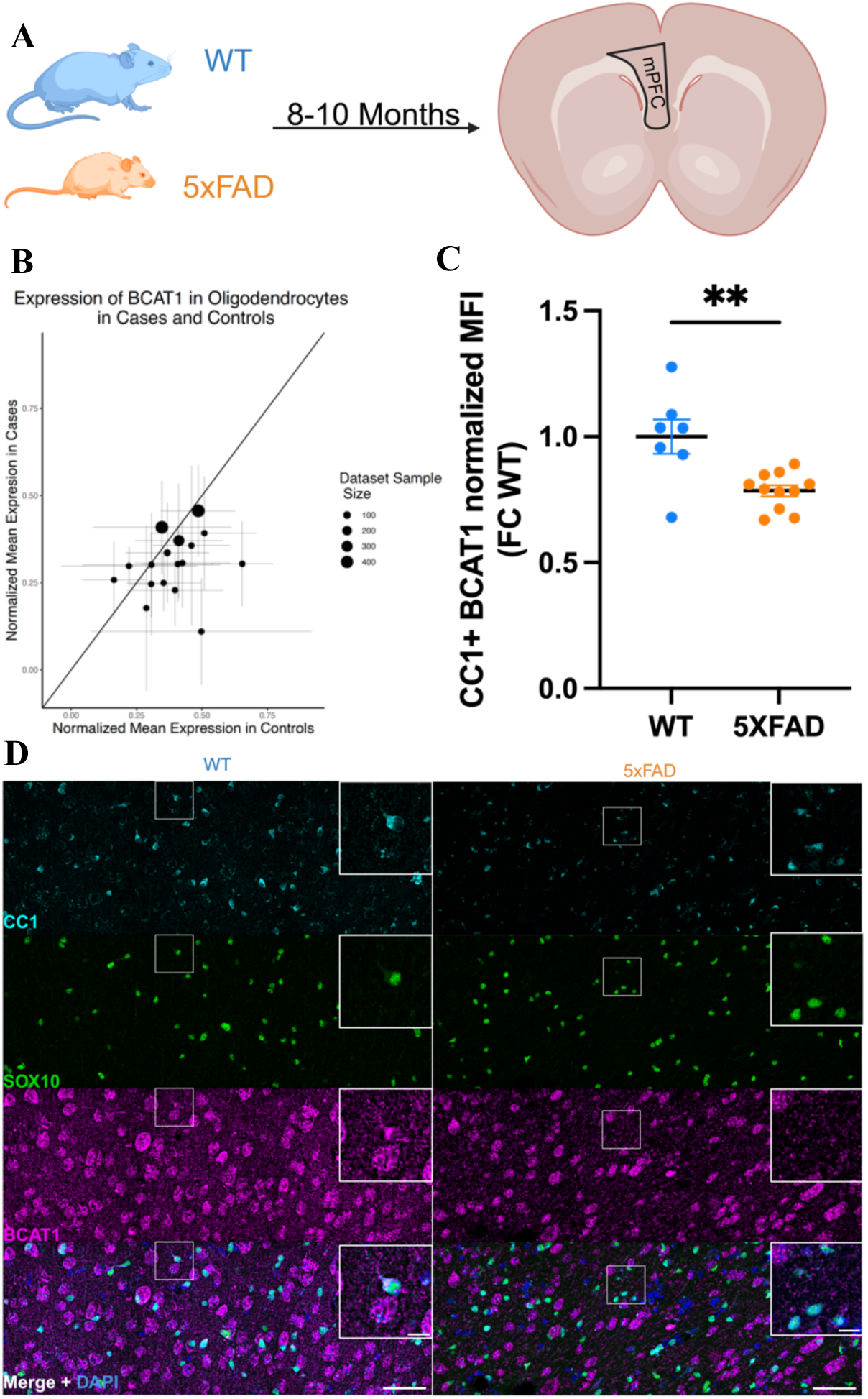
Experimental validation of a meta-Analysis AD DEG in a mouse model. **A)** Schematic of experiment measuring cell type-specific expression in the medial prefrontal cortex of 5xFAD mice from 8-10 months old. **B)** Expression of B*CAT1* in oligodendrocytes in human postmortem snRNA-seq datasets. Values above the line (intercept=0, slope=1) are up-regulated, while values below the line are down-regulated. Error bars are standard deviations in all plots. **C)** Protein expression of *BCAT1* in oligodendrocytes of 5xFAD and WT mice obtained by quantifying the mean fluorescent intensity (MFI) expressed as fold change (FC) over WT animals (n=7 WT, 11 5xFAD mice). Data represented as mean ± s.e.m. Results are significant at p=0.0026 (students two-tailed unpaired t-test). **D)** Representative multiplexed immunohistochemistry (IHC) staining of cortical slices from a 5xFAD mouse of *BCAT1* and 2 oligodendrocyte specific markers (*SOX10* and *CC1*) along with the merged image (Scale bar=50µM large images, 10 µM insets).

We compared our meta-analysis DEGs to those exhibiting differential snATAC-seq peaks, given differential expression should be correlated with differential chromatin accessibility. We used differentially accessible peak data from a study performing snATAC-seq on prefrontal cortex of 44 AD cases and 48 controls^52^ and found the genes whose promoters were nearest to the peaks. We found that the SumRank DEGs had significant (p<2.2e-16) enrichment for the top genes from differential accessibility analysis and, remarkably, even more enrichment than the DEGs from RNA of the same individuals from whom the snATAC-seq data were derived (Supplementary Figure 15), providing an additional mode of validation. We then assessed the intersection of the 708 unique AD DEGs at the 3.65 -log10(p-value) cutoff with genes found in the largest AD GWASs^63–65^ and found 9 unique genes out of the 105 genes in GWAS to be shared (Supplementary Table 21; p=1.3e-4, Fisher’s exact test). When we looked at the intersection with AD whole-exome studies^79–81^, 4 of the 28 genes were shared (p=1.1e-4, Fisher’s exact test). Of the 1187 unique PD DEGs at the 3.35 -log10(p-value) cutoff, there were 6 unique genes out of the 72 genes in PD GWAS^55^ shared (p=2.0e-05). Despite this indicating a statistically significant enrichment, it still represents a relatively minor overlap, suggesting that the genetic variants underlying predisposition to AD are often not the same as the genes whose expression are altered downstream of individuals with multiple years of AD. We also evaluated the overlap of AD, HD, and PD genes and found 56 shared up-regulated genes amongst all 3 diseases (6 found in 2 cell types) and many more up- and down-regulated genes shared between pairs of the diseases (Supplementary Data File 9). It is likely that these shared genes represent a common neurodegenerative biological pathway, but no significant GO enrichment was found.

### Adaptation of non-parametric meta-analysis method uncovers sex-specific DEGs

The female sex-bias in AD^82^ motivated us to search for genes with sex-specific expression such that they were only up-regulated in one of the sexes. We performed two types of analyses to assess for sex-specific expression (Figure 6; Methods). In our first analysis we used DESeq2’s interaction term (SEX:Diagnosis) to look for genes with significant interaction between Sex and Diagnosis within each dataset. We then fed these values into the SumRank method, adding up the p-value ranks of the genes across each dataset, considering only the top 65% of datasets (to be consistent with the general analyses), and using permutations (permuting sex) to calibrate the p-values. This analysis will find all genes with significant differences in case vs. control gene expression between the sexes, but it could also find genes with decreased expression in one sex and unchanged expression in the other sex.

**Figure 6.**
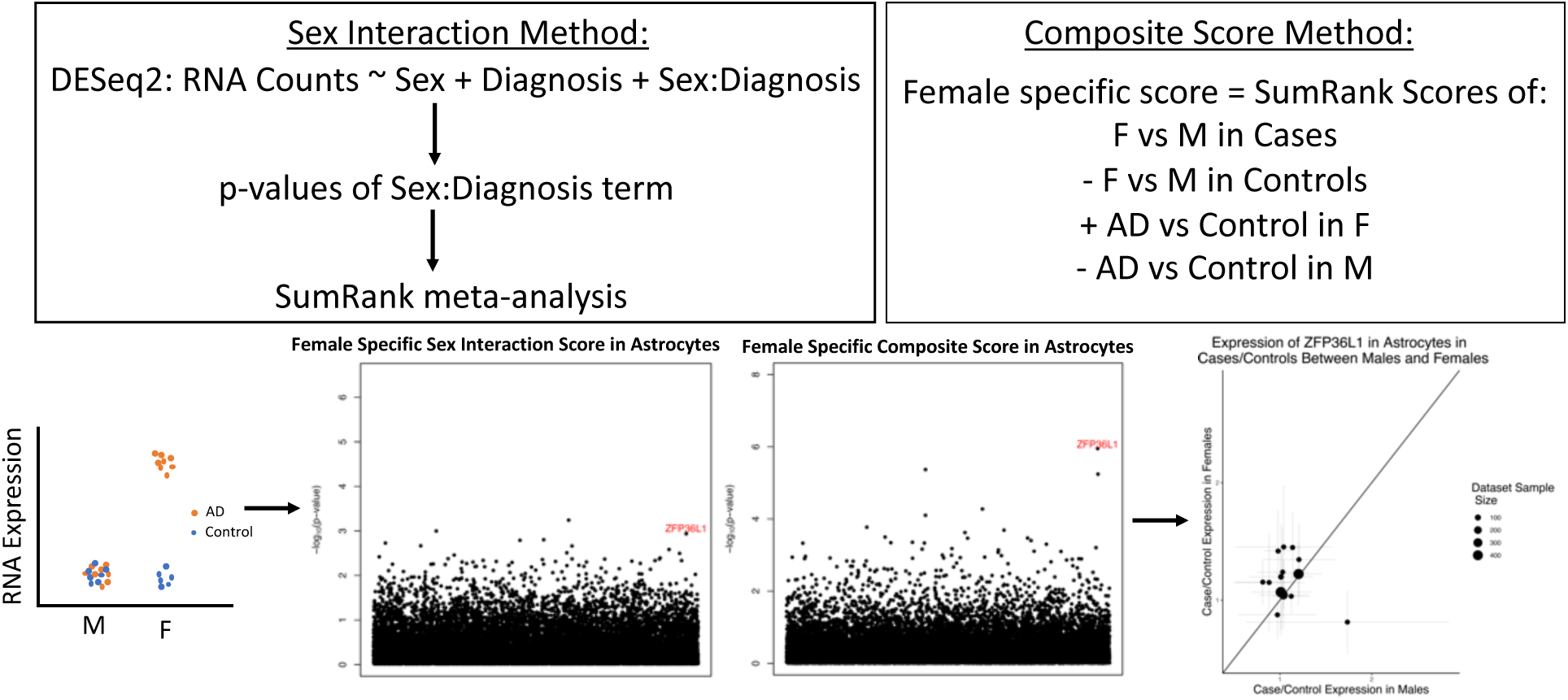
Schematic of the two methods used for assessing sex-specific expressed genes. The Sex Interaction method uses the SumRank meta-analysis on the p-values of the Sex:Diagnosis term from DESeq2, while the Composite Score method takes the composite of 4 different SumRank scores (shown here for female specific scores; the male specific score is defined analogously). On the bottom left is a schematic of an example female-specific expressed gene. The Manhattan plots highlight the *ZFP36L1* gene. The ratios of mean expression of cases over mean expression of controls of *ZFP36L1* in females (y-axis) and males (x-axis) are plotted in the bottom right. Values above the line (intercept=0, slope=1) are up-regulated in females more than males, while values below the line are up-regulated in males more than females. Error bars are standard deviations. Plots of the expression of *ZFP36L1* in individuals within each dataset are in Supplementary Figure 16.

In order to focus on genes that have up-regulated expression in one sex but are unchanged in the other, we devised another method that works by summing up four different scores to create a composite score. We performed differential expression and SumRank meta-analyses in DESeq2 to obtain p-values for scores between males and females in only cases and in only controls as well as cases vs controls in only males and in only females. Female specific scores were calculated as the sum of the -log10(p-values) of the cases vs. controls in females with the -log10(p-values) of the females vs. males in cases subtracted by the -log10(p-values) of the cases vs. controls in males and the -log10(p-values) of the females vs. males in controls. Male specific scores were calculated analogously, and we calibrated all p-values empirically with permutations.

At q-value or Benjamini-Hochberg based FDR cutoffs of 0.05 no genes were significant with both methods, so we loosened our thresholds. We looked for genes that had -log10(p-values) above 3.65 (the threshold chosen for the general analyses) in the Composite Score approach and were in the top 15 genes (0.1%) in the Sex Interaction approach. This led to the discovery of several female-specific genes, *SLITRK5* in OPCs, *ZFP36L1* and *DUSP1* in astrocytes, *DAPK2*, *APOE*, and *OR4N2* in GABA inhibitory neurons, and two male-specific genes, *MYC and IL16* in glutamatergic neurons (Figure 6, Supplementary Figure 16, Supplementary Data File 11). Of these only *ZFP36L1* and *SLITRK5* were significant in the composite method at an FDR 0.05 cutoff. *ZFP36L1* is a 3’UTR binding protein that influences transcriptional regulation and has been found to be a differentially expressed gene that is a candidate biomarker for AD^83–85^. Interestingly, the APOE risk factor is known have a stronger association with females relative to males^86^. We also applied this method to COVID-19 and found *CLU* in dendritic cells and monocytes, *MT1E* in other_T cells and *G0S2* in CD4 T cells as male-specific expressed and *CAMK1* in dendritic cells as female-specific expressed (Supplementary Data File 11).

The lack of clearly significant genes in any of the SumRank sex-specific analyses is likely due to insufficient power, because these analyses require at least twice as many individuals as the case-control analyses given the extra consideration of sex. In addition, it is also probable that the sex-specific effect sizes are much smaller than the effect sizes differentiating cases vs. controls more generally, so overall these results underscore the need for more data to better delineate these effects. We note that when we used the merge method with DESeq2 sex interaction, we found several genes that were significant at Bonferroni corrected p-value thresholds of 0.05, but these genes were not significant and ranked extremely low in the SumRank methods due to only being significant in one or a few datasets (Supplementary Figure 17), showing again the importance of reproducibility in these analyses (nevertheless, *CLU*, *G0S2 MT1E,* and *CAMK1* all had q-value FDR<0.1 in their respective cell types for the merge sex interaction method).

## Discussion and Conclusion

Here we assessed the reproducibility of DEGs across many AD, PD, SCZ and COVID-19 datasets. We find that DEGs from single AD and SCZ datasets generally have poor reproducibility and thus cannot predict case control status in other AD or SCZ datasets, though predictive power is improved with increased numbers of individuals in the study. In contrast, even small individual PD and COVID-19 studies have moderate predictive power for case control status in other datasets. This study provides strong evidence that for diseases of high heterogeneity like AD and SCZ, the DEGs of case-control datasets of relatively small sample sizes (fewer than 100 total individuals), even when derived in a statistically rigorous manner, have a low likelihood of being reproduced in many other datasets and thus are more likely to be dataset specific artifacts rather than reliable indicators of disease pathology. In contrast, acute diseases or those with more uniform responses, such as PD, HD, and COVID-19, produce DEGs with moderate reproducibility across studies.

This presents a paradox in that for diseases with heterogeneous gene expression and low reproducibility, likely including many neuropsychiatric diseases, it is *more* important to ensure that genes are found reproducibly across multiple studies to avoid false positives. Motivated by this, we provide here a path towards GWAS level of reproducibility through the development of a novel meta-analysis method (SumRank) that prioritizes reproducibility across datasets. We show that SumRank outperforms merging of datasets with batch correction (the standard scRNA-seq method) and combining effect sizes with inverse variance weighting (the standard GWAS method). The DEGs found by SumRank have improved specificity as measured by ability to predict case-control status in left out datasets and demonstrate that many previously highlighted genes thought to be differentially expressed in AD do not show differential expression across many datasets. The inverse variance method, though successfully utilized in GWAS, performs poorly for meta-analysis of scRNA-seq data due to dataset specific artifacts that are carried through, such that some genes with very low p-values in a small number of datasets are considered significant even though they are not differentially expressed in most datasets. This effect is much more pronounced in single cell studies relative to GWAS due to the lower stability of RNA expression relative to DNA, leading to greater propensity for very poorly calibrated p-values. The merge method generally works much better than the inverse variance method (likely due to DESeq2’s ability to have a dataset covariate correction) but still performs more poorly than the SumRank method for the same carried over artifact issue. Moreover, the merge method is much slower than the other methods as the merge process can take several hours, particularly for the large datasets.

With the SumRank method, we were able to discover previously known and novel COVID-19 biology, such as division of NK cells and down-regulation of ribosomal genes. For PD, we consistently found up-regulation of chaperone-mediated protein folding and protein localization to the nucleus and mitochondria across multiple cell types, with GWAS genes driving the signal. For HD, we find a dramatic down-regulation of synaptic function in FoxP2 neurons. We also use PPI networks to increase biological pathway clarity, demonstrating up-regulation of lipid transport AD and PD microglia and down-regulation of glutamate functioning via astrocytes and glutamatergic neurons, driven by key GWAS genes and consistent with known dysfunction of glutamatergic neurons in AD. Users can employ similar tools to improve biological insight from SumRank DEGs. We validate the *BCAT1* gene as down-regulated in oligodendrocytes of human and mice with AD.

A particularly interesting finding was that SCZ has the lowest reproducibility of the studied diseases, which could be due to its relative increased biological complexity. SumRank only slightly increases reproducibility for SCZ (mean AUC=0.55 for individual datasets and 0.62 for SumRank). We believe highly complex diseases like SCZ will likely require either sub-phenotyping to find more biologically homogeneous cohorts or much larger sample sizes and/or more datasets to achieve clear reproducible DEGs, similar to GWAS for SCZ and autism, despite the high heritability of these diseases^87^. AD has lower reproducibility than PD, HD, and COVID-19 but still has hundreds of DEGs. Nevertheless, the biological inference was less clear due to the large number of pathways discovered with less support for each one. This could indicate true biological heterogeneity (i.e. relatively more distinct pathways all contributing to this disease), which would decrease power for pathway inference. Analyzing reproducibility of networks using approaches such as WGCNA^88^ could yield increased power. Another avenue would be to find reproducible gene regulatory networks created by ATAC-seq, ChIP-Seq, protein-protein interaction, metabolomic, or proteomic^89^ data and assess their overlap with RNA expression to find reproducible biological pathways. We also note that for our analyses we pseudobulk at both broad cluster levels as well as more narrow neuron levels. Our choice of cluster resolution will inevitably obscure heterogeneity within the clusters, while there will cell type-specific DEGs at other cluster resolutions, some of which might be relevant for the disease. SumRank is a generic method that can be used at any cell type resolution, so additional studies can be performed to assess biological pathways that may be found in potentially more rare cell types or subpopulations.

Single-cell transcriptomic case-control studies have, to date, involved limited numbers of individuals for studies outside of AD and COVID-19, and for many neuropsychiatric disorders it likely will take many years to reach the same cohort sizes and number of studies as in AD and COVID-19. It is thus critical to apply the lessons learned from AD, PD, HD, COVID-19, and SCZ to diseases with increasing numbers of individuals sequenced. Our results suggest that when designing scRNA-seq case-control studies, it is more important to sequence a larger number of individuals rather than more cells once there are over ∼40 cells per cell type of interest (when pseudo-bulking). Investigators could also consider looking at extremes of phenotypes to increase power. Most importantly, it is critical for all studies, particularly small ones (fewer than 50 cases and controls each, based on observations from this study), to demonstrate clear reproducibility in the DEGs discovered and show that (ideally for each individual gene) this reproducibility exceeds the reproducibility expected by chance.

We lastly highlight limitations of the SumRank method and single-cell meta-analysis methods in general, which will be important to overcome in the future to produce GWAS-quality meta-analyses. For the SumRank method in particular, a substantial limitation is the lack of weighting, which can cause substantial power limitations. We were not able to come up with a reliable method for weighting the studies, because, for example, although there was a general correlation of predictability of DEGs (AUC) with number of individuals, the relationship was not uniform as some larger studies had poorer predictive power for reasons such as more heterogeneous phenotyping or poorer sequencing quality (e.g. multi-ome data in the Su COVID-19 dataset), so weighting by number of individuals, number of cells, or sequencing depth could lead to substantial biases. SumRank also requires multiple datasets of the same tissue region at the same context, or the method will usually lose power and fail to find context-specific signals. One possible way to find these signals is to first assess for more homogeneous clusters of individuals or datasets, apply SumRank to these subsets of the data, and then determine the characteristics of the individuals/datasets leading to the unique signals. The primary computational burden of SumRank is performing multiple rounds of differential expression after permutations. For 4 COVID-19 datasets on one cell type (Monocytes), the SumRank command itself takes only 36.8 minutes to run 1000 times on a standard laptop, while additional datasets add less than linear time (42.2 minutes for 8 COVID-19 datasets), while the merge process takes 44.4 and 119.3 minutes for 4 and 8 datasets, respectively, showing non-linear scaling, and the inverse variance process takes 111.7 and 149.7 minutes. However, running DESeq2 on 1,000 differential expression permutations of the same datasets takes 9.3 hours (if not parallelized), which will be required for all methods if p-values are obtained empirically. It is currently unclear how to speed up the differential expression process, but this study used clusters with parallelization to ensure reasonable analysis time. SumRank was used in this study with a discrete cluster-based analysis of cases and controls with separate analysis of cell types, though in future work it can potentially be adapted to analyze cell and phenotype heterogeneity on a continuous spectrum and with cellular communities rather than discrete cell types. Lastly, a SumRank-specific limitation is optimization of choice of number of datasets to use, which can increase computational time.

Other limitations are generic to all single-cell meta-analysis methods. For example, there is currently no method to account for possible relatedness amongst the individuals either within or across datasets, unlike GWAS meta-analyses, which are now able to condition out relatedness without fully removing related individuals^90^. Accounting for relatedness is likely more difficult for RNA and other modalities relative to DNA, but future meta-analyses could potentially account for this by either having genotyping of all patients or looking for increased correlation in expression above the background. Similarly, population structure (e.g. individuals of a certain ethnic background being enriched in cases) could lead to spurious associations and must be accounted for in future analyses.

Refinement of GWAS methodologies, including addressing many of these issues, took over a decade^91^. Meta-analyses of single cell data face many challenges beyond those of genetic data, such as a greater propensity for dataset specific artifacts (due to the relative instability of RNA and potential for gene expression changes during technical processes), expression differences across tissues and tissue regions (increasing the noise when combining datasets), differences in life environments between cases and controls (e.g. medication use), and less clear principles for how genetic relatedness affects gene expression between individuals. On the other hand, the average effect sizes of RNA are usually much higher than genetic effect sizes, which are brought down due to natural selection, as evidenced by the mean effect size of individual DEGs for AD in this study being 1.40 relative to 1.05 for AD GWAS^63^. This means it is likely that lower sample sizes will be required for single cell case control analyses relative to GWAS. Nevertheless, it will be important to apply any applicable lessons from GWAS to single cell case control analyses, including the applications of GWAS results. For example, once there are an adequate number of studies of other neuropsychiatric traits, we believe the SumRank method can be adapted to perform cross-disorder analyses, which will aid in revealing shared biology between disorders, similar to cross-trait GWAS analyses^92^. Overall, this study is intended to take a strong step in bringing single cell case control studies to GWAS levels of reproducibility, which we hope will clarify the cell type-specific biological changes involved in different conditions, ultimately leading to more reliable drug targets to reverse disease pathophysiology^93^.

## Acknowledgments

We thank all members of the Satija lab and members of the CEGS Center for Integrated Cellular Analysis in New York city for helpful discussions and constructive criticism. We thank Li-Huei Tsai and Ravikiran Raju for providing data from Barker *et al.*, 2021. We thank Brad Ruzicka and Shahin Mohammadi for providing their code for the analyses of Ruzicka *et al.* 2024 and answering questions about their publication. This work was supported by the Chan Zuckerberg Initiative (EOSS-0000000082 and HCA-A-1704-01895 to R.S.) and the NIH (RM1HG011014-02, 1OT2OD026673-01, DP2HG009623-01, R01HD096770 and R35NS097404 to R.S., NIH-NINDS R01NS122316 and R21NS121786 to E.K., and NIH-NIMH T32MH019524 to D.A.).

## Data availability

All data are publicly available online (see Supplementary Data File 1 and Methods for details).

## Code availability

Code for all new analyses in this paper, including runnable software for the SumRank method, are available in a Github repository: https://github.com/nathan-nakatsuka/scRNA_Reproducibility.

## Online Methods

### Datasets

Count matrices were downloaded from GEO for GSE129308 (Otero-Garcia *et al.*^94^), GSE147528 (Leng *et al.*^95^), GSE140511 (Zhou *et al.*^96^), GSE138852 (Grubman *et al.*^97^), GSE174367 (Morabito *et al.*^98^), GSE157927 (Lau *et al.*^99^), GSE163577 (Yang *et al.*^100^), GSE183068 (Sayed *et al.*^101^), GSE148822 (Gerrits *et al.*^102^), GSE160936 (Smith *et al.*^103^), GSE167494 (Sadick *et al.*^104^), GSE157783 (Smajic *et al.*^10^), GSE184950 (Wang *et al.*^15^), GSE193688 (Adams *et al.*^14^), GSE243639 (Martirosyan *et al.*^12^), GSE148434 (Lee *et al.*^13^), GSE173731 (Garcia *et al.*^105^), GSE180928 (Lim *et al.*^23^), and GSE152058 (Matsushima *et al.*^21^). Other matrices were downloaded from Synapse (Mathys *et al.*, 2019^45^, Mathys *et al*., 2023^49^, Hoffman *et al.*^26^, Fujita *et al.*^24^, Ruzicka *et al*.^4^), CellxGene (Gabitto *et al.*^46^), Zenodo (Batiuk *et al*. 2022^5^: https://zenodo.org/records/6921620), NEMO (Ling *et al*.^6^ and Handsaker *et al.*^106^), the Broad Institute Single Cell Portal (SCP1768: Kamath *et al.*^11^), or from the authors directly (Barker *et al.*^107^). Relevant meta-data were also retrieved from the corresponding publications. COVID-19 datasets were obtained from Tian *et al.*^108^.

### Quality Control and Data Processing

Count matrices were first converted to Seurat objects using the Seurat V4 pipeline. Mitochondrial percentage, nCount_RNA, and nFeature_RNA were assessed for each dataset, and cells with outlier values were removed from the dataset (Supplementary Data File 1). Subsequently, SCTransform v2 was performed for normalization and variance stabilization of the data, then PCA was run with 30 PCs maintained, and UMAP was run on the PCA reduced dataset with dims 1:30 selected. Cell types were then determined by mapping to the class and subclass groupings of the Azimuth motor cortex for AD and SCZ datasets and the Azimuth PBMC reference for COVID-19 datasets using FindTransferAnchors in Seurat with 1:30 PCA dimensions, and refDR reduction, with all other settings left at default. Mapping to the Azimuth reference ensures that even if the mapping is not perfect, there likely will be no bias since the mapping quality should be similar for the cases and controls within each dataset. For PD and HD datasets the cells were mapped to the Kamath *et al.*^11^ PD and Matsushima *et al.*^21^ HD datasets, respectively, due to lack of other reliable midbrain and caudate references. For the Lim HD dataset we observed that the FoxP2 cells did not have reliable mapping, so we did not include them in our analyses (see Supplementary Note).

### Differential Expression

Each dataset was pseudobulked by obtaining either the aggregate sum of all counts (for DESeq2 analyses) or the mean value (for all other analyses) for each cell type at the Azimuth class or subclass level for each individual in each dataset. Differential expression was done by comparing cases to controls within each cell type and using multiple different methods. For our general analyses DESeq2^30^ was used to compare cases to controls with logfc.threshold and min.pct set to 0 to ensure that all genes were included (pseudocount.use was set at 1 due to the need for round count numbers for DESeq2). No normalization is needed prior to DESeq2 analyses, because DESeq2 performs internal normalization through its median of ratios method to account for sequencing depth and RNA composition. Mitochondrial genes were removed from all results and the final gene set was chosen as the intersection of all of the datasets for the particular disease leading to 15,201 genes for AD, 11,067 genes for COVID-19, 17,823 genes for PD, 14,833 genes for HD, and 17,420 genes for SCZ. To test down-regulation, differential expression was done between controls relative to cases with the same downstream process repeated as for the up-regulated genes. Violin plots were made in Seurat using the VlnPlot command after subsetting to the cell type and gene of interest. DESeq2 was also used in separate differential expression analyses while regressing out relevant clinical covariates (any of the following if they were present in the dataset’s metadata: sex, age, PMI, RIN, education level, ethnicity, language, age at death, batch, fixation interval, nCount_RNA, and nFeature_RNA) using design=∼Diagnosis+ClinicalCovariate. Differential expression was also done using logistic regression with the “FindMarkers” function in Seurat V4 with test.use=“LR” and latent.vars set to the clinical covariates with all other settings set to default. Linear regression was performed in R, fitting a model of Braak score on gene expression and clinical covariates using the “lm” function in base R with all other settings set to default.

To test the ability of each gene to predict case-control status in each dataset (as a separate analysis from the general differential expression analyses above), we used logistic regression models of case-control status with and without each gene as implemented in the “FindMarkers” function in Seurat V4 with test.use=“LR”, pseudocount.use=0.01, logfc.threshold=0, min.pct=0 (with all other settings at default) and obtained the log2fc and p-values for each gene separately for each cell type and each dataset. We then took the mean of each gene’s abs(log2fc) and signed -log10(p-values) (negative for genes with negative log2fc values) in all datasets to obtain each gene’s average ability to predict case-control status across all datasets (separately for each cell type).

To test the Ruzicka *et al.* differential expression pipeline, we converted the provided ACTIONet rds object into a singlecellexperiment object and separated the dataset into the McLean and MtSinai cohorts. We then created pseudobulk profiles with the mean of log-transformed counts within each individual and cell type. We filtered out the SZ3, SZ15, SZ24, SZ29, and SZ33 individuals and cells with capture rate less than 0.05 as done by Ruzicka *et al*. We then removed effect of batch and HTO variables using the removeBatchEffect function in limma^109^ version 3.46.0, while incorporating age (split in half into older age and younger age), sex, postmortem interval, and the log transform of average number of UMIs per cell. We then used muscat version 1.18.0 to perform differential expression with the limma-trend model using muscat default filtering for genes and min_cells=10 (see Supplementary Note for more details and explanation).

### SumRank Meta-Analysis

The genes of all datasets were ranked by their signed -log10(p-values), with genes having negative log2(fold-change)s being set to negative so that down-regulated genes would be at the bottom and up-regulated genes at the top. The ranks of each gene for each dataset were then normalized by first subtracting one from them and then dividing by one less than the total number of genes (so that the highest ranked gene was 0 and the lowest ranked gene was 1). To improve power, by removing the influence of datasets that might have poor scores for artifactual reasons, only the ranks of the top datasets were considered for each gene. The number of datasets chosen for consideration was based on the ability of its resulting gene set to most accurately predict case-control status in left-out datasets (measured by AUC; see below), with the additional specification that at least half of the datasets be used. AUC (area under the receiver operating characteristic (ROC) curve) is the area under the ROC curve, which plots sensitivity against specificity. We took the sum of the normalized ranks of the top datasets for each gene. If the sum was greater than the number of datasets divided by two, we set the value to the number of datasets divided by two (to ensure that genes that were consistently not differentially expressed would not be considered significant).

The AUC represents the probability that the model, if given a randomly chosen positive and negative example, will rank the positive higher than the negative, with 1.0 being a perfect score and 0.5 being the lowest score. This metric allows us to compare the performance of different models assuming the datasets are roughly balanced between cases and controls. The Irwin-Hall distribution is the theoretical null distribution for the SumRank statistic, because it assumes that the genes in each study are uniformly distributed and each study is independent of the other, and the Irwin-Hall distribution is the sum of independent, uniformly distributed random variables. We thus initially obtain p-values for each gene using an Irwin Hall distribution (two-sided) as implemented in the unifed version 1.1.6^110^ package, dirwin.hall function, with number of datasets as the number of uniform distributions specified. However, it is possible that genes are not uniformly distributed given the complexities of gene expression, and we also choose only a subset of datasets for each gene, so for both of these reasons the distribution will deviate from Irwin-Hall. We thus calibrated the p-values by permutations (see below).

### Merge Meta-Analysis

After quality control, the Seurat objects for each dataset were first subsetted to the relevant cell type and then merged using the Seurat merge function with all settings at default. The count matrices for the merged objects had 1 added to them (for a pseudocount) and were then converted to DESeq data set types with the DESeqDataSetFromMatrix command with design = ∼Diagnosis+Dataset, to provide some accounting for dataset specific batch effects (this design regresses out dataset specific artifacts). DESeq2 differential expression was then performed, and results were extracted for the Diagnosis variable (p-values and log2 fold-changes for each gene).

### Inverse Variance Meta-Analysis

Differential expression effect sizes (log2 fold-change) and standard errors for each gene and each dataset were obtained from the DESeq2 output as described in the Differential Expression section above (with design = ∼Diagnosis and no other variables regressed out). These summary statistics were then put into the metagen function from the meta version 6.5.0 R package^42^ to obtain combined effect sizes across the datasets with sm = “OR” (to specify odds ratio was used), fixed=FALSE, random=TRUE (to specify using a random effects model, given the expected heterogeneity in the datasets), method.tau=“REML” (restricted maximum likelihood method to obtain the estimator from inverse variance weighting), hakn=TRUE (Hartung and Knapp statistic adjustment), control=list(stepadj=0.1,maxiter=10000). These parameters are recommended to minimize the risk of false positives^42^. The effect sizes were obtained from TE.random and the p-values obtained from pval.random (two-sided). When we attempted to improve the inverse variance method by only taking a certain percentage of top datasets, we found that this did not increase the AUC, so we retained all datasets for this analysis.

### Permutations for obtaining empirical p-values

To calibrate p-values for case-control differential expression, permutations of case and control status were performed either 1,000 or 10,000 times by sampling without replacement from the diagnosis labels of each individual (1,000 times for the sex analyses and 10,000 times for the general case-control analyses). We chose 10,000 permutations for the case-control analyses, since this allows us to obtain p-values <1e-8, which is 1/(10,000*15,000), where 15,000 is the approximate number of genes tested (1,000 permutations allows us to obtain p-values<1e-7; since no gene reached near that p-value for the sex-specific analyses, we believed that 1,000 permutations would be sufficient). The relevant analysis procedures were then done in the standard way (as specified above) to obtain negative log p-values for each gene. The null distribution for the real data was then taken to be the full list of all negative log10 p-values across all permutations and all genes (i.e. the length of the list was the number of permutations times the number of genes). P-values for the real data were then calculated as the proportion of times the values (negative log10 p-values) of the null distribution list were higher than the value of the gene for the real data.

For the analyses of sex differences the permutations were done the same way except permuting the sexes within the controls and cases separately (and no permutations of diagnosis status). The sex specific analyses (see below) were then conducted in the same manner and empirical p-values for the real data were obtained with the same method as for the case-control differential expression.

### Leave One Out Analyses

The accuracy of genes obtained from each analysis was assessed by the ability of the genes to predict case-control or disease severity in left out datasets. For each analysis where this approach was conducted, the analysis was conducted with all datasets except one that was left out (alternating so that analyses were done with each dataset left out). The resulting gene sets were then used to create a “transcriptional score” for each individual specific to each cell type using the AddModuleScore_UCell from the UCell package (v1.3)^41^ with maxRank set to 16000 to ensure that all genes were used for the analyses. Scores of 0 were set to NA. UCell scores were normalized such that for each cell type, the minimum of the scores was subtracted from each score, and the results were then divided by the range of the scores for that cell type (maximum score minus minimum). Missing scores were then set to the mean of the scores of that cell type. When the gene set included multiple genes, a composite transcriptional score was created for each individual as the sum of the UCell scores across each cell type for up-regulated genes minus the sum of the UCell scores across each cell type for down-regulated genes (note: endothelial cells in Alzheimer’s disease datasets were not used due to incomplete coverage on all datasets for this cell type and the observation that including it decreased AUC).

A logistic regression model was created from the UCell scores of each individual and their diagnosis statuses using all datasets except the left out dataset. This model was then tested on the UCell scores and diagnosis statuses of the left out dataset with AUC determined from “auc” function of the pROC R package version 1.18.4^111^. To determine the ability of the genes to predict disease severity, a linear regression model was created from the UCell scores of each individual and their disease severities (Braak scores for Alzheimer’s disease, on a scale of 0 to 6, and a scale from 0 to 3 for COVID-19, with 1 indicating mild, 2 indicating moderate, 3 indicating severe based on clinical status of the patients). For the disease severity calculations only disease cases were used to prevent confounding from ability to predict general case vs. control status. For COVID-19 analyses, only datasets that had all cell types were used. For AD analyses, the Barker dataset was not used for disease severity calculations, because this dataset specifically focused on individuals with high Braak scores (some of whom had normal cognition and some of whom had impaired cognition).

We used the matrix of UCell scores for each individual across all datasets and all cell types and performed a heatmap using R with the settings symm=T and all other settings set to default. RCA Gene Lists were obtained specific for each cell type by using each individual gene to create a UCell score for each dataset and then following the same process as above. We separated the genes into up- and down-regulated sets based on whether the mean expression of the gene was higher in cases relative to controls or vice versa in all datasets. We then ranked each list by their mean AUC in predicting case-control status of the individuals in each dataset. These lists were called “RCA Gene List” throughout the paper. Relative Classification Accuracy was defined as the AUCs from the RCA Gene List, normalized by subtracting the minimum value for the particular disease and dividing by the range of AUCs for that disease.

Hoffman, Fujita, MathysCell, and Stephenson dataset individual down-samplings were performed by taking a random sample (with replacement) of cases and controls 20 times for each number of cases and controls and repeating the standard individual dataset analyses as described above. Cell number down-sampling was performed by randomly taking different proportions (0.001, 0.005, 0.001, 0.05, 0.1, 0.5) of cells from each dataset and then performing differential expression and SumRank meta-analyses as described above. AD datasets were also down-sampled one at a time either from the most reproducible (as measured by gene set AUC) or the least reproducible. SumRank meta-analysis was then performed with these down-sampled sets of datasets with 65% of datasets chosen unless this number was less than 7 in which case either 7 datasets were chosen or all datasets were chosen (if this was less than 7).

### Sex specific analyses

Two methods were used to determine sex-specific differential expression. The first method we call the Sex Interaction Method. In this method, differential expression was performed for each dataset with DESeq2 using the counts matrix of the data subsetted to cell type and using design = ∼Sex+Diagnosis+Sex:Diagnosis. This design tests the effect of each term (Sex, Diagnosis, and Sex:Diagnosis) on gene expression. We took the effect sizes of the interaction term (SexF.DiagnosisAD) and obtained their p-values for each dataset. The signed -log10(p-values) for each dataset were then combined using the SumRank meta-analysis method described above with p-values calibrated empirically using permutations as described above.

The second method we call the Composite Score Method. In this method, four different scores were combined to create a composite score. Standard differential expression was performed in DESeq2 between males and females in only cases (Score 1) and in only controls (Score 2) as well as cases vs controls in only males (Score 3) and in only females (Score 4). SumRank meta-analyses were then performed for each of these individual analysis types to obtain -log10(p-values). For female specificity the composite score was calculated as the sum of the -log10(p-values) of the females vs. males in cases with the -log10(p-values) of the cases vs. controls in females subtracted by the -log10(p-values) of the females vs. males in controls and the -log10(p-values) of the cases vs. controls in males (Score 1 + Score4 - Score3 - Score 2). For male specificity the composite score was calculated as the sum of the -log10(p-values) of the males vs. females in cases with the -log10(p-values) of the cases vs. controls in males subtracted by the -log10(p-values) of the males vs. females in controls and the -log10(p-values) of the cases vs. controls in females. These p-values were then calibrated empirically with permutations as described above. We looked for genes that had -log10(p-values)>3.65 in one of the analyses and were in the top 15 (0.1%) of genes in the other analysis.

For several of the COVID-19 datasets, some of the sex statuses of the individuals were not listed, so these were obtained by creating a composite RNA score of Y chromosome genes (*NLGN4Y*, *LINC00278*, *TTTY14*, *TMSB4Y*, *EIF1AY*, *USP9Y*, *KDM5D*, *ZFY*, *UTY*, *DDX3Y*, and *RPS4Y1*), which were able to differentiate sexes in the dataset well (total expression of these genes greater than 10 was defined as genetic male).

The ratio of mean expression of cases over mean expression of controls for females and males were calculated for plotting. The standard deviations for these were calculated by the error propagation formula as 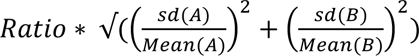, where Ratio is mean(A)/mean(B), and A is the expression in cases, while B is the expression in controls. Standard deviations were calculated separately for males and females and both were plotted.

### Human Genetic Comparisons

Significant genes from Genome Wide Association Studies (GWAS) of Alzheimer’s Disease^63–65^ and Parkinson’s Disease^55^ were inferred as the genes most proximal to the genome-wide significant genetic variants from the studies or those prioritized through various metrics by the study authors. Significant genes from AD whole-exome association studies^79–81^ were inferred as the genes with exons harboring the significant genetic variant. We assessed statistical significance of overlap of the meta-analysis genes with human genetic genes by Fisher’s exact test (two-sided) as implemented in R (fisher.test function).

### Gene Ontology Analyses

Cluster Profiler 4.0^53^ was used to find biological pathways with statistically significant enrichment from the meta-analysis gene sets. The organism was set to human (org.Hs.eg.db), ont (subontology) was set to BP (biological process), and pvaluecutoff was set to 0.05. The up- and down-regulated gene sets were analyzed with these settings, with the rest of the settings at default.

COVID-19 pathways were also analyzed by comparing the overlap of the up-regulated genes in each cell type to the gene sets derived from a database generated by Perturb-Seq experiments in which 6 cell lines were stimulated with different perturbations (interferon-beta, interferon-gamma, transforming growth factor beta 1, and tumor necrosis factor-alpha) and then had expression of individual genes knocked down with CRISPR guides to assess the effect of each gene on the perturbation response. This provided more specific gene sets for these pathways than could be obtained by standard gene ontology^54^. The specific pathways were coded as IFNG_REMOVE_IFNB; IFNB_REMOVE_IFNG; IFNG_REMOVE_TNFA; TNFA_REMOVE_IFNG; IFNB_REMOVE_TNFA; TNFA_REMOVE_IFNB; TNFA_REMOVE_TGFB1; TGFB1_REMOVE_TNFA, where each gene set was the genes involved in the specific perturbation pathway that were not involved in other pathways. The overlap of the meta-analysis up-regulated genes with the top 100 genes from each pathway was examined to determine more specifically the pathways involved in COVID-19 in each cell type, where the dominant pathway was determined as the pathway with the highest overlap after removing the genes from other pathways with high overlap.

### Overlap Calculations

Gene Set Enrichment Analysis (GSEA) was used as implemented in the Cluster Profiler 4.0^53^ and MSigDB v7.5.1^50^ packages. Gene Ontology gene sets were obtained with the msigdbr command with species “Homo sapiens”, category “C5”, subcategory “GO:BP”. GSEA was then performed on the ranked, signed -log10(p-values) of differential expression obtained from pseudo-bulked DESeq2 analyses (as in the Differential Expression section above) using the GSEA command with pvalueCutoff=1.1 and all other settings at default. The pvalueCutoff was set to 1.1 to ensure all pathways were present in the final output. Overlap of GSEA pathways was also performed after down-sampling datasets or individuals. In the first analysis, the top 20 up-regulated GSEA pathways (ranked by p-value) were obtained from the up-regulated genes after SumRank was performed with all AD datasets performing GSEA as above. This was then compared with the top 100 up-regulated pathways after performing SumRank with different numbers of datasets removed. These analyses were done with all broad cluster types (oligodendrocytes, astrocytes, OPCs, glutamatergic neurons, endothelial cells, GABA-ergic inhibitory neurons, and microglia) and for both up- and down-regulated genes and the average proportion of overlap was calculated. In the second analysis, the top 20 up-regulated GSEA pathways were obtained from the up-regulated genes of a random subset with 340 individuals (170 cases and 170 controls) of the MathysCell dataset. This was then compared with the top 100 up-regulated pathways based on differential expression with different down-samplings of individuals from the MathysCell dataset and proportion overlap was calculated in the same way. The same analyses were then done with down-regulated genes.

Gene set consistency was analyzed by obtaining the top 25 ranked genes after SumRank was performed with all AD datasets. The ranks of these genes were then obtained in the signed -log10(p-value) ordered SumRank genes after down-sampling datasets. These analyses were done with all broad cluster types and for both up- and down-regulated genes and each individual rank was plotted. The top 25 genes were obtained from the signed -log10(p-value) ordered genes of a random subset with 340 individuals (170 cases and 170 controls) of the MathysCell dataset. The ranks of these genes were then assessed in the signed -log10(p-value) ordered SumRank genes after down-sampling individuals. For each down-sampling the mean rank across all broad cluster types and up- and down-regulated genes were plotted.

### Mapping to Mathys et al. Nature, 2024 AD atlas

AD datasets were mapped to an atlas composed of the AD individuals from Mathys Nature 2024^51^ using the same FindTransfersAnchors approach as described above for major_cell_type and cell_type_high_resolution of the Mathys dataset. The data were then pseudobulked at the individual level within each cell type and reproducibility was assessed as described above.

### snATAC-Seq Analyses

Differentially accessible peaks from pseudo-bulked broad cell types were obtained from the Supplementary file of Xiong *et al*., Cell 2023^52^, who performed snATAC-seq on AD and control individuals and assessed for differential chromatin accessibility after pseudo-bulking to broad cell types (assigning cell types after integrating ATAC and RNA data). The genes corresponding to these peaks were found by overlapping the peak locations with the nearest gene promoter using the distancetoNearest function in the GenomicRanges R package version 3.2.0^112^ using hg38 annotations obtained through the AnnotationHub version 3.2.0 R package^113^. The genes corresponding to differentially accessible peaks were then ranked by p-value after subsetting to up- or down-regulated, and then overlapped for each cell type with meta-analysis DEGs after taking different percentages of the top ATAC-seq genes. Overlap enrichment was calculated by Fisher’s exact test. To find the DEGs from the Xiong RNA, we performed DESeq2 differential expression on the snRNA-seq of only the individuals that also had snATAC-seq performed on them. snATAC-seq peaks were visualized in UCSC Genome Browser hg38^114^.

### Memento Analyses

Memento version 0.12^43^ was used for differential expression analyses of PD data to compare with DESeq2 and logistic regression of pseudobulk data. Non-pseudobulked single-cell PD data were analyzed using the “fixed effects binary testing with multiple samples” method according to the Memento documentation tutorial with capture_rate of 0.07 and min_perc_group of 0.9 with all other settings left as default. Genes that were filtered out were set to effect size of 0 with p-value of 1. The de_coef value and de_pval were used for effect sizes and p-values, respectively, and SumRank was performed on the resulting Memento differential expression results as described above.

### Protein-Protein Interaction (PPI) Network Mapping

DEGs (BH corrected p-value<0.05) were input into STRING version 12.0^62^, which created PPI networks of the DEGs, connecting each DEG to another DEG if a PPI was known to exist between them. These networks were input into Cytoscape version 3.10.3^115^ and the “Analyze Network” function was used to find network characteristics for each gene (e.g. betweenness centrality). The ClueGO app of Cytoscape was used to analyze enriched GO pathways of the genes in the most densely connected networks.

### Mice

Mice were bred in-house or obtained from the Jackson Laboratory (JAX). Mice were housed in a 12-h light–dark cycle in a temperature-controlled and humidity-controlled environment with water and food provided ad libitum. Both males and females were used in this study. The following mouse strain was used: B6.Cg-Tg(APPSwFlLon,PSEN1*M146L*L286V)6799Vas/Mmjax (5xFAD; JAX 034848). For analysis of BCAT1 staining in oligodendrocytes, 8-10 month old mice were used. Animals were housed at New York University (NYU) Medical Center Animal Facility under specific pathogen–free conditions. All procedures were approved by the NYU School of Medicine Institutional Animal Care and Use Committee and complied with approved ethical regulations.

### Tissue Collection and Processing

Mice were perfused with cold 1xPBS followed by 4%PFA. Brains were removed, post fixed overnight, cryopreserved in 30% sucrose, and cryo-embedded in OCT. 40 µM coronal cryosections were generated between bregma 1.335-.745. For staining at least two sections containing mPFC were used for multiplexed IHC.

### Immunohistochemistry (IHC), imaging, and quantification

Coronal brain slices were rinsed 3x in PBS for 10 min each prior to antigen retrieval. For antigen retrieval slides were emersed in .1M citrate buffer, microwaved until boiling, and incubated for 15 minutes at 99°C in a water bath. Afterwards slides were returned to room temperature, rinsed 2x 10 min in PBS and blocked in 10% normal donkey serum (Jackson ImmunoResearch AB_2337258), 1% BSA, .25%tritonX100, with Mouse on Mouse IG blocking reagent (Vector Labs BMK-2202) in 1xPBS for 2hrs at room temperature. Sections were then stained with the following primary antibodies; Mouse anti CC1 (1:200, Sigma OP80), Goat anti SOX10 (1:200, R&D Systems AF2864-SP), and Rabbit anti BCAT1 (1:200, Proteintech 13640-1-AP) overnight in blocking solution with Mouse on Mouse protein concentrate instead of IG blocking reagent (Vector Labs BMK-2202) at 4°C. The next day sections were then washed 3x with PBST and incubated for 2hrs at RT with the following secondary antibodies all at 1:500; Alexa488 Donkey anti goat (Jackson ImmunoResearch 705-545-003), Alexa568 Donkey anti Mouse (Invitrogen A-31571), Alexa647 Donkey anti Rabbit (Jackson ImmunoResearch 711-605-152) in blocking solution with Mouse on Mouse protein concentrate (Vector Labs BMK-2202). Sections were then washed 3x with PBST and mounted with Fluoromount-G Mounting Medium, with DAPI (Invitrogen 00-4959-52). Z-stack tiled images of the mPFC were acquired using a LSM 800 Confocal microscope (Zeiss) using a 40x oil immersion objective (Na 1.3). Quantitative analysis was conducted on at least 2 slices per animal using the Fiji package for ImageJ software by a researcher blind to the experimental groups. After applying a median filter (2 pixel radius) to the *BCAT1* channel, SOX10+ CC1+ double positive oligodendrocyte cytoplasms were drawn by hand with the polygon tool. *BCAT1* mean fluorescent intensity was quantified per cell, normalized over *BCAT1* background staining, and averaged per animal. Data was expressed as FC over WT samples normalized for each batch of staining.

## Extended Data

**Extended Data Table 1.** Reproducibility of individual AD datasets by several metrics. For all analyses here the DEG lists included the same number of top genes (based on the 814 SumRank genes with -log10(p-value)>3.65). The mean number of DEGs per cell type is calculated from a q-value based FDR threshold of 0.05 after filtering out genes with logfc<0.25 and less than 10% detection in both cases and controls (reproducibility metrics with these DEGs are shown in Supplementary Table 13). Individual Gene AUC List is the list of genes ranked by their individual ability to distinguish cases from controls in all datasets. Relative Classification Accuracy is the normalized AUC of individual genes in their ability to distinguish diagnosis status in each dataset. Signed -log10(p-value)s were from comparisons of logistic regression models on disease status with and without each gene (see Methods for more details).

| Dataset | Mean AUC when using DEGs as a Group to Predict Diagnoses of Other Datasets | Specificity: Percentage of DEGs in Top 10% of Individual Gene AUC List | Mean Relative Classification Accuracy of Individual DEGs | Mean abs(log2fc) and signed -log10(p-value)s of individual genes in logistic regressions of diagnosis status in each dataset | Mean Correlation Between Predicted and Actual Disease Severity of Left-Out Datasets | Mean Number of DEGs per Cell Type | Number of AD Individuals | Number of Control Individuals | Total Number of Individuals | Mean nCount_RNA per Cell | Mean Number of Cells Per Individual |
| --- | --- | --- | --- | --- | --- | --- | --- | --- | --- | --- | --- |
| OteroGarcia | 0.55 | 38 | 41.1 | 0.08; 0.25 | NA | 182 | 8 | 8 | 16 | 2679 | 7506 |
| Leng_EC | 0.62 | 36 | 28.8 | 0.09; 0.09 | NA | 468 | 3 | 3 | 6 | 7399 | 6904 |
| Lau | 0.63 | 32 | 41.1 | 0.17; 0.43 | 0.27 | 0 | 12 | 9 | 21 | 4220 | 8165 |
| Gerrits_OTC | 0.63 | 31 | 32.2 | 0.09; 0.19 | -0.23 | 14 | 10 | 8 | 18 | 1790 | 19035 |
| YangCortex | 0.64 | 34 | 29.3 | 0.07; 0.12 | -0.15 | 160 | 4 | 4 | 8 | 7641 | 4603 |
| Grubman | 0.65 | 21 | 41.3 | 0.13; 0.25 | 0.18 | 72 | 6 | 6 | 12 | 1217 | 3550 |
| Zhou | 0.65 | 33 | 40.3 | 0.13; 0.37 | 0.18 | 234 | 21 | 11 | 32 | 1193 | 11672 |
| Sayed | 0.67 | 22 | 53.0 | 0.17; 0.62 | 0.30 | 1086 | 47 | 8 | 55 | 11899 | 6769 |
| Smith_EC | 0.68 | 23 | 38.5 | 0.13; 0.29 | 0.11 | 0 | 6 | 6 | 12 | 6908 | 3651 |
| Morabito | 0.69 | 29 | 41.2 | 0.12; 0.40 | 0.21 | 1 | 11 | 7 | 18 | 9681 | 6769 |
| Barker | 0.70 | 31 | 44.6 | 0.11; 0.29 | -0.26 | 0 | 9 | 9 | 18 | 1435 | 14494 |
| Sadick | 0.72 | 37 | 41.0 | 0.13; 0.37 | 0.22 | 5 | 10 | 6 | 16 | 1138 | 7383 |
| Mathys | 0.73 | 46 | 47.7 | 0.11; 0.35 | 0.34 | 0 | 24 | 24 | 48 | 3029 | 1472 |
| Gorabitto | 0.74 | 29 | 39.4 | 0.14; 0.49 | 0.19 | 1809 | 72 | 7 | 79 | 4073 | 15570 |
| MathysCell | 0.75 | 51 | 58.5 | 0.15; 0.68 | 0.30 | 160 | 252 | 175 | 427 | 10619 | 5489 |
| Fujita | 0.76 | 37 | 46.4 | 0.10; 0.50 | 0.31 | 81 | 286 | 156 | 442 | 12135 | 4705 |
| Hoffman | 0.80 | 51 | 54.7 | 0.19; 0.77 | NA | 1068 | 150 | 149 | 299 | 12291 | 4781 |
| Average | 0.68 | 34 | 42.3 | 0.12; 0.38 | 0.12 | 285 | 54.8 | 35.1 | 89.8 | 5844 | 7795 |

**Extended Data Table 2.**
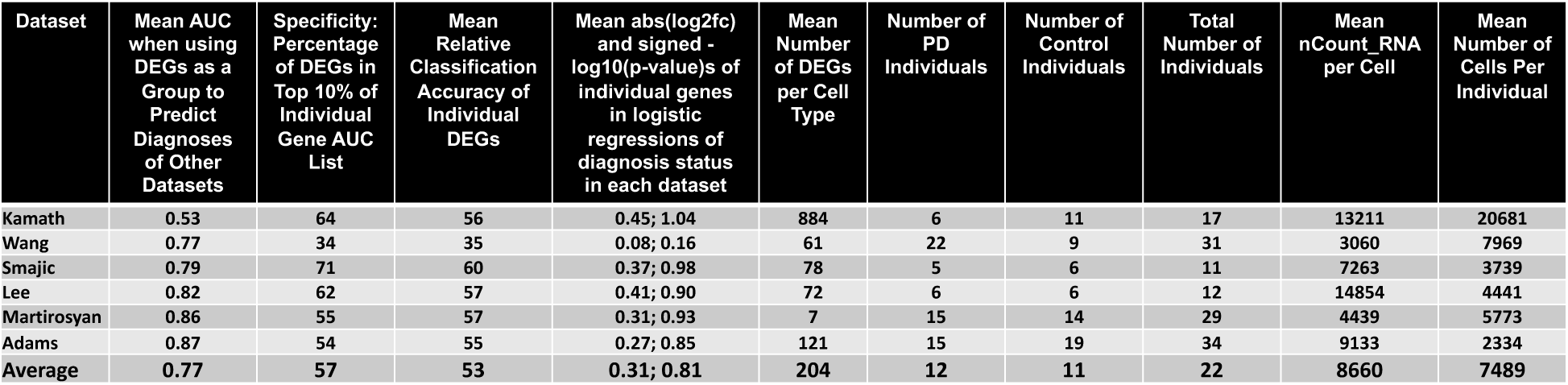
Reproducibility of individual PD datasets by several metrics. For all analyses here the DEG lists included the same number of top genes (based on the 1,527 SumRank genes with -log10(p-value)>3.35). The mean number of DEGs per cell type is calculated from a q-value based FDR threshold of 0.05 after filtering out genes with logfc<0.25 and less than 10% detection in both cases and controls (reproducibility metrics with these DEGs are shown in Supplementary Table 14). Individual Gene AUC List is the list of genes ranked by their individual ability to distinguish cases from controls in all datasets. Relative Classification Accuracy is the normalized AUC of individual genes in their ability to distinguish diagnosis status in each dataset. Signed -log10(p-value)s were from comparisons of logistic regression models on disease status with and without each gene (see Methods for more details).

| Dataset | Mean AUC when using DEGs as a Group to Predict Diagnoses of Other Datasets | Specificity: Percentage of DEGs in Top 10% of Individual Gene AUC List | Mean Relative Classification Accuracy of Individual DEGs | Mean abs(log2fc) and signed -log10(p-value)s of individual genes in logistic regressions of diagnosis status in each dataset | Mean Number of DEGs per Cell Type | Number of PD Individuals | Number of Control Individuals | Total Number of Individuals | Mean nCount_RNA per Cell | Mean Number of Cells Per Individual |
| --- | --- | --- | --- | --- | --- | --- | --- | --- | --- | --- |
| Kamath | 0.53 | 64 | 56 | 0.45; 1.04 | 884 | 6 | 11 | 17 | 13211 | 20681 |
| Wang | 0.77 | 34 | 35 | 0.08; 0.16 | 61 | 22 | 9 | 31 | 3060 | 7969 |
| Smajic | 0.79 | 71 | 60 | 0.37; 0.98 | 78 | 5 | 6 | 11 | 7263 | 3739 |
| Lee | 0.82 | 62 | 57 | 0.41; 0.90 | 72 | 6 | 6 | 12 | 14854 | 4441 |
| Martirosyan | 0.86 | 55 | 57 | 0.31; 0.93 | 7 | 15 | 14 | 29 | 4439 | 5773 |
| Adams | 0.87 | 54 | 55 | 0.27; 0.85 | 121 | 15 | 19 | 34 | 9133 | 2334 |
| Average | 0.77 | 57 | 53 | 0.31; 0.81 | 204 | 12 | 11 | 22 | 8660 | 7489 |

**Extended Data Table 3.** Reproducibility of individual COVID-19 datasets by several metrics. For all analyses here the DEG lists included the same number of top genes (based on the 937 SumRank genes with -log10(p-value)>3.90). The mean number of DEGs per cell type is calculated from a q-value based FDR threshold of 0.05 after filtering out genes with logfc<0.25 and less than 10% detection in both cases and controls (reproducibility metrics with these DEGs are shown in Supplementary Table 15). Individual Gene AUC List is the list of genes ranked by their individual ability to distinguish cases from controls in all datasets. Relative Classification Accuracy is the normalized AUC of individual genes in their ability to distinguish diagnosis status in each dataset. Signed - log10(p-value)s were from comparisons of logistic regression models on disease status with and without each gene (see Methods for more details). The datasets with NA for mean AUC have insufficient cells for at least one of the major cell types leading to inability to create reliable UCell scores for those datasets.

| Dataset | Mean AUC when using DEGs as a Group to Predict Diagnoses of Other Datasets | Specificity: Percentage of DEGs in Top 10% of Individual Gene AUC List | Mean Relative Classification Accuracy of Individual DEGs | Mean abs(log2fc) and signed -log10(p-value)s of individual genes in logistic regressions of diagnosis status in each dataset | Mean Correlation Between Predicted and Actual Disease Severity of Left-Out Datasets | Mean Number of DEGs per Cell Type | Number of COVID-19 Individuals | Number of Control Individuals | Total Number of Individuals | Mean nCount_RNA per Cell | Mean Number of Cells Per Individual |
| --- | --- | --- | --- | --- | --- | --- | --- | --- | --- | --- | --- |
| Su | 0.51 | 54 | 39.1 | 0.25; 0.55 | 0.23 | 402 | 129 | 16 | 145 | 2763 | 3859 |
| Schulteschrepping | 0.7 | 55 | 40.4 | 0.30; 0.78 | -0.23 | 1710 | 27 | 38 | 65 | 3901 | 3415 |
| Yu | 0.71 | 47 | 37.7 | 0.32; 0.53 | NA | 57 | 7 | 3 | 10 | 1008 | 34064 |
| Zhu | 0.71 | 57 | 42.6 | 0.40; 0.88 | -0.55 | 491 | 5 | 3 | 8 | 2079 | 4562 |
| Liao | 0.75 | 51 | 39.0 | 0.42; 0.72 | 0.13 | 423 | 9 | 4 | 13 | 5396 | 4528 |
| Trump | 0.76 | 41 | 34.4 | 0.25; 0.45 | 0.21 | 405 | 32 | 16 | 48 | 9962 | 1837 |
| Wen | 0.76 | 56 | 42.3 | 0.36; 0.69 | 0.05 | 147 | 10 | 5 | 15 | 376 | 3664 |
| Lee | 0.8 | 58 | 44.9 | 0.49; 0.79 | 0.19 | 42 | 11 | 4 | 15 | 6077 | 3993 |
| Wilk | 0.82 | 75 | 52.1 | 0.60; 1.31 | 0.21 | 436 | 7 | 6 | 13 | 2636 | 3398 |
| Arunachalam | 0.83 | 66 | 48.9 | 0.61; 1.14 | 0.17 | 482 | 7 | 5 | 12 | 7897 | 4972 |
| Combes | 0.84 | 69 | 49.7 | 0.59; 1.21 | -0.27 | 987 | 20 | 14 | 34 | 3331 | 2669 |
| Stephenson | 0.85 | 71 | 50.3 | 0.67; 1.22 | 0.2 | 783 | 86 | 23 | 109 | 2197 | 5853 |
| Bacher | NA | NA | 35.4 | 0.17; 0.28 | NA | 75 | 14 | 6 | 20 | 3892 | 5221 |
| Chua | NA | NA | 25.0 | 0.03; 0.11 | NA | 347 | 19 | 5 | 24 | 8419 | 6692 |
| Kusnadi | NA | NA | 32.0 | 0.28; 0.48 | NA | 227 | 37 | 9 | 46 | 5287 | 1829 |
| Meckiff | NA | NA | 32.4 | 0.20; 0.41 | NA | 344 | 37 | 9 | 46 | 7132 | 2904 |
| Average | 0.75 | 58 | 40.4 | 0.37; 0.72 | 0.03 | 460 | 28.6 | 10.4 | 38.9 | 4522 | 5841 |

**Extended Data Figure 1.**
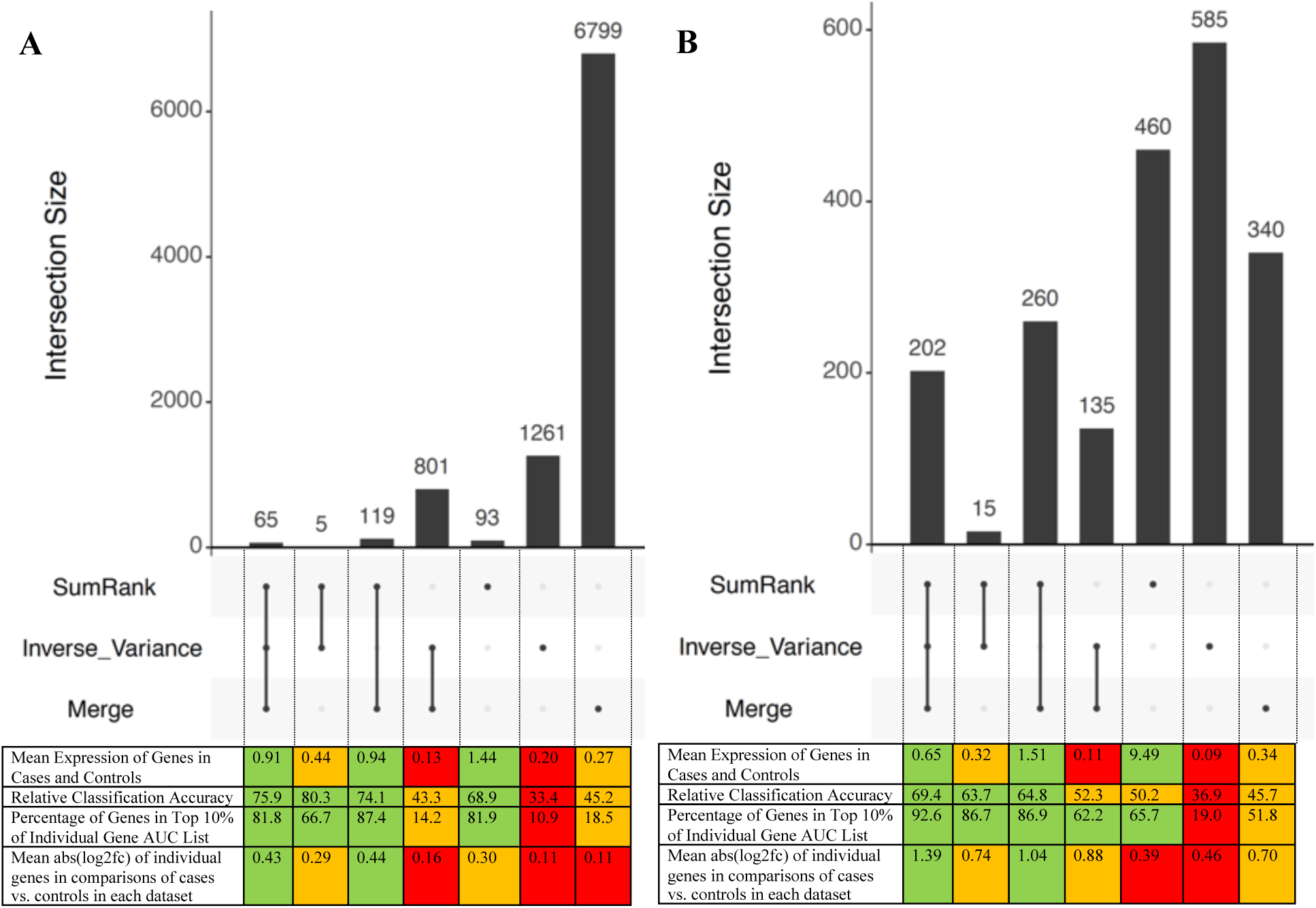
UpSet R plots^44^ of AD and COVID-19 genes discovered with different meta-analysis methods.. **A)** Plot of AD genes discovered based on a q-value based FDR cutoff of 0.05 used in all meta-analyses. **B)** Plot of COVID-19 genes discovered between the meta-analysis methods using the same number of genes for all meta-analyses (based on the 937 SumRank genes with -log10(p-value)>3.90). The plots show the intersection of genes discovered between the meta-analysis methods and the mean expression of the genes, relative classification accuracy (the normalized mean AUC of the individual genes in ability to predict diagnoses in all datasets), percentage of genes in top 10% of RCA Gene List, and mean abs(log2fc) of individual genes in comparisons of cases vs. controls in each dataset. Results are taken across all cell types. Color coding is based on the relative quality of the value, with green indicating the best values, orange indicating moderate, and red indicating poor.

**Extended Data Figure 2.**
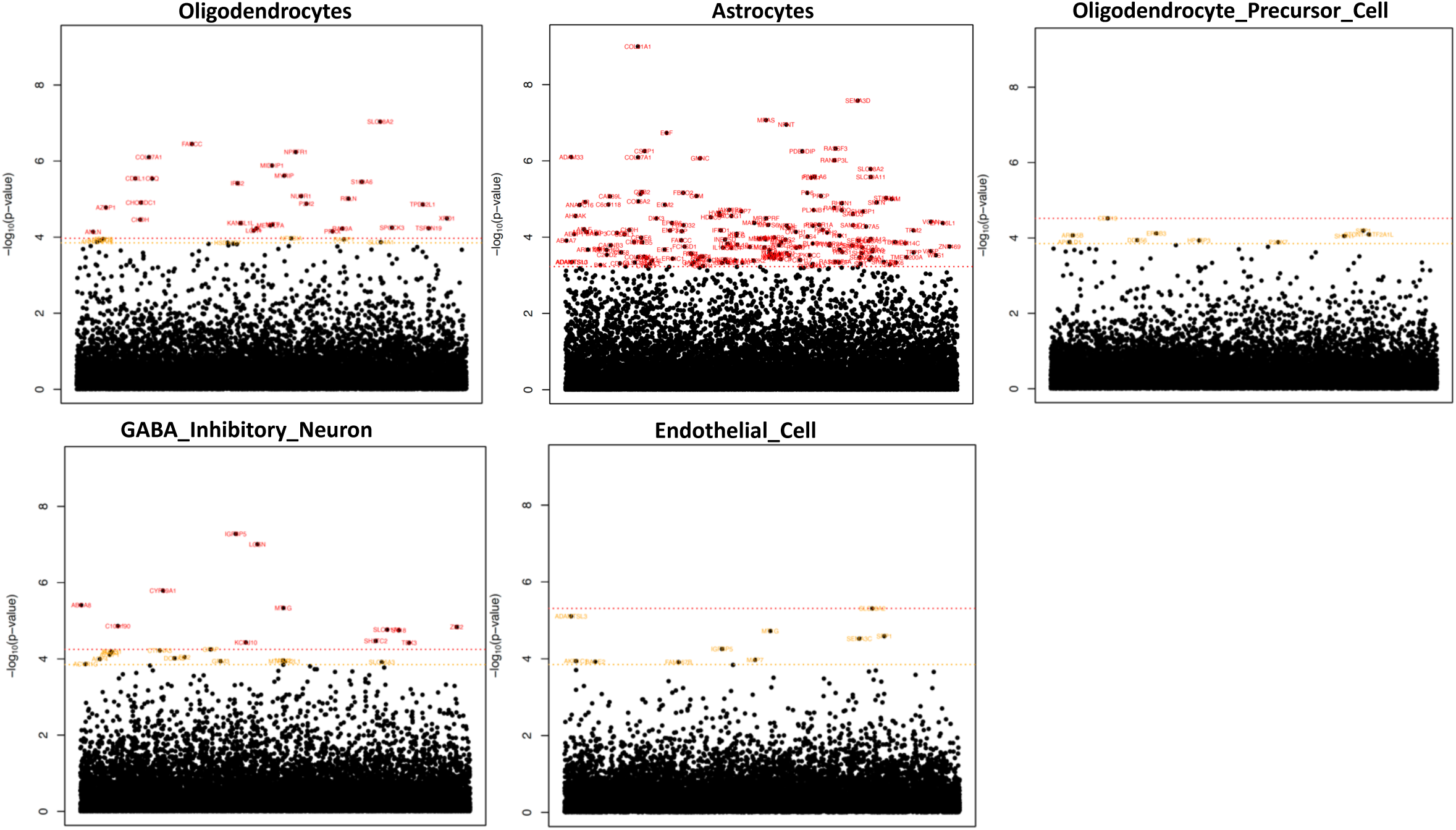
Manhattan plots of up-regulated genes in AD. Microglia and glutamatergic neurons are shown in Figure 4. Significance threshold is in red with 0.05 FDR cutoff (Benjamini-Hochberg). In orange is a - log10(p-value) cutoff that maximizes AUC (3.65 for AD; not shown if it is higher than the FDR cutoff red line). The x-axis are genes arranged in alphabetical order. Supplementary Data File 3 provides all genes with their p-values.

**Extended Data Figure 3.**
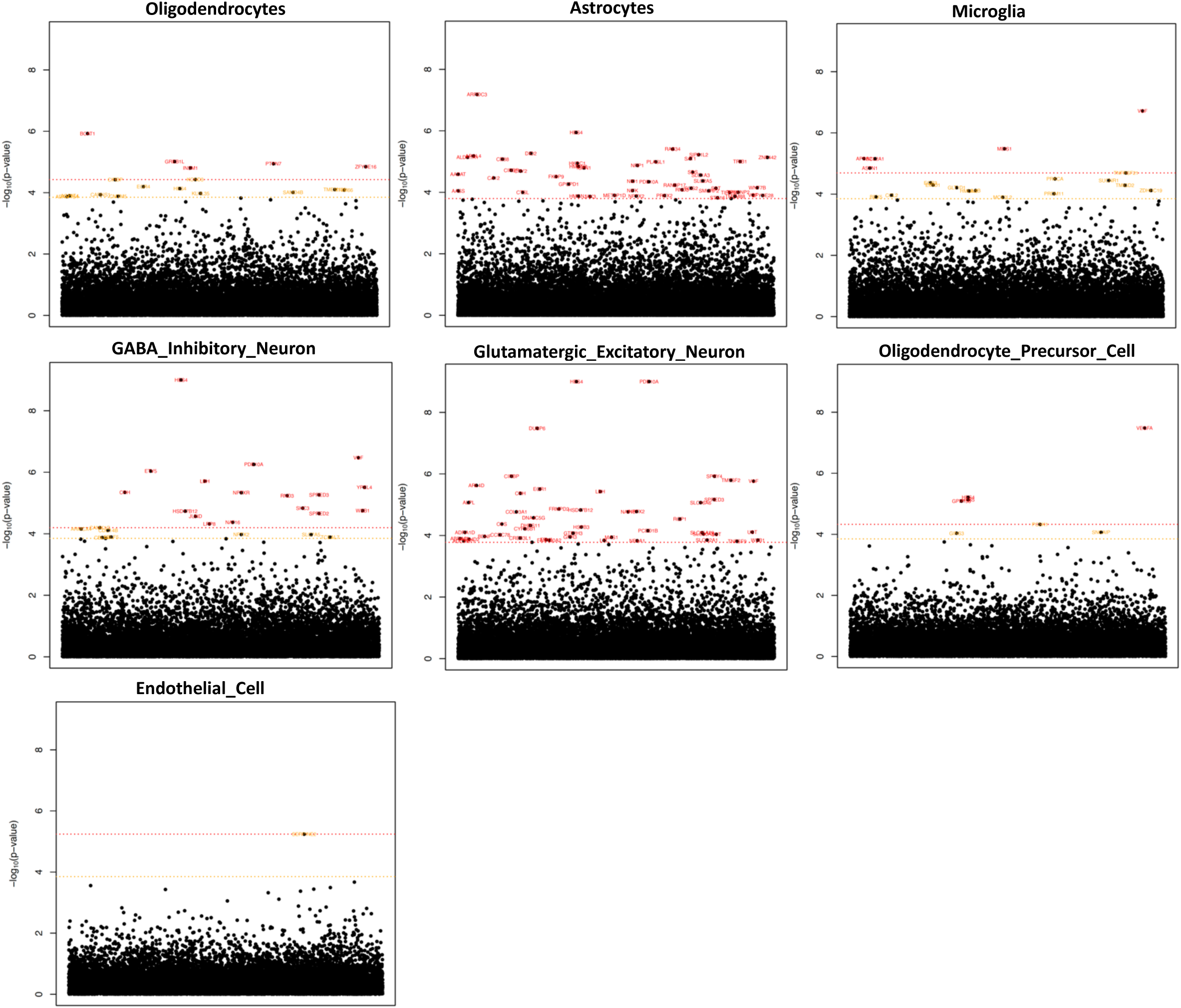
Manhattan plots of down-regulated genes in AD. Significance threshold is in red with 0.05 FDR cutoff (Benjamini-Hochberg). In orange is a -log10(p-value) cutoff that maximizes AUC (3.65 for AD; not shown if it is higher than the FDR cutoff red line). The x-axis are genes arranged in alphabetical order. Supplementary Data File 3 provides all genes with their p-values.

**Extended Data Figure 4.**
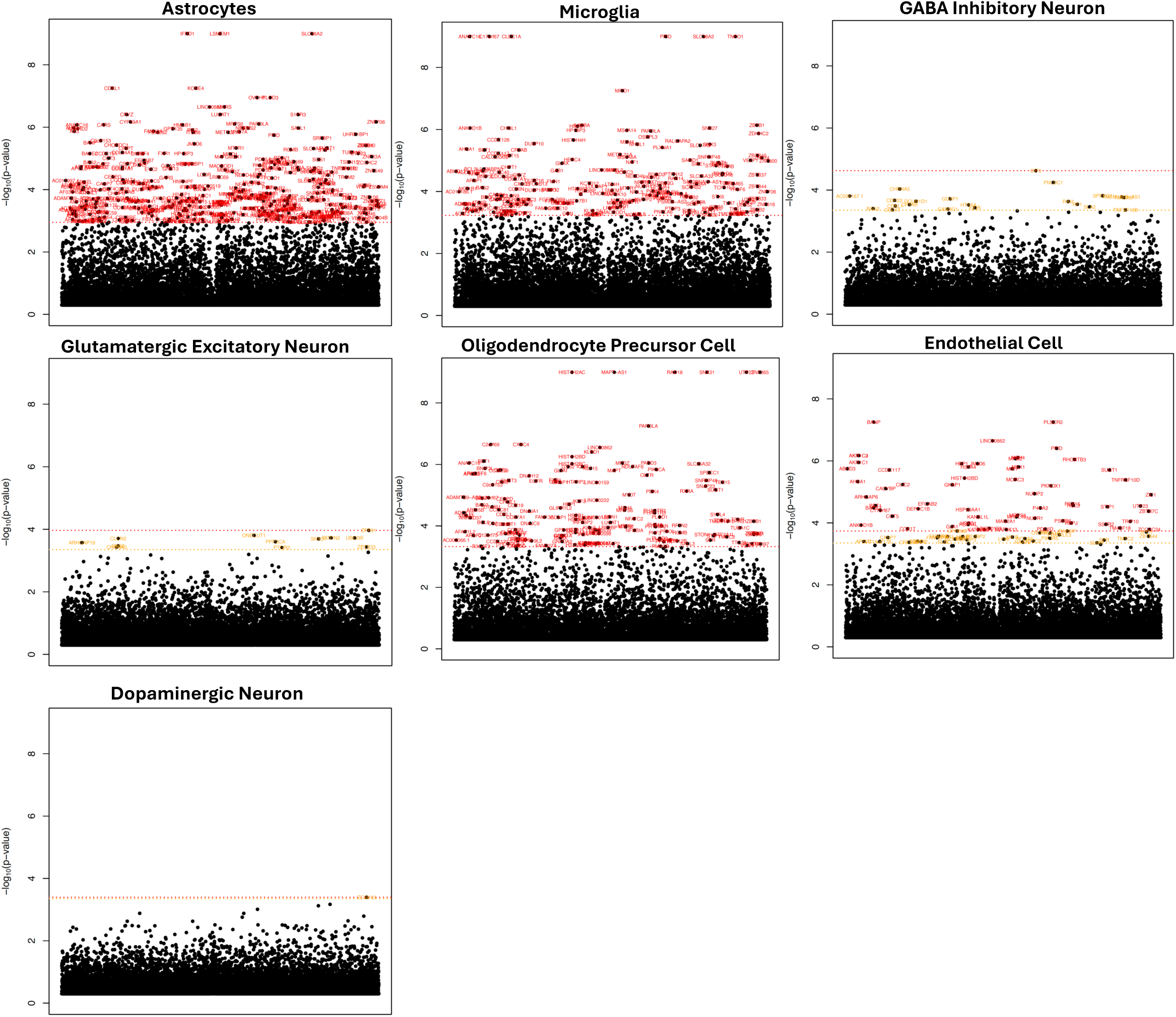
Manhattan plots of up-regulated genes in PD. Oligodendrocytes are shown in Figure 4. Significance threshold is in red with 0.05 FDR cutoff (Benjamini-Hochberg). In orange is a -log10(p-value) cutoff that maximizes AUC (3.35 for PD; not shown if it is higher than the FDR cutoff red line). The x-axis are genes arranged in alphabetical order. Supplementary Data File 4 provides all genes with their p-values.

**Extended Data Figure 5.**
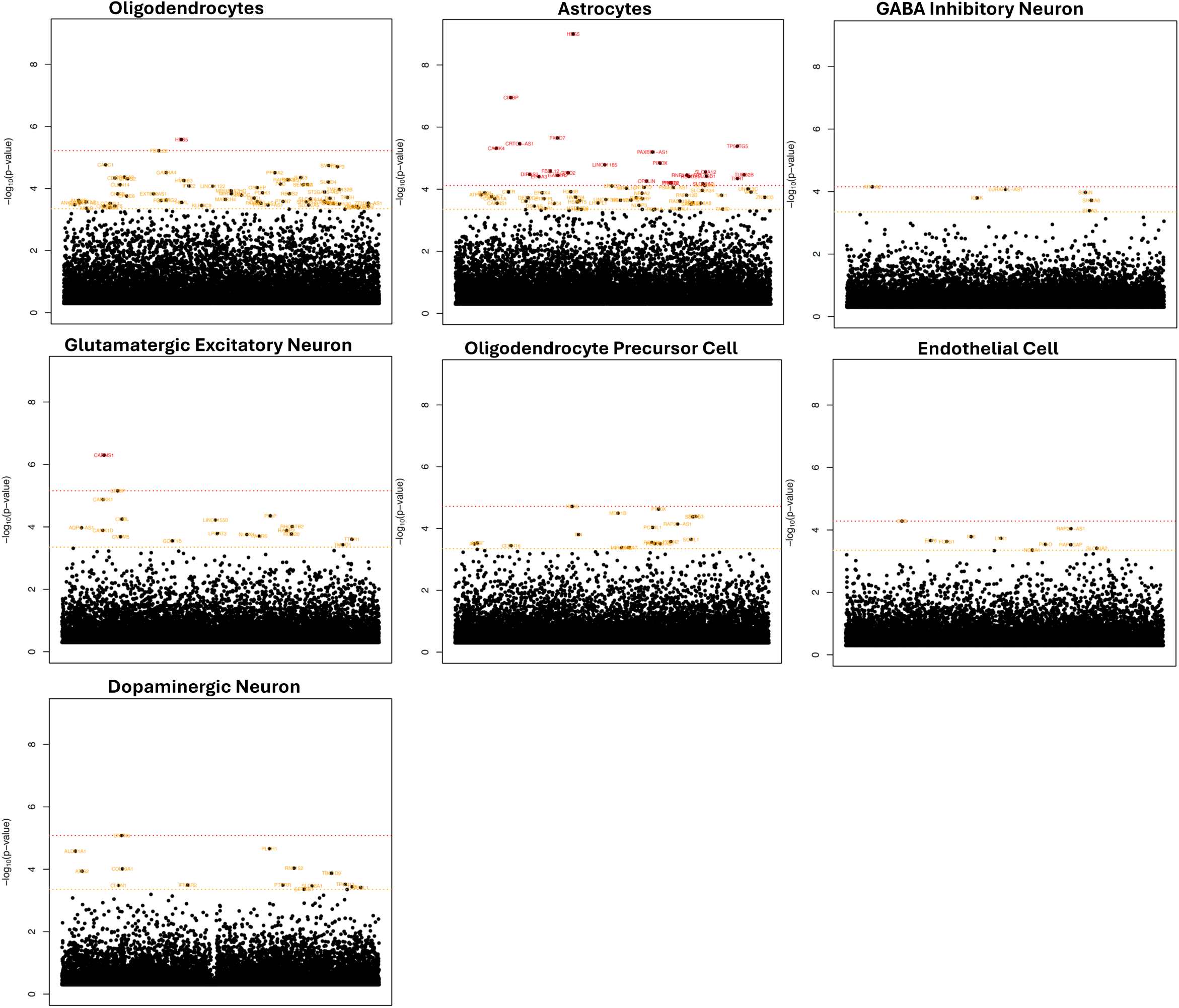
Manhattan plots of down-regulated genes in PD. Microglia are shown in Figure 4. Significance threshold is in red with 0.05 FDR cutoff (Benjamini-Hochberg). In orange is a -log10(p-value) cutoff that maximizes AUC (3.35 for PD; not shown if it is higher than the FDR cutoff red line). The x-axis are genes arranged in alphabetical order. Supplementary Data File 4 provides all genes with their p-values.

**Extended Data Figure 6.**
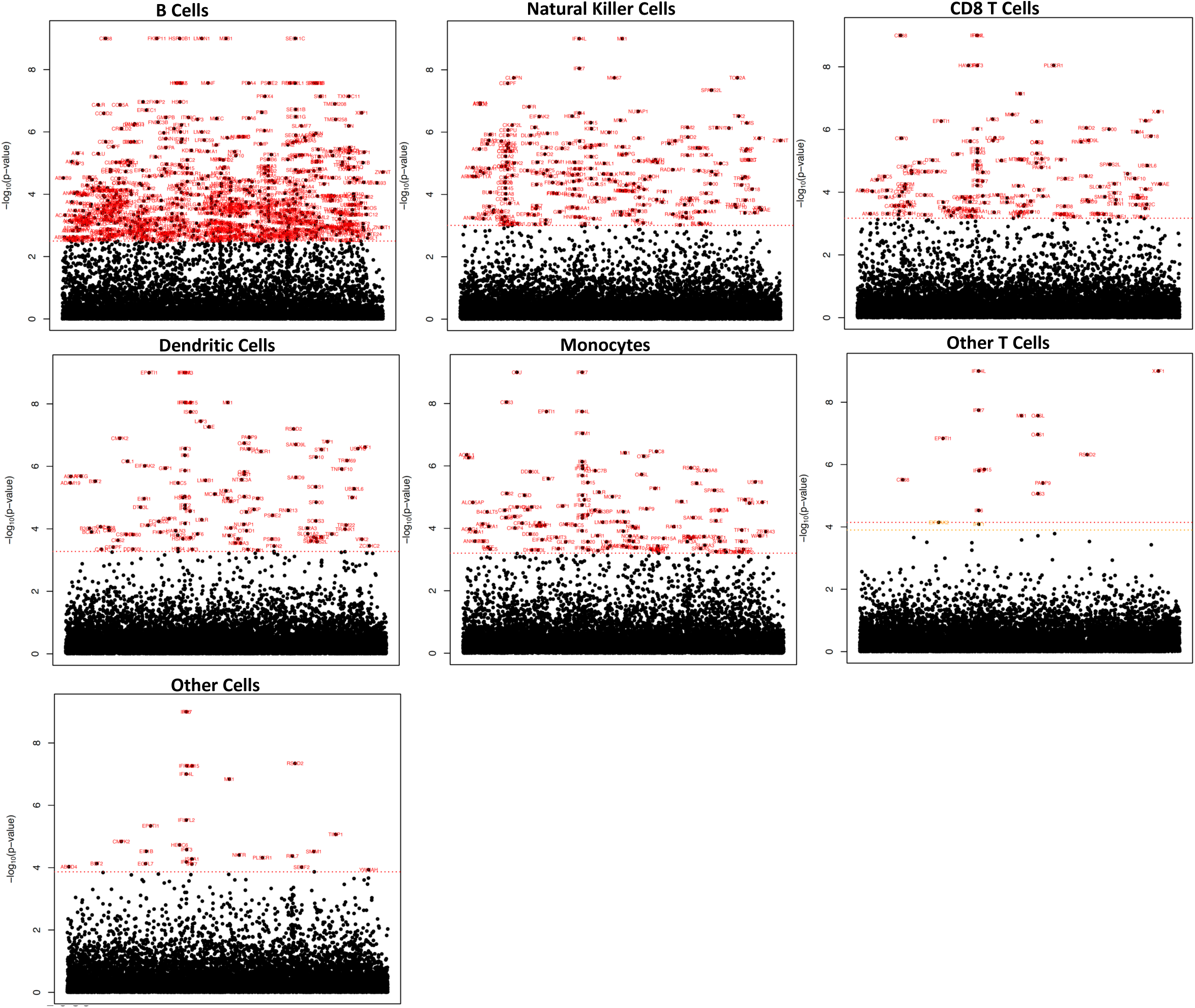
Manhattan plots of up-regulated genes in COVID-19. CD4 T cells are shown in Figure 4. Significance threshold is in red with 0.05 FDR cutoff (Benjamini-Hochberg). In orange is a -log10(p-value) cutoff that maximizes AUC (3.90 for COVID-19; not shown if it is higher than the FDR cutoff red line). The x-axis are genes arranged in alphabetical order. Supplementary Data File 5 provides all genes with their p-values.

**Extended Data Figure 7.**
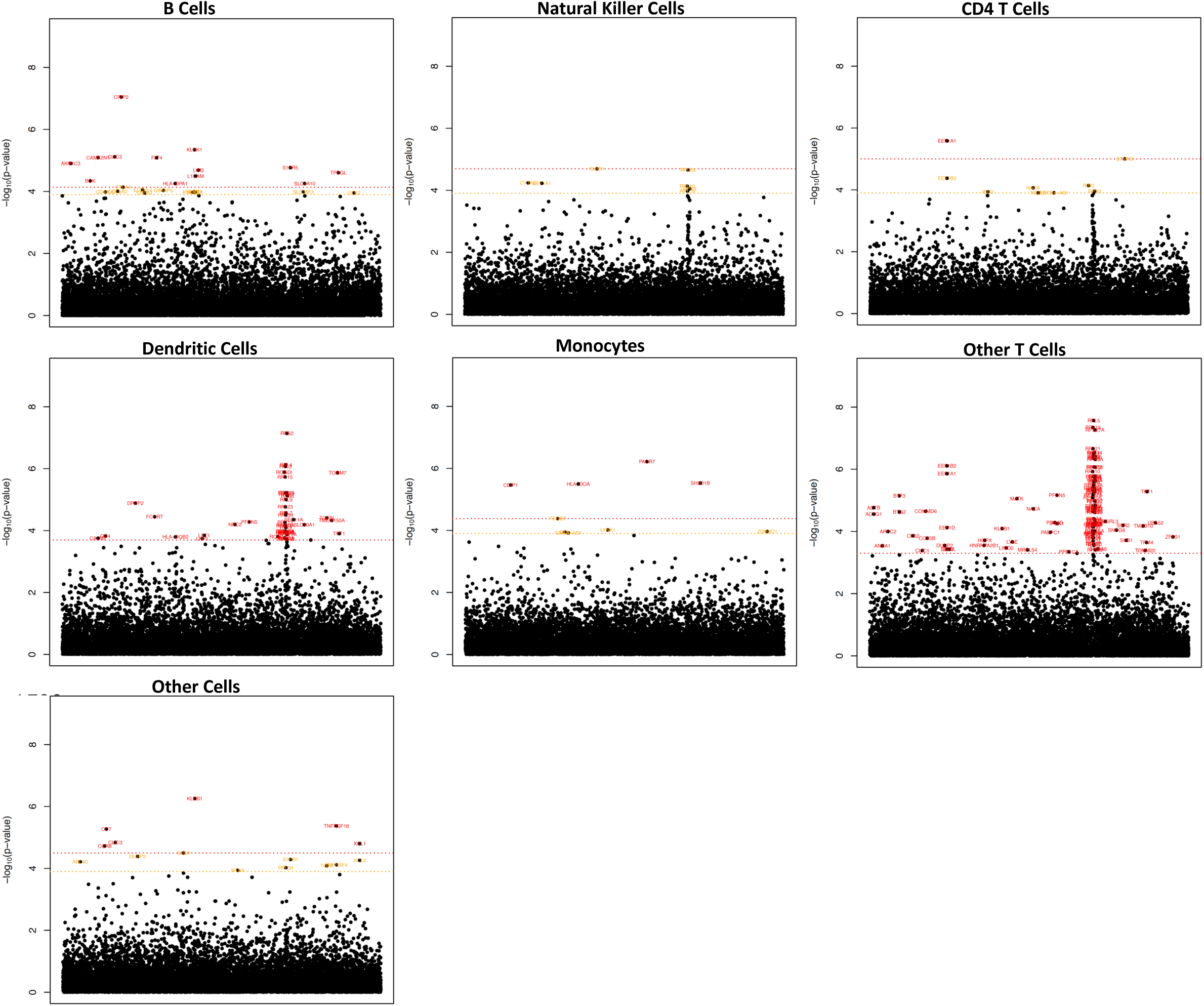
Manhattan plots of down-regulated genes in COVID-19. CD8 T cells are shown in Figure 4. Significance threshold is in red with 0.05 FDR cutoff (Benjamini-Hochberg). In orange is a -log10(p-value) cutoff that maximizes AUC (3.90 for COVID-19; not shown if it is higher than the FDR cutoff red line). The x-axis are genes arranged in alphabetical order. Supplementary Data File 5 provides all genes with their p-values.

## Supplementary Information

**Legends for Supplementary Data Files**

**Supplementary Data File 1**

Meta-data about all datasets used in this study.

**Supplementary Data File 2**

AUCs from SumRank meta-analyses in AD, PD, SCZ and COVID-19 with different percentage thresholds and p-value cutoffs.

**Supplementary Data File 3**

Output of AD meta-analyses, including p-values and effect sizes for all genes across all cell types, PPI network results, and correlation of scores with disease severity.

**Supplementary Data File 4**

Output of PD meta-analyses, including p-values and effect sizes for all genes across all cell types and PPI network results.

**Supplementary Data File 5**

Output of COVID-19 meta-analyses, including p-values and effect sizes for all genes across all cell types and correlation of scores with disease severity.

**Supplementary Data File 6**

Output of HD meta-analyses, including p-values and effect sizes for all genes across all cell types and PPI network results.

**Supplementary Data File 7**

Output of SCZ meta-analyses, including p-values and effect sizes for all genes across all cell types.

**Supplementary Data File 8**

Output of AD meta-analyses after mapping to MathysNaure2024.

**Supplementary Data File 9**

Output of gene ontology (GO) pathways for AD, PD, HD, SCZ and COVID-19 and shared DEGs between AD, PD, and HD.

**Supplementary Data File 10**

Overlap of COVID-19 DEGs with gene sets generated by a Perturb-Seq experiment.

**Supplementary Data File 11**

Output of AD and COVID-19 sex-difference meta-analyses, including p-values for all genes across all cell types.

## Supplementary Note

### Ruzicka et al. Schizophrenia Analyses

We observed a discrepancy between our results of differential expression in individual datasets and those of Ruzicka *et al*.^4^. In particular, they used a modified version of the muscat^116^ workflow and reported 6,056 DEGs in the McLean cohort and 2,666 DEGs in the Mt Sinai cohort across 25 cell types, while our analysis using DESeq2 and a q-value based lfdr cutoff of 0.05 only produced 14 DEGs across 7 cell types when using their dataset and combining the two cohorts. To understand this discrepancy, we first split the Ruzicka datasets into the McLean and MtSinai cohorts (as done in their study) and ran our DESeq2 pipeline. This produced only 79 DEGs for the McLean cohort and 1 DEG for the Mt Sinai cohort. We then ran our DESeq2 pipeline using Azimuth higher resolution cell types (19 cell types) and obtained 345 DEGs for the McLean cohort and 1 DEG for the Mt Sinai cohort. When we used our DESeq2 pipeline with the 25 Ruzicka cell type labels, we obtained 611 DEGs for the McLean cohort and 0 DEGs for the Mt Sinai cohort, showing that cell type labels are not the primary driver of the differences.

We then compared our differential expression pipelines. We followed the methods of Ruzicka *et al*. and used the limma-trend^109^ method in muscat for differential expression after pseudobulking using the mean of log-transformed counts with the Ruzicka cell labels, removing SZ3, SZ15, SZ24, SZ29, and SZ33, and using limma::removeBatchEffect to account for age, sex, PMI, and umi count, as done in their manuscript. We obtained 5,456 DEGs for the McLean cohort and 2,848 DEGs for the Mt Sinai cohort at a q-value based lfdr<0.05, approximately the same as the Ruzicka study (with 90.3% of these DEGs being shared with the Ruzicka DEGs), showing that we could approximately reproduce their results. We then used the same Ruzicka muscat pipeline but used summation of counts (rather than log-transformed counts) for pseudobulking and DESeq2 for differential expression. We obtained 2,474 DEGs for the McLean cohort and 5 DEGs for the MtSinai cohort, more similar to the numbers of our pipeline (which also uses summation of raw counts and DESeq2). When we used the mean of counts (rather than log-transformed counts) with limma-trend, we obtained 362 DEGs for the McLean cohort and 163 DEGs for the MtSinai cohort, though with evidence for poorer fits (increased numbers of genes filtered out).

We then ran the data through the recommended muscat tutorial (https://www.bioconductor.org/packages/devel/bioc/vignettes/muscat/inst/doc/analysis.html), which uses summation of raw counts for pseudobulk and differential expression with the default settings (i.e. logistic regression). When removeBatchEffect is not used to regress out covariates, we obtained 994 DEGs for the McLean cohort and 9 DEGs for the MtSinai cohort based on q-value based lfdr<0.05 (16 and 0 DEGs are obtained with adjusted p-value<0.05/25, correcting for number of cell types tested). When we used limma::removeBatchEffect as above to correct the counts matrix we obtained 1,240 DEGs for the McLean cohort and 1 DEG for the MtSinai cohort. When we used the mean of raw counts for pseudobulk, we obtained 9 DEGs for the McLean cohort and 0 DEGs for the MtSinai cohort, and when we used mean of logcounts for psuedobulk, we obtained 0 DEGs for the McLean cohort and 0 DEGs for the MtSinai cohort. In conclusion, we found that our method for DEG calling in individual datasets was more conservative than the Ruzicka method and that parameter choice had a substantial effect on the number of DEGs in these individual dataset analyses with the Ruzicka method of using limma-trend with pseudobulk of log-transformed counts producing substantially more DEGs than other methods but still with low relative reproducibility across datasets (see below). It will be important for future studies to evaluate the relative merits and disadvantages of both differential expression approaches.

Most importantly, however, we emphasize that the differences in calling DEGs in individual datasets do not affect any of the key conclusions in our study. The SumRank method evaluates relative ranks across datasets without using any threshold cutoffs (i.e. the entire set of genes are used), and our reproducibility assessments used equal numbers of genes per dataset. Our conclusions about SCZ’s relative lower reproducibility compared to other diseases were based on using the same pipeline in each disease. We chose the number of meta-analysis DEGs to maximize reproducibility (i.e. adding more DEGs did not increase AUC). When we split up the Ruzicka dataset into the 2 different cohorts and ran our analyses treating them as different datasets, the meta-analysis maximum AUC did not increase (max AUC of 0.59 using genes at -log10(p-value) cutoff of 3.5 vs 0.62 with them combined as one dataset), and the individual Ruzicka datasets only have marginally increased AUCs (Ruzicka_MtSinai=0.52, Ruzicka_McLean=0.55, Batiuk=0.58, Ling=0.63). When using the separated Ruzicka cohorts with Azimuth higher resolution cell types, the meta-analysis AUC does not increase (0.58). When using Ruzicka cell type labels and our DESeq2 pipeline for differential expression then choosing the top ranking genes as DEGs, we found that the maximum AUC of MtSinai cohort for predicting McLean phenotypes was 0.68 and 0.61 of McLean cohort for predicting MtSinai phenotypes (here we tried different numbers of DEGs and found the max AUC at 20 up- and 20 down-regulated genes for each cell type), still much below those of AD datasets with similar sample sizes. When we used the DEGs from the Ruzicka manuscript, the AUC of Mt Sinai cohort for predicting McLean phenotypes was 0.59, and the AUC of the McLean cohort for predicting MtSinai phenotypes was 0.63. We thus believe the results still support SCZ as a disease with lower reproducibility of differential expression than AD, PD, and COVID-19, a finding consistent with Figure 6 of Ruzicka *et al*.

### HD FoxP2 Analyses

We sought to understand the dramatic number of down-regulated DEGs specifically in FoxP2 neurons in HD. While this was analogous to the pattern with glutamatergic neurons in AD (few up-regulated DEGs and many down-regulated DEGs), the magnitude of this difference was much more stark. We first assessed the mapping and cell type labeling of these cells. We observed that the Lim dataset had poor mapping for these cells, particularly in the HD individuals, because the HD cells labeled as FoxP2 neurons in this dataset did not show clear expression of FoxP2 marker genes (Supplementary Figures 18-19). We thus removed those cells from our analyses. However, we found that even with the decreased power (only 3 datasets), the effect still remained (the results with the Lim FoxP2 neurons removed are shown throughout the paper; the results without the Lim dataset removed are in Supplementary Figure 20).

It is currently unclear why HD produces such a dramatic number of down-regulated DEGs in FoxP2 neurons (without a similar number of up-regulated DEGs). We observed clear evidence of conserved cell type markers for FoxP2 neurons in the HD cells (Supplementary Figure 18), and the cell type mapping scores of FoxP2 cells are similar to those of other cell types (Supplementary Table 22), except for the Lim dataset, which was excluded from HD analyses for FoxP2. We also assessed the total number of RNA molecules (nCount_RNA) and unique RNA molecules (nFeature_RNA) that each cell type produced to assess whether FoxP2 cells might have globally diminished RNA expression. However, we found that although HD cells generally had lower total and unique RNA molecules, the FoxP2 neurons did not have a bigger decrease in RNA than that of other cell types (Supplementary Tables 23-24 and Supplementary Figure 20). We also observed that the proportion of FoxP2 neurons (relative to other cell types) decreased in HD, but this decrease was less than that of D2 MSNs (Supplementary Table 25), which did not show the dramatic number of down-regulated DEGs. In addition, a decreased number of cells should not, by itself, cause down-regulated DEGs in pseudobulk differential expression methods. Overall, then, it appears that HD produces a specific change in FoxP2 neurons that causes them to down-regulate many genes that appear to be related to important functions of these cells, but the mechanism of this is currently unclear. This is an important area for future research.

## Supplementary Figures

**Supplementary Table 1.** Reproducibility of genes in AD datasets using DESeq2 q-value adjusted p-values to define DEGs (FDR<0.05). Genes with logfc<0.25 and less than 10% detection in both cases and controls were filtered out.

| Cell Type | 0 | 1 | 2 | 3 | 4 | 5 | 6 | 7 | 8 | 9 | 10 | 11 | 12 | 13 | 14 | 15 | 16 | 17 |
| --- | --- | --- | --- | --- | --- | --- | --- | --- | --- | --- | --- | --- | --- | --- | --- | --- | --- | --- |
| Oligodendrocytes | 26426 | 3470 | 424 | 64 | 11 | 7 | 0 | 0 | 0 | 0 | 0 | 0 | 0 | 0 | 0 | 0 | 0 | 0 |
| Astrocytes | 26706 | 3050 | 551 | 73 | 17 | 4 | 1 | 0 | 0 | 0 | 0 | 0 | 0 | 0 | 0 | 0 | 0 | 0 |
| Oligodendrocyte Precursor Cells | 28762 | 1571 | 65 | 4 | 0 | 0 | 0 | 0 | 0 | 0 | 0 | 0 | 0 | 0 | 0 | 0 | 0 | 0 |
| Glutamatergic Excitatory Neurons | 22140 | 6319 | 1546 | 328 | 56 | 11 | 2 | 0 | 0 | 0 | 0 | 0 | 0 | 0 | 0 | 0 | 0 | 0 |
| Endothelial Cells | 30166 | 233 | 3 | 0 | 0 | 0 | 0 | 0 | 0 | 0 | 0 | 0 | 0 | 0 | 0 | 0 | 0 | 0 |
| GABA Inhibitory Neurons | 22110 | 5815 | 1918 | 462 | 83 | 11 | 3 | 0 | 0 | 0 | 0 | 0 | 0 | 0 | 0 | 0 | 0 | 0 |
| Microglia | 26788 | 3242 | 311 | 47 | 13 | 1 | 0 | 0 | 0 | 0 | 0 | 0 | 0 | 0 | 0 | 0 | 0 | 0 |

Number of Up or Down-Regulated Genes Present in Each Number of Datasets

**Supplementary Table 2.** Reproducibility of genes in PD datasets using DESeq2 q-value adjusted p-values to define DEGs (FDR<0.05). Genes with logfc<0.25 and less than 10% detection in both cases and controls were filtered out.

| Cell Type | 0 | 1 | 2 | 3 | 4 | 5 | 6 |
| --- | --- | --- | --- | --- | --- | --- | --- |
| Oligodendrocytes | 32850 | 2471 | 286 | 37 | 2 | 0 | 0 |
| Astrocytes | 34143 | 1417 | 81 | 5 | 0 | 0 | 0 |
| Oligodendrocyte Precursor Cells | 34989 | 642 | 14 | 1 | 0 | 0 | 0 |
| Glutamatergic Excitatory Neurons | 34626 | 1006 | 14 | 0 | 0 | 0 | 0 |
| Endothelial Cells | 35311 | 319 | 16 | 0 | 0 | 0 | 0 |
| GABA Inhibitory Neurons | 34579 | 1055 | 12 | 0 | 0 | 0 | 0 |
| Microglia | 34688 | 950 | 8 | 0 | 0 | 0 | 0 |
| Dopaminergic Neurons | 34793 | 853 | 0 | 0 | 0 | 0 | 0 |

Number of Up or Down-Regulated Genes Present in Each Number of Datasets

**Supplementary Table 3.** Reproducibility of genes in COVID-19 datasets using DESeq2 q-value adjusted p-values to define DEGs (FDR<0.05). Genes with logfc<0.25 and less than 10% detection in both cases and controls were filtered out.

| Cell Type | 0 | 1 | 2 | 3 | 4 | 5 | 6 | 7 | 8 | 9 | 10 | 11 | 12 | 13 | 14 | 15 | 16 |
| --- | --- | --- | --- | --- | --- | --- | --- | --- | --- | --- | --- | --- | --- | --- | --- | --- | --- |
| B Cells | 15272 | 3485 | 1434 | 720 | 427 | 363 | 332 | 85 | 12 | 4 | 0 | 0 | 0 | 0 | 0 | 0 | 0 |
| NK Cells | 18871 | 2630 | 383 | 120 | 61 | 45 | 17 | 5 | 2 | 0 | 0 | 0 | 0 | 0 | 0 | 0 | 0 |
| CD8 T Cells | 16981 | 4024 | 843 | 194 | 51 | 21 | 10 | 7 | 3 | 0 | 0 | 0 | 0 | 0 | 0 | 0 | 0 |
| Dendritic Cells | 20704 | 1159 | 192 | 46 | 18 | 6 | 5 | 3 | 1 | 0 | 0 | 0 | 0 | 0 | 0 | 0 | 0 |
| Other | 19438 | 2253 | 371 | 55 | 9 | 6 | 0 | 1 | 0 | 1 | 0 | 0 | 0 | 0 | 0 | 0 | 0 |
| CD4 T Cells | 15822 | 4647 | 1172 | 317 | 111 | 36 | 19 | 7 | 2 | 0 | 1 | 0 | 0 | 0 | 0 | 0 | 0 |
| Monocytes | 13709 | 4879 | 2161 | 822 | 344 | 136 | 45 | 22 | 10 | 4 | 2 | 0 | 0 | 0 | 0 | 0 | 0 |
| Other T Cells | 19495 | 2461 | 144 | 22 | 9 | 2 | 0 | 1 | 0 | 0 | 0 | 0 | 0 | 0 | 0 | 0 | 0 |

Number of Up or Down-Regulated Genes Present in Each Number of Datasets

**Supplementary Table 4.** Reproducibility of genes in SCZ datasets using DESeq2 q-value adjusted p-values to define DEGs (FDR<0.05). Genes with logfc<0.25 and less than 10% detection in both cases and controls were filtered out.

| Cell Type | 0 | 1 | 2 | 3 |
| --- | --- | --- | --- | --- |
| Oligodendrocytes | 34840 | 0 | 0 | 0 |
| Astrocytes | 34840 | 0 | 0 | 0 |
| Oligodendrocyte Precursor Cells | 34840 | 0 | 0 | 0 |
| Glutamatergic Excitatory Neurons | 34838 | 2 | 0 | 0 |
| Endothelial Cells | 34840 | 0 | 0 | 0 |
| GABA Inhibitory Neurons | 34835 | 5 | 0 | 0 |
| Microglia | 34826 | 14 | 0 | 0 |

Number of Up or Down-Regulated Genes Present in Each Number of Datasets

**Supplementary Table 5.** Reproducibility of genes in HD datasets using DESeq2 q-value adjusted p-values to define DEGs (FDR<0.05). Genes with logfc<0.25 and less than 10% detection in both cases and controls were filtered out. Note: FOXP2 cells were not used for the Lim dataset, so that cell type only had 3 possible datasets.

| Cell Type | 0 | 1 | 2 | 3 | 4 |
| --- | --- | --- | --- | --- | --- |
| Oligodendrocytes | 25827 | 3164 | 538 | 127 | 10 |
| Astrocytes | 22298 | 5895 | 1161 | 277 | 35 |
| Oligodendrocyte Precursor Cells | 27283 | 2266 | 113 | 4 | 0 |
| Cholinergic Interneurons | 29666 | 0 | 0 | 0 | 0 |
| Ciliary Ependymal Cells | 28587 | 1025 | 52 | 2 | 0 |
| D1 Medium Spiny Neurons | 21440 | 7131 | 1043 | 48 | 4 |
| D2 Medium Spiny Neurons | 21868 | 7591 | 198 | 8 | 1 |
| FOXP2 Neurons | 21465 | 7544 | 604 | 53 | 0 |
| Endothelial Cells | 28934 | 702 | 28 | 1 | 1 |
| GABAergic Interneuron | 29454 | 209 | 3 | 0 | 0 |
| Microglia | 26975 | 2510 | 167 | 12 | 2 |
| Mural Cells | 29339 | 320 | 7 | 0 | 0 |
| PV Interneuron | 26929 | 2677 | 62 | 0 | 0 |
| Secretory Ependymal Cells | 28353 | 1279 | 34 | 0 | 0 |
| T Cells | 29665 | 1 | 0 | 0 | 0 |

Number of Up or Down-Regulated Genes Present in Each Number of Datasets

**Supplementary Table 6.** Reproducibility of genes in AD datasets using the top 200 genes of each dataset. Genes are ranked by p-value to define DEGs and genes with logfc<0.25 and less than 10% detection in both cases and controls were filtered out.

| Cell Type | 0 | 1 | 2 | 3 | 4 | 5 | 6 | 7 | 8 | 9 | 10 | 11 | 12 | 13 | 14 | 15 | 16 | 17 |
| --- | --- | --- | --- | --- | --- | --- | --- | --- | --- | --- | --- | --- | --- | --- | --- | --- | --- | --- |
| Oligodendrocytes | 25589 | 3963 | 639 | 122 | 60 | 18 | 7 | 2 | 1 | 1 | 0 | 0 | 0 | 0 | 0 | 0 | 0 | 0 |
| Astrocytes | 25533 | 3906 | 685 | 179 | 58 | 24 | 14 | 0 | 1 | 2 | 0 | 0 | 0 | 0 | 0 | 0 | 0 | 0 |
| Oligodendrocyte Precursor Cells | 25464 | 4180 | 607 | 118 | 25 | 6 | 0 | 2 | 0 | 0 | 0 | 0 | 0 | 0 | 0 | 0 | 0 | 0 |
| Glutamatergic Excitatory Neurons | 25730 | 3914 | 586 | 118 | 33 | 12 | 6 | 2 | 0 | 1 | 0 | 0 | 0 | 0 | 0 | 0 | 0 | 0 |
| Endothelial Cells | 25732 | 3991 | 594 | 73 | 10 | 1 | 0 | 1 | 0 | 0 | 0 | 0 | 0 | 0 | 0 | 0 | 0 | 0 |
| GABA Inhibitory Neurons | 25514 | 4194 | 550 | 100 | 29 | 9 | 4 | 1 | 0 | 1 | 0 | 0 | 0 | 0 | 0 | 0 | 0 | 0 |
| Microglia | 25361 | 4147 | 669 | 134 | 59 | 18 | 7 | 5 | 2 | 0 | 0 | 0 | 0 | 0 | 0 | 0 | 0 | 0 |

Number of Up or Down-Regulated Genes Present in Each Number of Datasets

**Supplementary Table 7.** Reproducibility of genes in PD datasets using the top 200 genes of each dataset. Genes are ranked by p-value to define DEGs and genes with logfc<0.25 and less than 10% detection in both cases and controls were filtered out.

| Cell Type | 0 | 1 | 2 | 3 | 4 | 5 | 6 |
| --- | --- | --- | --- | --- | --- | --- | --- |
| Oligodendrocytes | 33563 | 1828 | 205 | 41 | 6 | 3 | 0 |
| Astrocytes | 33630 | 1708 | 247 | 50 | 9 | 0 | 2 |
| Oligodendrocyte Precursor Cells | 33572 | 1816 | 202 | 44 | 12 | 0 | 0 |
| Glutamatergic Excitatory Neurons | 33519 | 1989 | 132 | 6 | 0 | 0 | 0 |
| Endothelial Cells | 33528 | 1895 | 173 | 42 | 7 | 1 | 0 |
| GABA Inhibitory Neurons | 33538 | 1964 | 130 | 14 | 0 | 0 | 0 |
| Microglia | 33526 | 1888 | 191 | 34 | 7 | 0 | 0 |
| Dopaminergic Neurons | 34003 | 1566 | 74 | 3 | 0 | 0 | 0 |

Number of Up or Down-Regulated Genes Present in Each Number of Datasets

**Supplementary Table 8.** Reproducibility of genes in COVID-19 datasets using the top 200 genes of each dataset. Genes are ranked by p-value to define DEGs and genes with logfc<0.25 and less than 10% detection in both cases and controls were filtered out.

| Cell Type | 0 | 1 | 2 | 3 | 4 | 5 | 6 | 7 | 8 | 9 | 10 | 11 | 12 | 13 | 14 | 15 | 16 |
| --- | --- | --- | --- | --- | --- | --- | --- | --- | --- | --- | --- | --- | --- | --- | --- | --- | --- |
| B Cells | 18617 | 2600 | 564 | 186 | 69 | 34 | 21 | 18 | 11 | 10 | 3 | 1 | 0 | 0 | 0 | 0 | 0 |
| NK Cells | 18684 | 2618 | 540 | 120 | 74 | 40 | 34 | 14 | 4 | 4 | 2 | 0 | 0 | 0 | 0 | 0 | 0 |
| CD8 T Cells | 18021 | 3237 | 588 | 152 | 67 | 33 | 22 | 11 | 2 | 0 | 1 | 0 | 0 | 0 | 0 | 0 | 0 |
| Dendritic Cells | 18494 | 2851 | 546 | 139 | 51 | 21 | 14 | 9 | 4 | 3 | 1 | 1 | 0 | 0 | 0 | 0 | 0 |
| Other | 18263 | 3073 | 603 | 141 | 31 | 11 | 7 | 2 | 2 | 1 | 0 | 0 | 0 | 0 | 0 | 0 | 0 |
| CD4 T Cells | 18288 | 3015 | 552 | 176 | 60 | 27 | 10 | 4 | 1 | 0 | 1 | 0 | 0 | 0 | 0 | 0 | 0 |
| Monocytes | 18812 | 2702 | 441 | 109 | 40 | 16 | 7 | 4 | 2 | 1 | 0 | 0 | 0 | 0 | 0 | 0 | 0 |
| Other T Cells | 18413 | 3006 | 471 | 126 | 51 | 35 | 18 | 10 | 4 | 0 | 0 | 0 | 0 | 0 | 0 | 0 | 0 |

Number of Up or Down-Regulated Genes Present in Each Number of Datasets

**Supplementary Table 9.** Reproducibility of genes in SCZ datasets using the top 200 genes of each dataset. Genes are ranked by p-value to define DEGs and genes with logfc<0.25 and less than 10% detection in both cases and controls were filtered out.

| Cell Type | 0 | 1 | 2 | 3 |
| --- | --- | --- | --- | --- |
| Oligodendrocytes | 34023 | 811 | 6 | 0 |
| Astrocytes | 33668 | 1144 | 28 | 0 |
| Oligodendrocyte Precursor Cells | 33716 | 1097 | 26 | 1 |
| Glutamatergic Excitatory Neurons | 33687 | 1129 | 23 | 1 |
| Endothelial Cells | 33674 | 1133 | 32 | 1 |
| GABA Inhibitory Neurons | 33931 | 897 | 10 | 2 |
| Microglia | 33651 | 1178 | 11 | 0 |

Number of Up or Down-Regulated Genes Present in Each Number of Datasets

**Supplementary Table 10.**
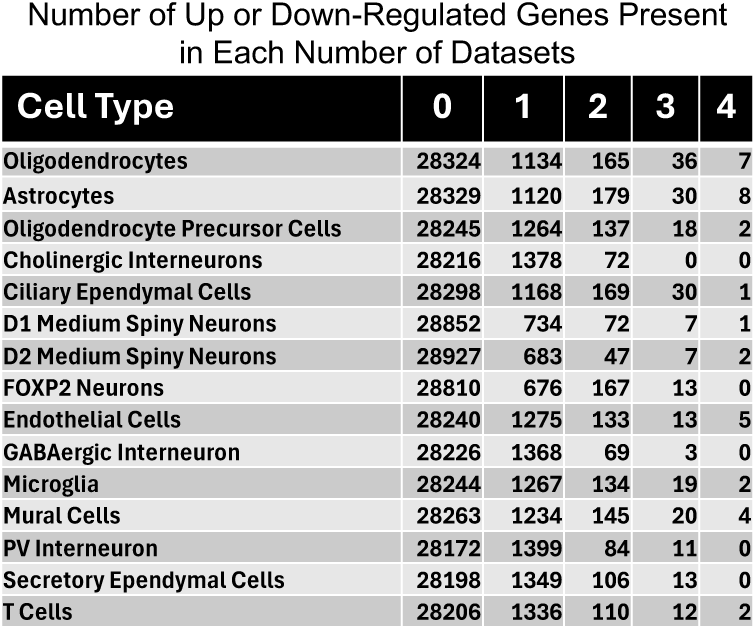
Reproducibility of genes in HD datasets using the top 200 genes of each dataset. Genes are ranked by p-value to define DEGs and genes with logfc<0.25 and less than 10% detection in both cases and controls were filtered out. Note: FOXP2 cells were not used for the Lim dataset, so that cell type only had 3 possible datasets.

**Supplementary Table 11.**
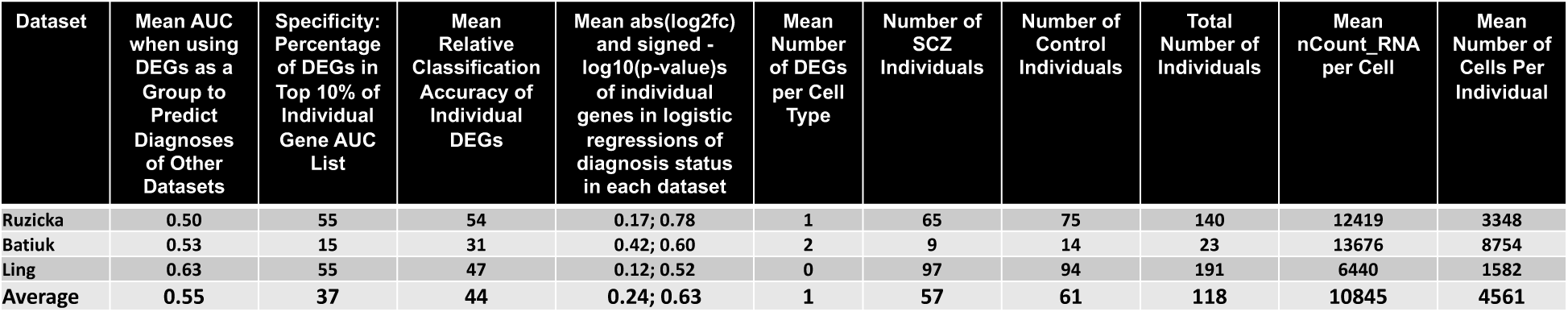
Reproducibility of individual SCZ datasets by several metrics. For all analyses here the DEG lists included the same number of top genes (based on the 98 SumRank genes with -log10(p-value)>3.40). The mean number of DEGs per cell type is calculated from a q-value based FDR threshold of 0.05 after filtering out genes with logfc<0.25 and less than 10% detection in both cases and controls (reproducibility metrics with these DEGs are not shown due to the very small number of DEGs meeting this criteria). Individual Gene AUC List is the list of genes ranked by their individual ability to distinguish cases from controls in all datasets. Relative Classification Accuracy is the normalized AUC of individual genes in their ability to distinguish diagnosis status in each dataset. Signed -log10(p-value)s were from comparisons of logistic regression models on disease status with and without each gene (see Methods for more details). We note that the Individual Gene AUC List and Relative Classification Accuracy are likely less accurate for SCZ than the other diseases due to the low number of datasets.

**Supplementary Table 12.** Reproducibility of individual HD datasets by several metrics. For all analyses here the DEG lists included the same number of top genes (based on the 837 SumRank genes with -log10(p-value)>3.30). The mean number of DEGs per cell type is calculated from a q-value based FDR threshold of 0.05 after filtering out genes with logfc<0.25 and less than 10% detection in both cases and controls (reproducibility metrics with these DEGs are not shown due to the very small number of DEGs meeting this criteria). Individual Gene AUC List is the list of genes ranked by their individual ability to distinguish cases from controls in all datasets. Relative Classification Accuracy is the normalized AUC of individual genes in their ability to distinguish diagnosis status in each dataset. Signed -log10(p-value)s were from comparisons of logistic regression models on disease status with and without each gene (see Methods for more details). We note that due to the substantial difference in size between the Handsaker dataset relative to the others, the mean AUCs for individual datasets are not easily comparable because the Handsaker DEGs do not need to predict the Handsaker dataset while the other datasets need to do so.

| Dataset | Mean AUC when using DEGs as a Group to Predict Diagnoses of Other Datasets | Specificity: Percentage of DEGs in Top 10% of Individual Gene AUC List | Mean Relative Classification Accuracy of Individual DEGs | Mean abs(log2fc) and signed -log10(p-value)s of individual genes in logistic regressions of diagnosis status in each dataset | Mean Number of DEGs per Cell Type | Number of HD Individuals | Number of Control Individuals | Total Number of Individuals | Mean nCount_RNA per Cell | Mean Number of Cells Per Individual |
| --- | --- | --- | --- | --- | --- | --- | --- | --- | --- | --- |
| Garcia | 0.80 | 50 | 52.9 | 0.51; 1.30 | 87 | 5 | 5 | 10 | 10250 | 4815 |
| Lim | 0.82 | 53 | 52.2 | 0.74; 1.87 | 447 | 6 | 5 | 11 | 9509 | 2618 |
| Matsushima | 0.84 | 50 | 50.3 | 0.53; 1.46 | 276 | 7 | 8 | 15 | 7273 | 4730 |
| Handsaker | 0.96 | 39 | 45.4 | 0.55; 2.84 | 2695 | 50 | 53 | 103 | 10884 | 5643 |
| Average | 0.85 | 48 | 50.2 | 0.58; 1.87 | 877 | 54.8 | 35.1 | 89.8 | 9479 | 4452 |

**Supplementary Figure 1.**
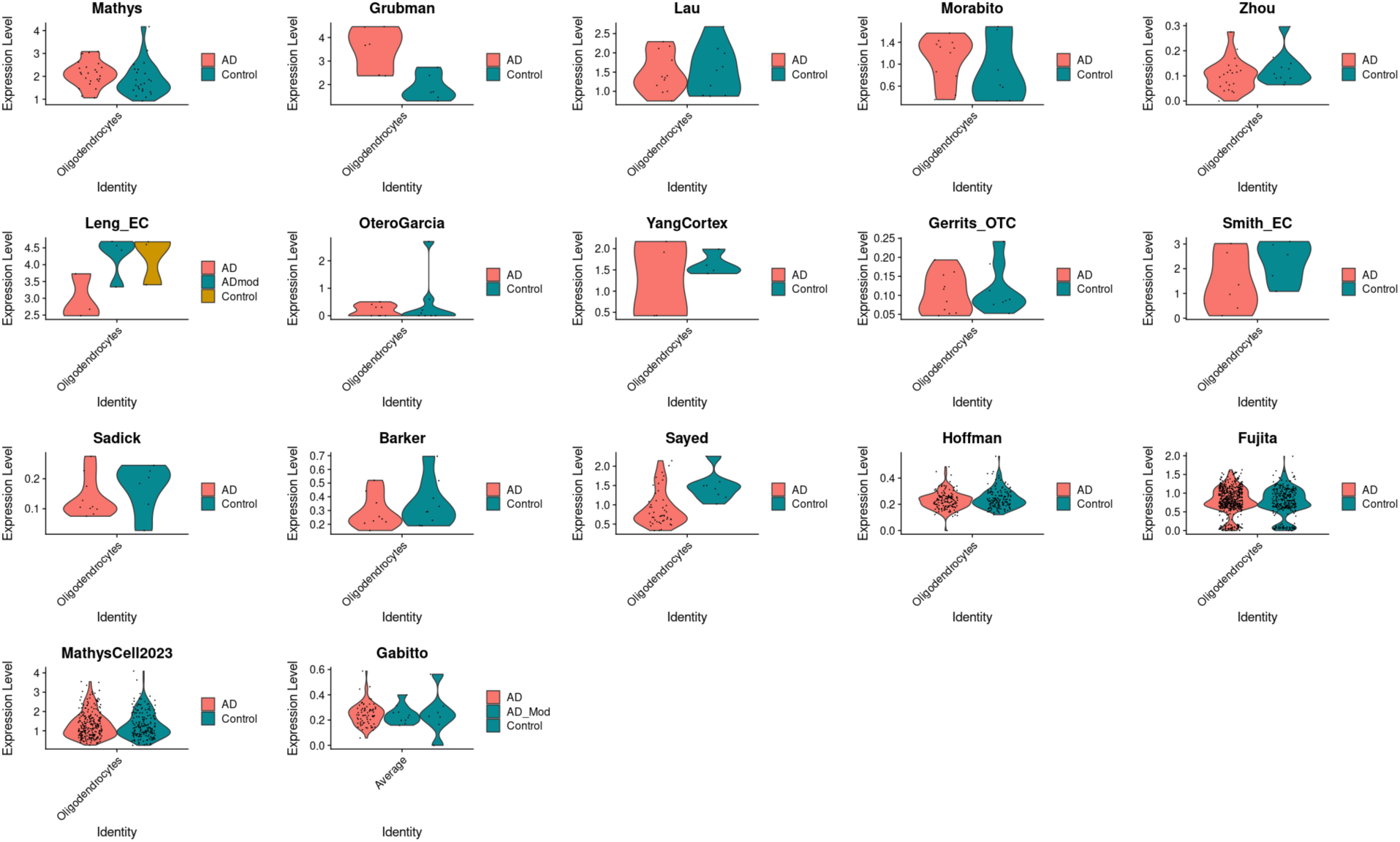
Violin plots of expression of the *LINGO1* gene in AD datasets.

**Supplementary Table 13.** Reproducibility of individual AD datasets by several metrics with q-value based DEGs. For all analyses here the DEG lists were determined by a q-value based FDR threshold of 0.05 after filtering out genes with logfc<0.25 and less than 10% detection in both cases and controls. RCA Gene List is the list of genes ranked by their individual ability to distinguish cases from controls in all datasets. Relative Classification Accuracy is the normalized AUC of individual genes in their ability to distinguish diagnosis status in each dataset. Signed - log10(p-value)s were from comparisons of logistic regression models on disease status with and without each gene (see Methods for more details). The datasets with NA have 0 DEGs at this threshold.

| Dataset | Mean AUC when using DEGs as a Group to Predict Diagnoses of Other Datasets | Specificity: Percentage of DEGs in Top 10% of Individual Gene AUC List | Mean Relative Classification Accuracy of Individual DEGs | Mean abs(log2fc) and signed - log10(p-value)s of individual genes in logistic regressions of diagnosis status in each dataset | Mean Correlation Between Predicted and Actual Disease Severity of Left-Out Datasets | Mean Number of DEGs per Cell Type |
| --- | --- | --- | --- | --- | --- | --- |
| OteroGarcia | 0.53 | 38 | 40.5 | 0.06; 0.38 | 0.10 | 182 |
| Gerrits_OTC | 0.58 | 34 | 40.8 | 0.15; 0.34 | -0.19 | 14 |
| Smith_EC | 0.59 | 39 | 81.7 | 0.24; 0.39 | 0.06 | 0 |
| Sadick | 0.61 | 52 | 55.2 | 0.11; 0.52 | -0.25 | 5 |
| Morabito | 0.62 | 49 | 55.1 | 0.10; 0.49 | 0.05 | 1 |
| YangCortex | 0.63 | 18 | 28.7 | 0.09; 0.18 | -0.10 | 160 |
| Zhou | 0.65 | 26 | 40.3 | 0.07; 0.26 | 0.16 | 234 |
| Grubman | 0.67 | 29 | 35.0 | 0.10; 0.29 | 0.08 | 72 |
| Lau | 0.69 | 80 | 84.0 | 0.00; 0.80 | 0.21 | 0 |
| SeaAD | 0.69 | 22 | 32.7 | 0.06; 0.22 | 0.24 | 1809 |
| Leng_EC | 0.70 | 19 | 29.1 | 0.10; 0.19 | -0.19 | 468 |
| Sayed | 0.72 | 33 | 40.3 | 0.05; 0.33 | 0.29 | 1086 |
| Hoffman | 0.76 | 48 | 46.6 | 0.09; 0.48 | -0.24 | 1068 |
| MathysCell2023 | 0.76 | 55 | 54.4 | 0.12; 0.55 | 0.22 | 160 |
| Fujita | 0.77 | 74 | 51.6 | 0.20; 0.74 | 0.34 | 81 |
| Mathys | NA | NA | NA | NA; NA | NA | 0 |
| Barker | NA | NA | NA | NA; NA | NA | 0 |
| Average | 0.66 | 41 | 47.7 | 0.10; 0.41 | 0.05 | 314 |

**Supplementary Table 14.**
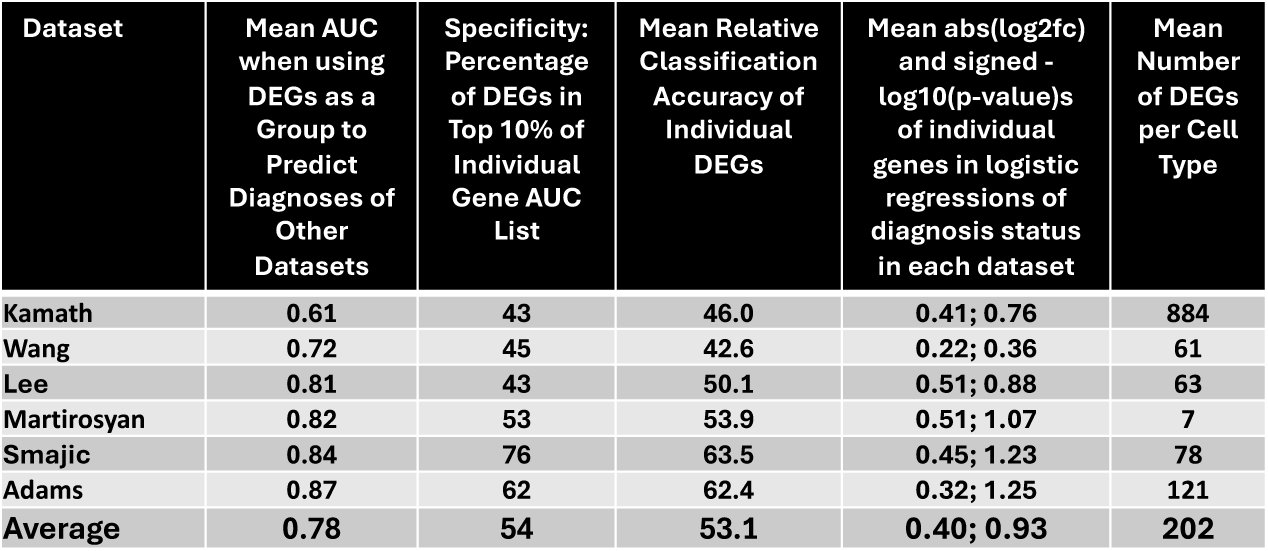
Reproducibility of individual PD datasets by several metrics with q-value based DEGs. For all analyses here the DEG lists were determined by a q-value based FDR threshold of 0.05 after filtering out genes with logfc<0.25 and less than 10% detection in both cases and controls. RCA Gene List is the list of genes ranked by their individual ability to distinguish cases from controls in all datasets. Relative Classification Accuracy is the normalized AUC of individual genes in their ability to distinguish diagnosis status in each dataset. Signed - log10(p-value)s were from comparisons of logistic regression models on disease status with and without each gene (see Methods for more details).

| Dataset | Mean AUC when using DEGs as a Group to Predict Diagnoses of Other Datasets | Specificity: Percentage of DEGs in Top 10% of Individual Gene AUC List | Mean Relative Classification Accuracy of Individual DEGs | Mean abs(log2fc) and signed - log10(p-value)s of individual genes in logistic regressions of diagnosis status in each dataset | Mean Number of DEGs per Cell Type |
| --- | --- | --- | --- | --- | --- |
| Kamath | 0.61 | 43 | 46.0 | 0.41; 0.76 | 884 |
| Wang | 0.72 | 45 | 42.6 | 0.22; 0.36 | 61 |
| Lee | 0.81 | 43 | 50.1 | 0.51; 0.88 | 63 |
| Martirosyan | 0.82 | 53 | 53.9 | 0.51; 1.07 | 7 |
| Smajic | 0.84 | 76 | 63.5 | 0.45; 1.23 | 78 |
| Adams | 0.87 | 62 | 62.4 | 0.32; 1.25 | 121 |
| Average | 0.78 | 54 | 53.1 | 0.40; 0.93 | 202 |

**Supplementary Table 15.**
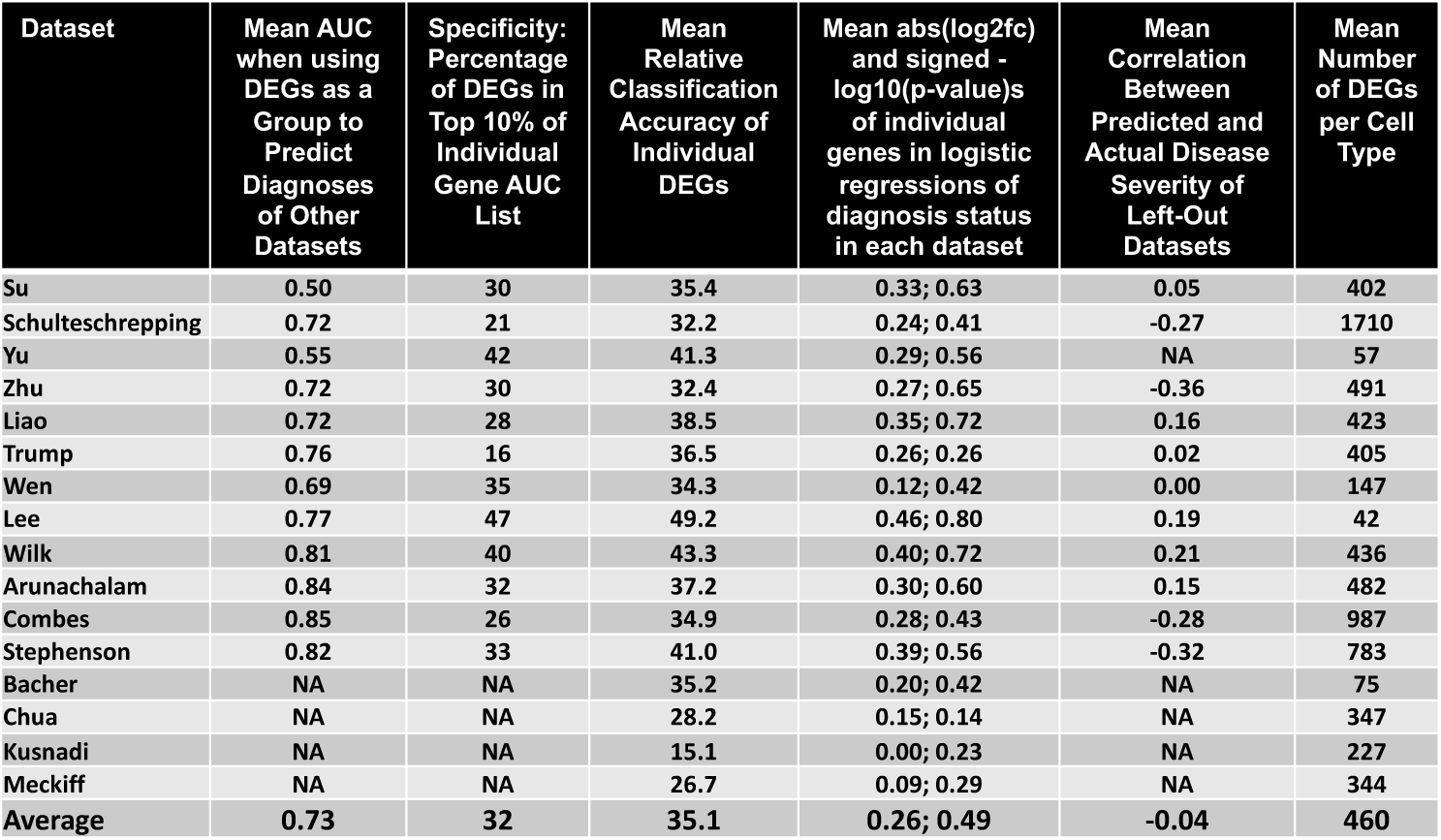
Reproducibility of individual COVID-19 datasets by several metrics with q-value based DEGs. For all analyses here the DEG lists were determined by a q-value based FDR threshold of 0.05 after filtering out genes with logfc<0.25 and less than 10% detection in both cases and controls. RCA Gene List is the list of genes ranked by their individual ability to distinguish cases from controls in all datasets. Relative Classification Accuracy is the normalized AUC of individual genes in their ability to distinguish diagnosis status in each dataset. Signed -log10(p-value)s were from comparisons of logistic regression models on disease status with and without each gene (see Methods for more details). The datasets with NA for mean AUC have insufficient cells for at least one of the major cell types leading to inability to create reliable UCell scores for those datasets.

**Supplementary Figure 2.**
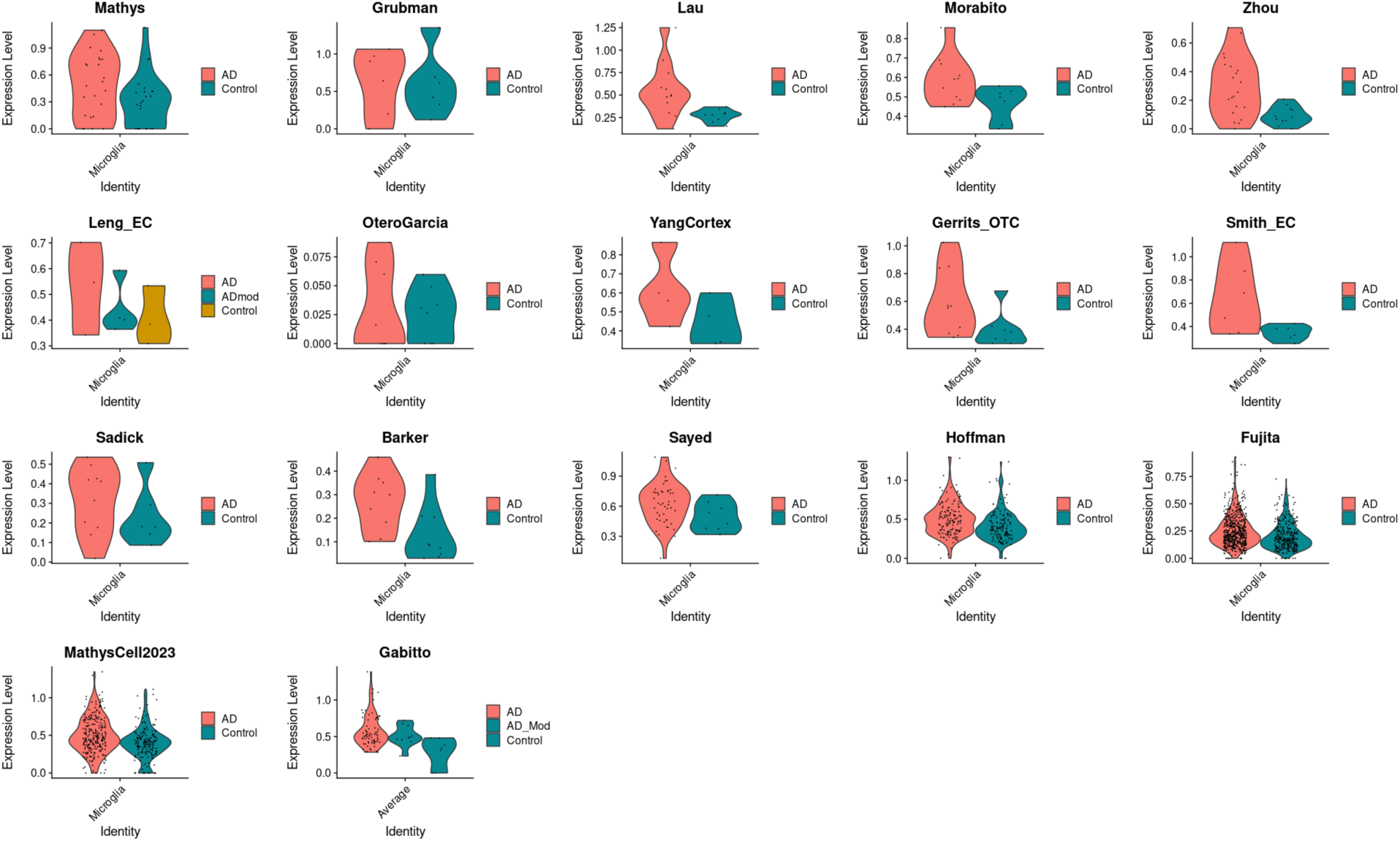
Violin plots of expression of the *RASGRP3* gene in microglia of AD datasets.

**Supplementary Table 16.** Reproducibility metrics with different conditions. The following covariates were regressed out if they were present in the metadata for the dataset: sex, age, PMI, RIN, education level, ethnicity, language, age at death, batch, fixation interval, nCount_RNA, and nFeature_RNA. For all analyses here the DEG lists included the same number of top genes (based on the 814 SumRank genes with -log10(p-value)>3.65). Individual Gene AUC List is the list of genes ranked by their individual ability to distinguish cases from controls in all datasets. Relative Classification Accuracy is the normalized AUC of individual genes in their ability to distinguish diagnosis status in each dataset. Signed -log10(p-value)s were from comparisons of logistic regression models on disease status with and without each gene (see Methods for more details). The Barker dataset was removed from the linear regression analysis due to its focus on individuals with similar Braak scores but differing cognitive impairment.

| Analysis Method | Mean AUC when using DEGs as a Group to Predict Diagnoses of Left-Out Datasets | Specificity: Percentage of DEGs in Top 10% of RCA Gene List | Mean Relative Classification Accuracy of Individual DEGs | Mean abs(log2fc) of individual genes in comparisons of cases vs. controls in each dataset | Mean Negative log10 p-value of individual genes in logistic regressions of diagnosis status in each dataset |
| --- | --- | --- | --- | --- | --- |
| Original: DESeq2 without regressing out covariates in all 21 datasets | 0.784 | 73 | 64.4 | 0.33 | 1.16 |
| DESeq2 regressing out covariates in all 21 datasets | 0.771 | 70 | 65.5 | 0.36 | 1.17 |
| DESeq2 without regressing out covariates in 11 datasets of 10+ cases | 0.778 | 70 | 64.3 | 0.31 | 1.21 |
| DESeq2 regressing out covariates in 11 datasets of 10+ cases | 0.793 | 66 | 65.0 | 0.34 | 1.16 |
| Logistic Regression regressing out covariates in all 21 datasets | 0.761 | 78 | 68.6 | 0.28 | 1.23 |
| Linear Regression on Braak Score regressing out covariates in all datasets (except Barker dataset) | 0.759 | 76 | 66.9 | 0.25 | 1.21 |

**Supplementary Table 17.** Reproducibility metrics comparing Memento on single-cell level data to DESeq2 on pseudobulk data.

| Disease | Gene Set Type | Mean AUC when using DEGs as a Group to Predict Diagnoses of Left-Out Datasets | Specificity: Percentage of DEGs in Top 10% of RCA Gene List | Mean Relative Classification Accuracy of Individual DEGs | Mean absolute log2fc of individual genes between cases and controls in each dataset |
| --- | --- | --- | --- | --- | --- |
| PD | Pseudobulk+DESeq2: Mean of Individual Datasets | 0.77 | 57 | 53.0 | 0.31 |
| PD | Pseudobulk+DESeq2: SumRank | 0.88 | 87 | 71.0 | 0.52 |
| PD | Memento: Mean of Individual Datasets | 0.84 | 33 | 40.4 | 0.16 |
| PD | Memento: SumRank | 0.87 | 60 | 70.6 | 0.52 |

**Supplementary Table 18.** Reproducibility metrics when all AD datasets are subsetted to 6 cases and 6 controls each (Leng_EC and YangCortex are not present due to not having sufficient sample size). For all analyses here the DEG lists included the same number of top genes (based on the 814 SumRank genes with -log10(p-value)>3.65). Individual Gene AUC List is the list of genes ranked by their individual ability to distinguish cases from controls in all datasets. Relative Classification Accuracy is the normalized AUC of individual genes in their ability to distinguish diagnosis status in each dataset. Signed -log10(p-value)s were from comparisons of logistic regression models on disease status with and without each gene (see Methods for more details).

| Dataset | Mean AUC when using DEGs as a Group to Predict Diagnoses of Left-Out Datasets | Specificity: Percentage of DEGs in Top 10% of RCA Gene List | Mean Relative Classification Accuracy of Individual DEGs | Mean abs(log2fc) of individual genes in comparisons of cases vs. controls in each dataset | Mean Negative log10 p-value of individual genes in logistic regressions of diagnosis status in each dataset |
| --- | --- | --- | --- | --- | --- |
| Mathys | 0.70 | 20 | 30.4 | 0.02 | 0.05 |
| Grubman | 0.64 | 36 | 41.3 | 0.13 | 0.25 |
| Lau | 0.71 | 28 | 37.8 | 0.11 | 0.28 |
| Morabito | 0.67 | 43 | 51.9 | 0.15 | 0.51 |
| Zhou | 0.68 | 41 | 46.2 | 0.13 | 0.38 |
| OteroGarcia | 0.55 | 24 | 31.3 | 0.02 | 0.06 |
| Gerrits OTC | 0.60 | 29 | 40.2 | 0.11 | 0.31 |
| Smith EC | 0.66 | 29 | 38.5 | 0.13 | 0.29 |
| Sadick | 0.72 | 32 | 43.1 | 0.13 | 0.37 |
| Barker | 0.69 | 35 | 45.9 | 0.10 | 0.29 |
| Sayed | 0.65 | 35 | 44.4 | 0.11 | 0.35 |
| Hoffman | 0.65 | 11 | 23.8 | -0.02 | -0.01 |
| Fujita | 0.64 | 11 | 24.0 | 0.00 | -0.01 |
| MathysCell | 0.68 | 25 | 36.2 | 0.08 | 0.22 |
| Gabbito | 0.86 | 32 | 39.0 | 0.13 | 0.43 |

**Supplementary Table 19.**
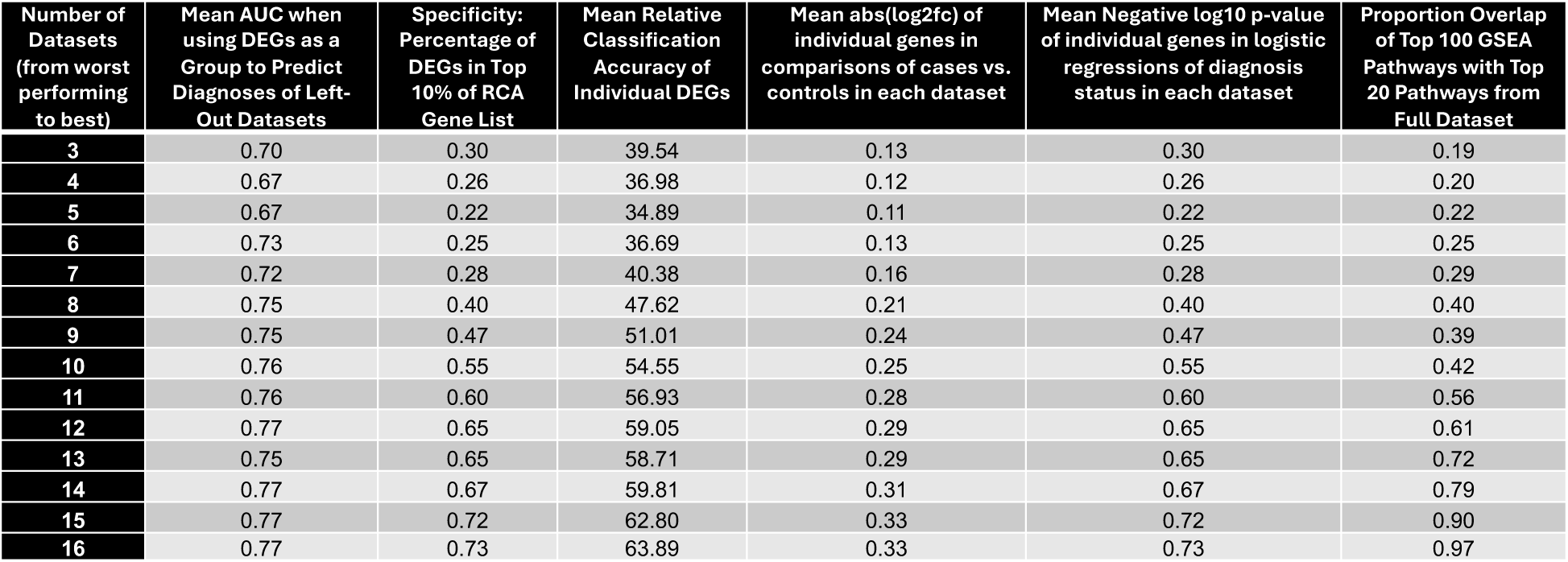
Reproducibility metrics of SumRank meta-analysis DEGs when AD datasets successively added from datasets with lowest AUC to datasets with highest AUC. For all analyses here the DEG lists included the same number of top genes (based on the 814 SumRank genes with -log10(p-value)>3.65). Individual Gene AUC List is the list of genes ranked by their individual ability to distinguish cases from controls in all datasets. Relative Classification Accuracy is the normalized AUC of individual genes in their ability to distinguish diagnosis status in each dataset. Signed -log10(p-value)s were from comparisons of logistic regression models on disease status with and without each gene. Proportion overlap of GSEA pathways is the proportion of the top 20 GSEA pathways from the full dataset that are present in the top 100 GSEA pathways from SumRank genes for each combination of datasets (see Methods for more details).

**Supplementary Table 20.** Reproducibility metrics of SumRank meta-analysis DEGs when AD datasets successively added from datasets with highest AUC to datasets with lowest AUC. For all analyses here the DEG lists included the same number of top genes (based on the 814 SumRank genes with -log10(p-value)>3.65). Individual Gene AUC List is the list of genes ranked by their individual ability to distinguish cases from controls in all datasets. Relative Classification Accuracy is the normalized AUC of individual genes in their ability to distinguish diagnosis status in each dataset. Signed -log10(p-value)s were from comparisons of logistic regression models on disease status with and without each gene. Proportion overlap of GSEA pathways is the proportion of the top 20 GSEA pathways from the full dataset that are present in the top 100 GSEA pathways from SumRank genes for each combination of datasets (see Methods for more details).

| Number of Datasets (from best performing to worst) | Mean AUC when using DEGs as a Group to Predict Diagnoses of Left-Out Datasets | Specificity: Percentage of DEGs in Top 10% of RCA Gene List | Mean Relative Classification Accuracy of Individual DEGs | Mean abs(log2fc) of individual genes in comparisons of cases vs. controls in each dataset | Mean Negative log10 p-value of individual genes in logistic regressions of diagnosis status in each dataset | Proportion Overlap of Top 100 GSEA Pathways with Top 20 Pathways from Full Dataset |
| --- | --- | --- | --- | --- | --- | --- |
| 3 | 0.77 | 0.57 | 61.85 | 0.21 | 0.57 | 0.53 |
| 4 | 0.77 | 0.54 | 58.84 | 0.23 | 0.54 | 0.57 |
| 5 | 0.75 | 0.58 | 59.19 | 0.23 | 0.58 | 0.54 |
| 6 | 0.75 | 0.58 | 59.46 | 0.24 | 0.58 | 0.56 |
| 7 | 0.75 | 0.59 | 59.12 | 0.24 | 0.59 | 0.58 |
| 8 | 0.77 | 0.64 | 61.50 | 0.27 | 0.64 | 0.68 |
| 9 | 0.78 | 0.64 | 62.32 | 0.29 | 0.64 | 0.76 |
| 10 | 0.77 | 0.71 | 64.82 | 0.30 | 0.71 | 0.75 |
| 11 | 0.74 | 0.73 | 65.73 | 0.30 | 0.73 | 0.77 |
| 12 | 0.74 | 0.75 | 66.37 | 0.31 | 0.75 | 0.82 |
| 13 | 0.75 | 0.74 | 65.48 | 0.31 | 0.74 | 0.84 |
| 14 | 0.75 | 0.74 | 64.99 | 0.31 | 0.74 | 0.88 |
| 15 | 0.77 | 0.73 | 64.77 | 0.33 | 0.73 | 0.89 |
| 16 | 0.75 | 0.74 | 64.91 | 0.33 | 0.74 | 0.96 |

**Supplementary Figure 3.**
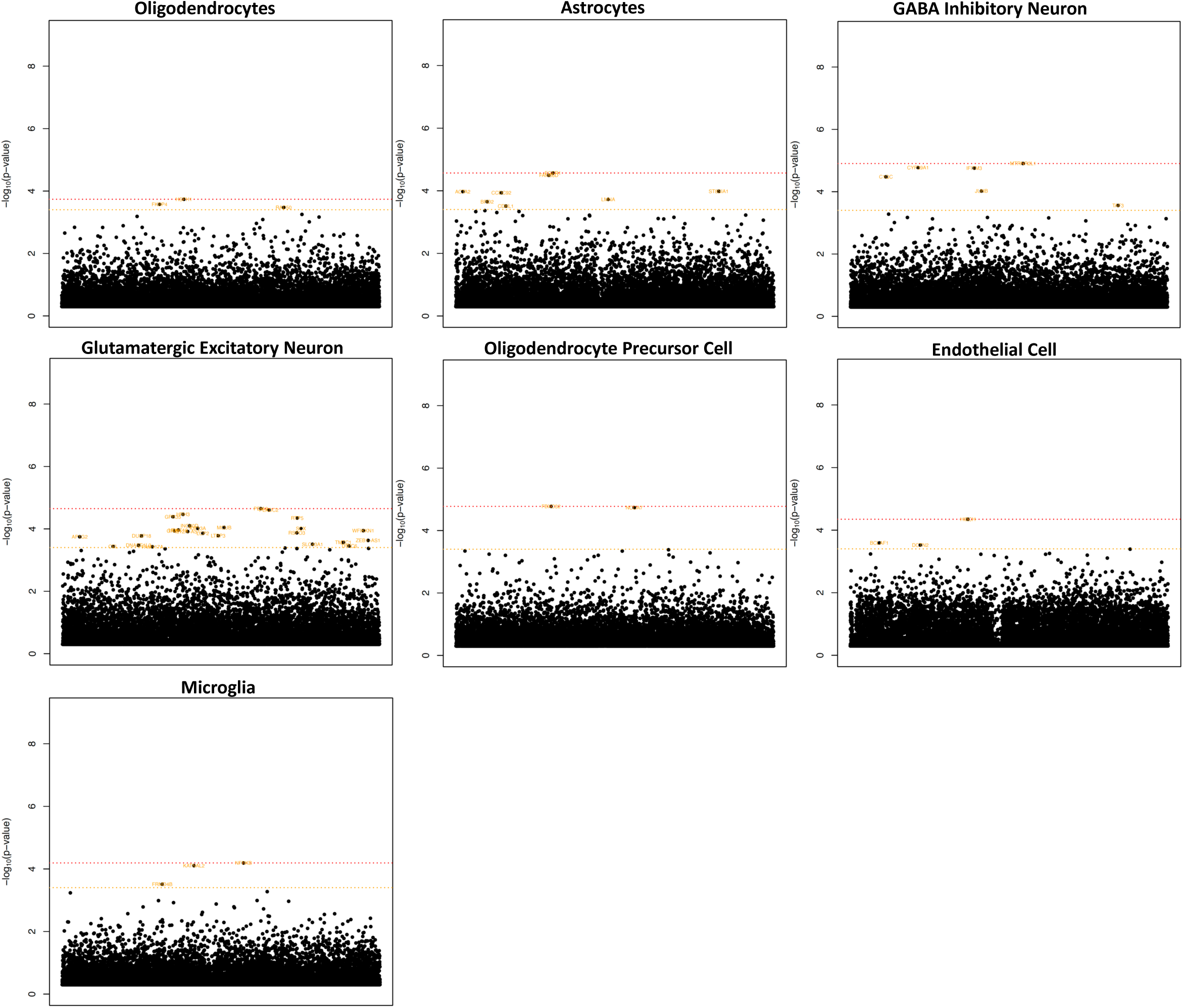
Manhattan plots of up-regulated genes in SCZ. Significance threshold is in red with 0.05 FDR cutoff (Benjamini-Hochberg). In orange is a -log10(p-value) cutoff that maximizes AUC (3.40 for SCZ). The x-axis are genes arranged in alphabetical order. Supplementary Data File 7 provides all genes with their p-values.

**Supplementary Figure 4.**
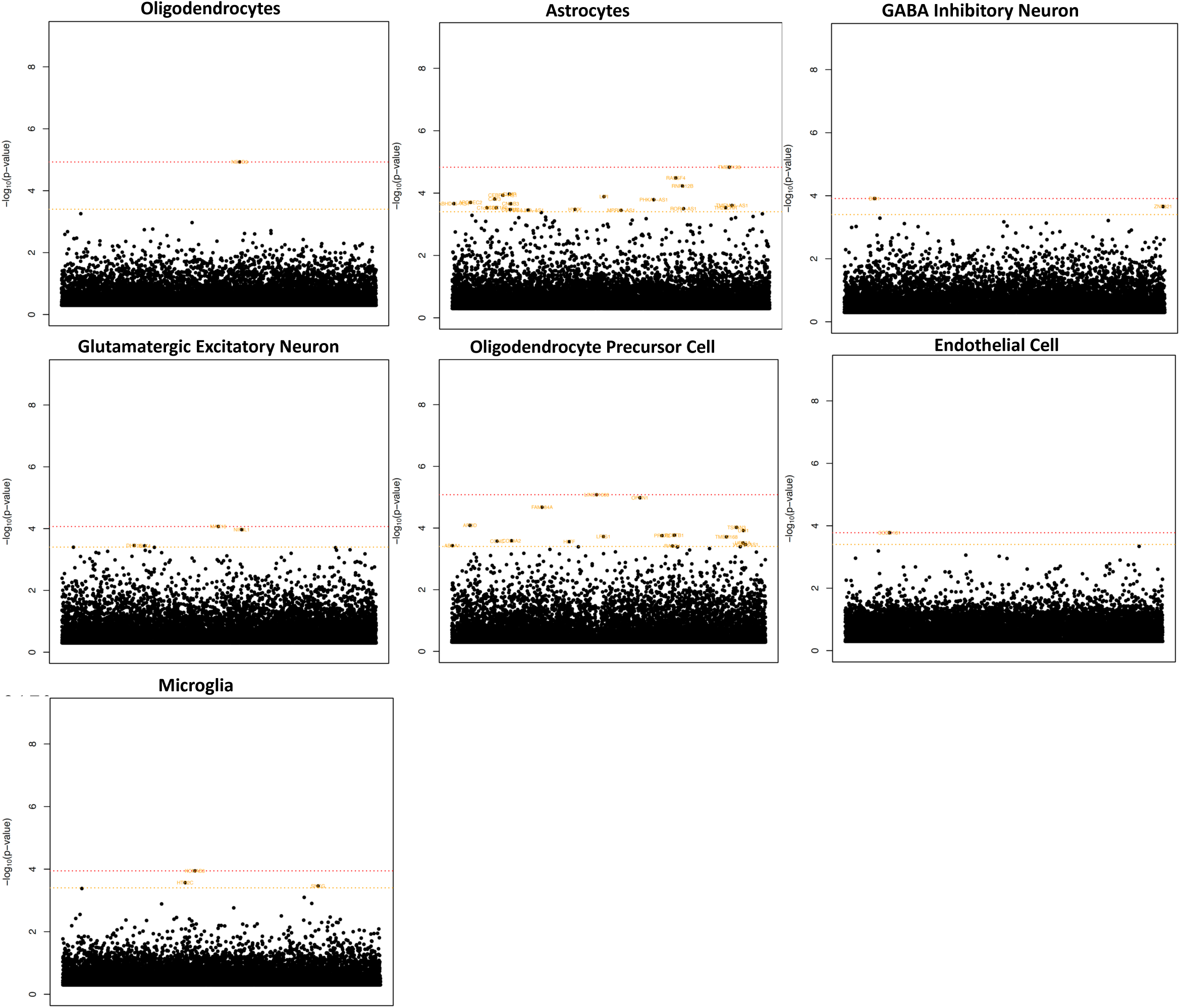
Manhattan plots of down-regulated genes in SCZ. Significance threshold is in red with 0.05 FDR cutoff (Benjamini-Hochberg). In orange is a -log10(p-value) cutoff that maximizes AUC (3.40 for SCZ). The x-axis are genes arranged in alphabetical order. Supplementary Data File 7 provides all genes with their p-values.

**Supplementary Figure 5.**
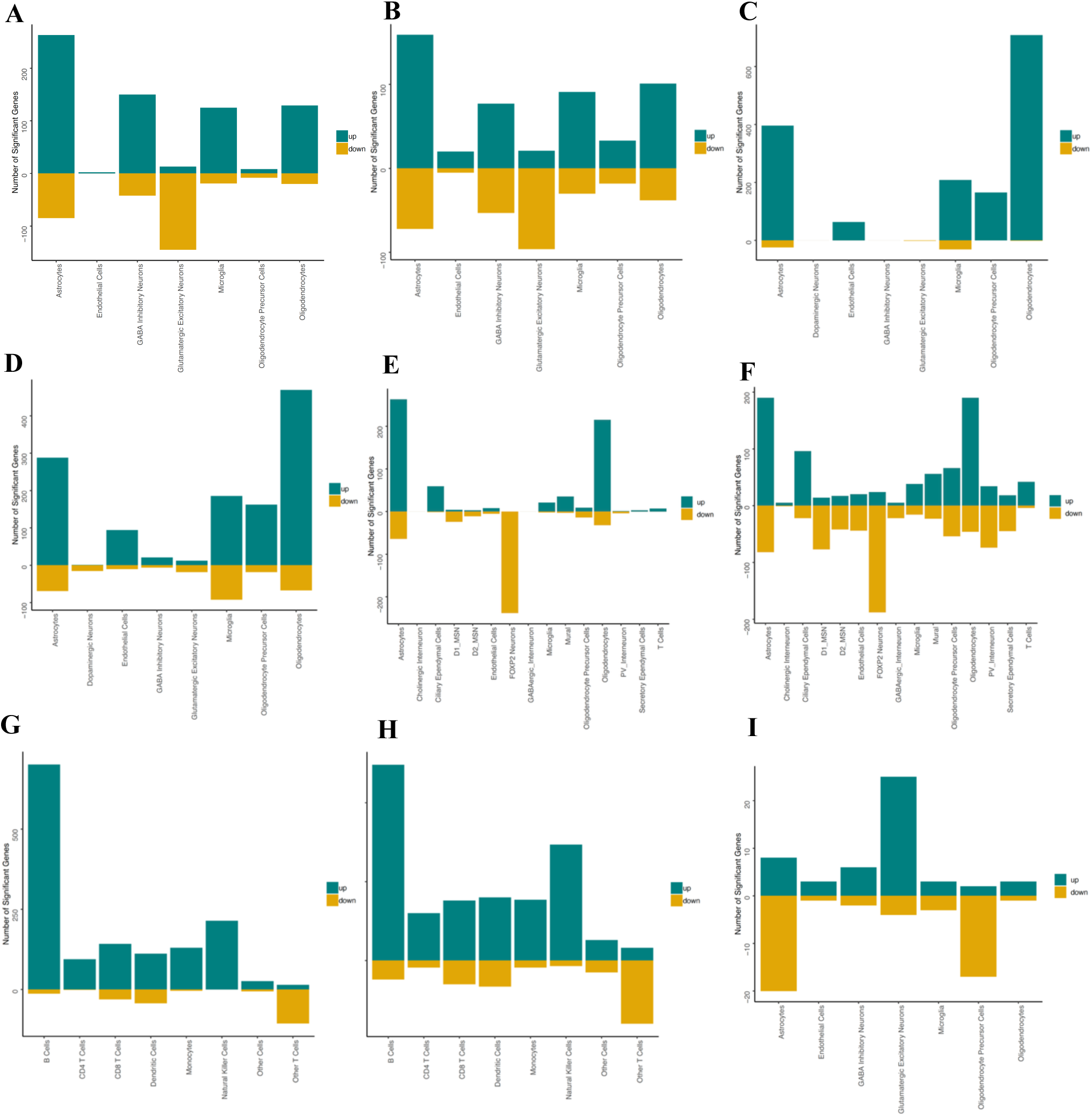
Number of up- and down-regulated genes in AD, PD, COVID-19, and SCZ. A-B) Number of up- and down-regulated genes in AD with a cutoff of 0.05 from Benjamini-Hochberg corrected p-values or a -log10(p-value)>3.65, respectively. **C-D)** Number of up- and down-regulated genes in PD with a cutoff of 0.05 from Benjamini-Hochberg corrected p-values or a -log10(p-value)>3.35, respectively. **E-F)** Number of up- and down-regulated genes in HD with a cutoff of 0.05 from Benjamini-Hochberg corrected p-values or a -log10(p-value)>3.30, respectively. (note: FOXP2 cells from the Lim dataset are not included; see Supplementary Note for details). **G-H)** Number of up- and down-regulated genes in COVID-19 with a cutoff of 0.05 from Benjamini-Hochberg corrected p-values or a -log10(p-value)>3.90, respectively. **I)** Number of up- and down-regulated genes in SCZ with a cutoff -log10(p-value)>3.40. At an FDR cutoff of 0.05 no DEGs are present for SCZ so no plot is shown.

**Supplementary Figure 6.**
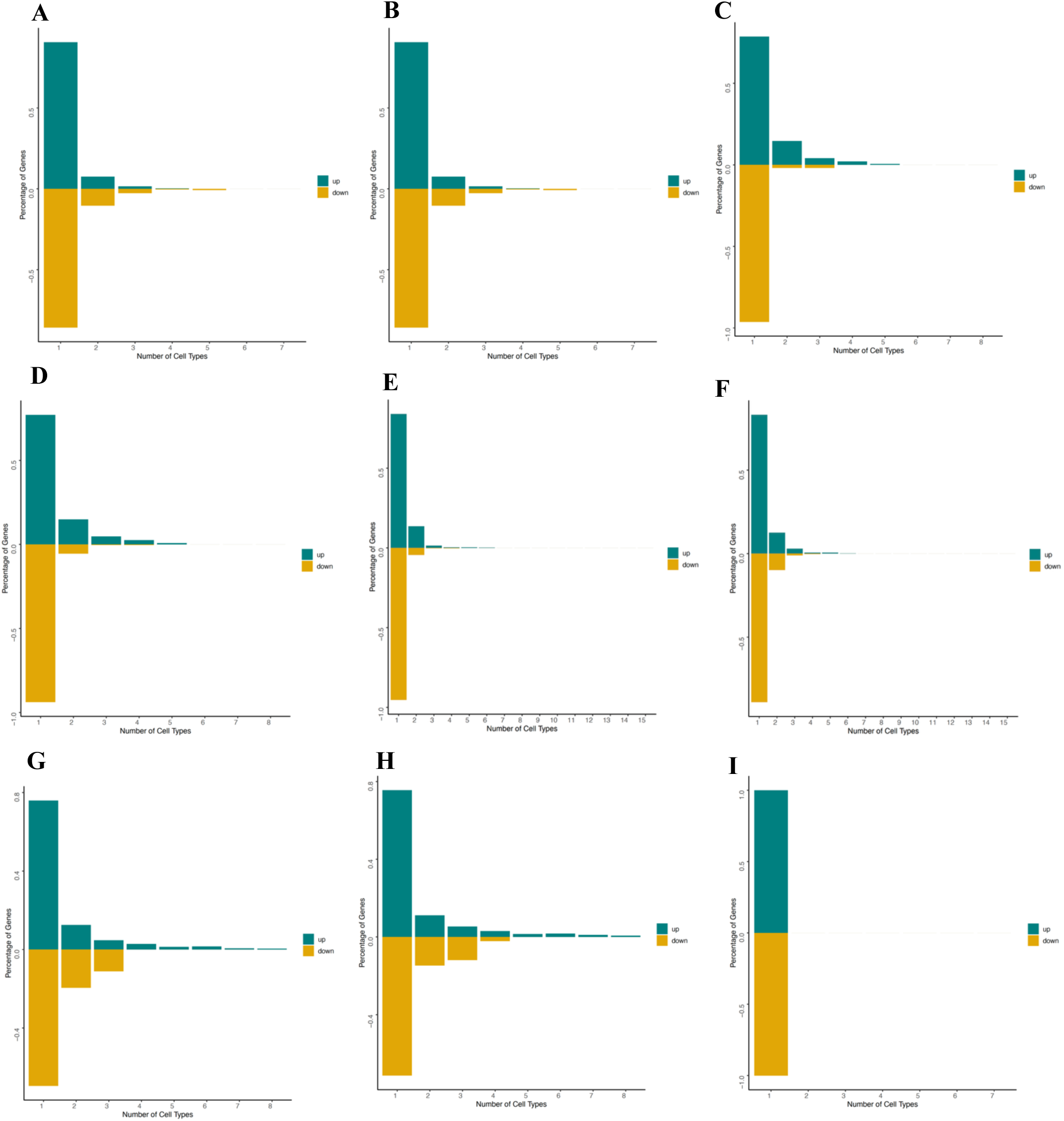
Number of cell types each DEG is present in for AD, PD, HD, COVID-19, and SCZ. A-B) Percentage of genes present in each number of cell types in AD with a cutoff of 0.05 from Benjamini-Hochberg corrected p-values or a -log10(p-value)>3.65, respectively. **C-D)** Percentage of genes present in each number of cell types in PD with a cutoff of 0.05 from Benjamini-Hochberg corrected p-values or a -log10(p-value)>3.35, respectively. **E-F)** Percentage of genes present in each number of cell types in HD with a cutoff of 0.05 from Benjamini-Hochberg corrected p-values or a -log10(p-value)>3.30, respectively. **G-H)** Percentage of genes present in each number of cell types in COVID-19 with a cutoff of 0.05 from Benjamini-Hochberg corrected p-values or a -log10(p-value)>3.90, respectively. **I)** Percentage of genes present in each number of cell types in SCZ with a cutoff -log10(p-value)>3.40. At an FDR cutoff of 0.05 no DEGs are present for SCZ so no plot is shown.

**Supplementary Figure 7.**
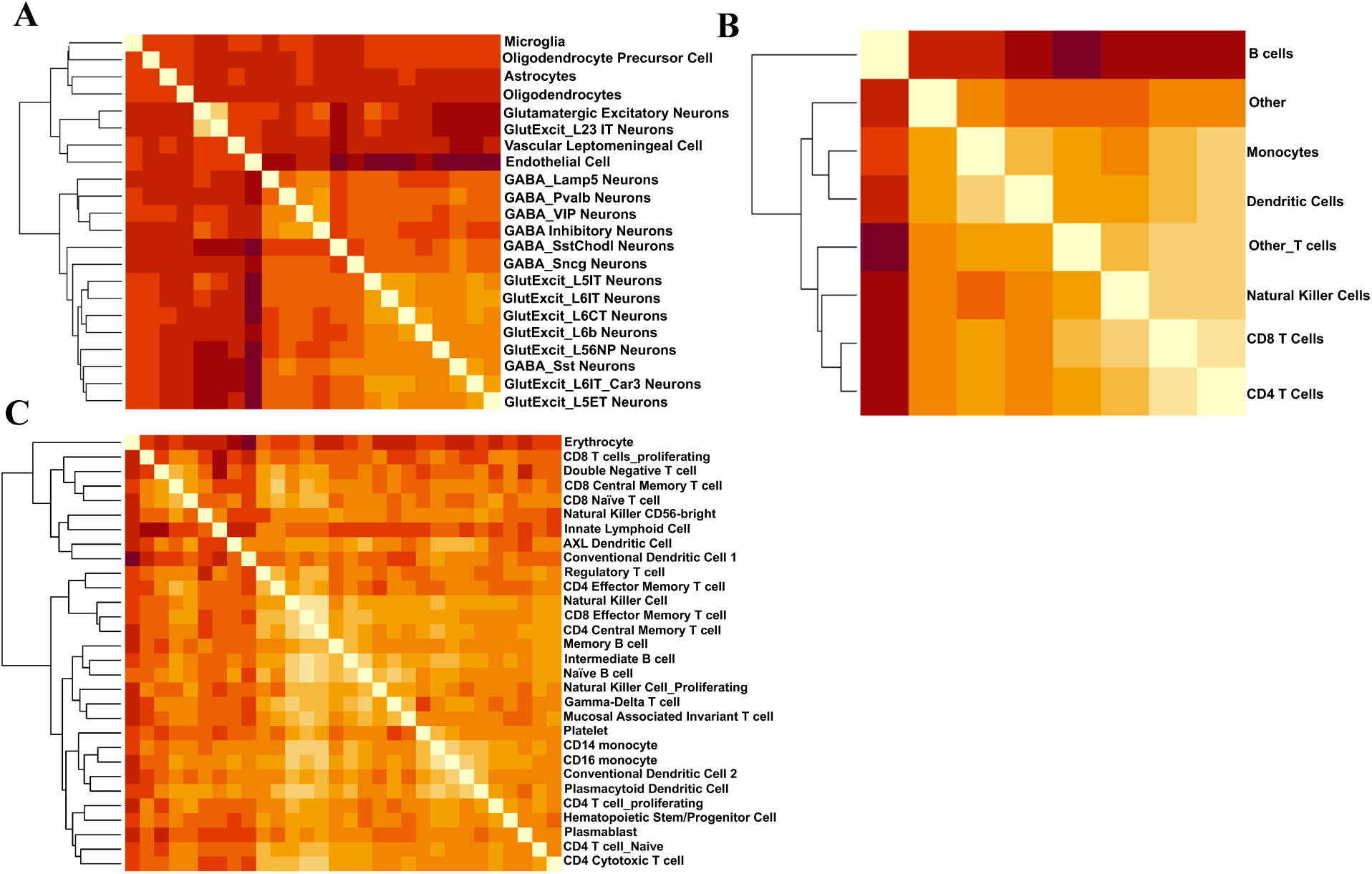
Heatmaps of correlations of UCell scores across cell types. **A)** Correlations in AD at cell type level l2. **B)** Correlations in COVID-19 at cell type level l1. **C)** Correlations in COVID-19 UCell scores at cell type level l2.

**Supplementary Figure 8.**
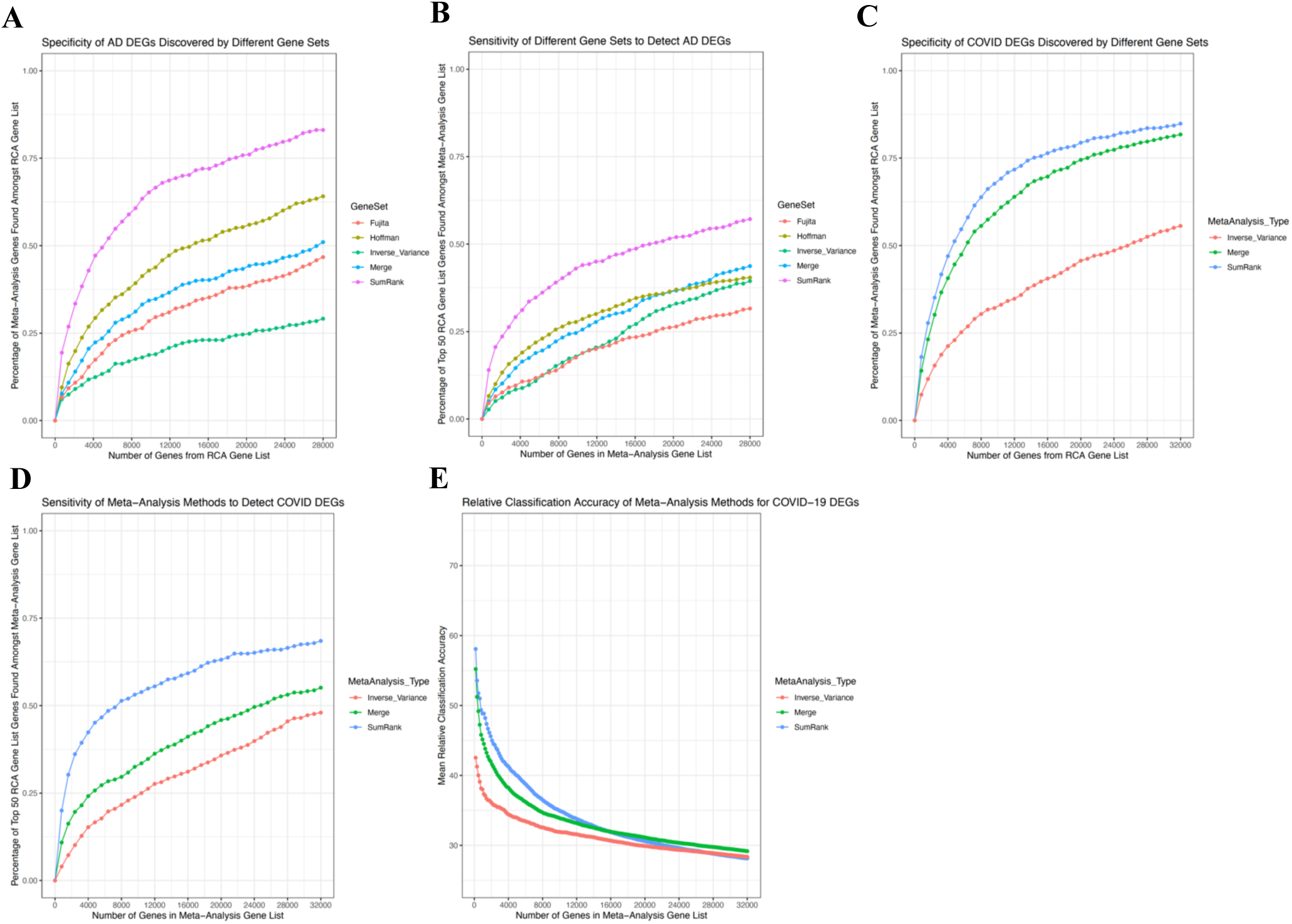
Comparisons of AD and COVID-19 gene sets discovered by different meta-analysis methods. AD DEGs are compared based on their **A)** specificity, as measured by the percentage of their genes that intersect with the RCA Gene List (at different thresholds), and **B)** specificity, as measured by the percentage of the top 50 RCA Gene List genes found in the meta-analysis DEG list at different thresholds. Results are taken across all cell types. The same analyses are shown for COVID-19 in **C)** and **D)**. **E)** Relative Classification Accuracy, the mean AUC of individual genes in their ability to distinguish diagnosis status in each dataset (averaged over all genes in the gene set). The number of genes for A-E are spread evenly across up and down-regulated and all the different cell types.

**Supplementary Figure 9.**
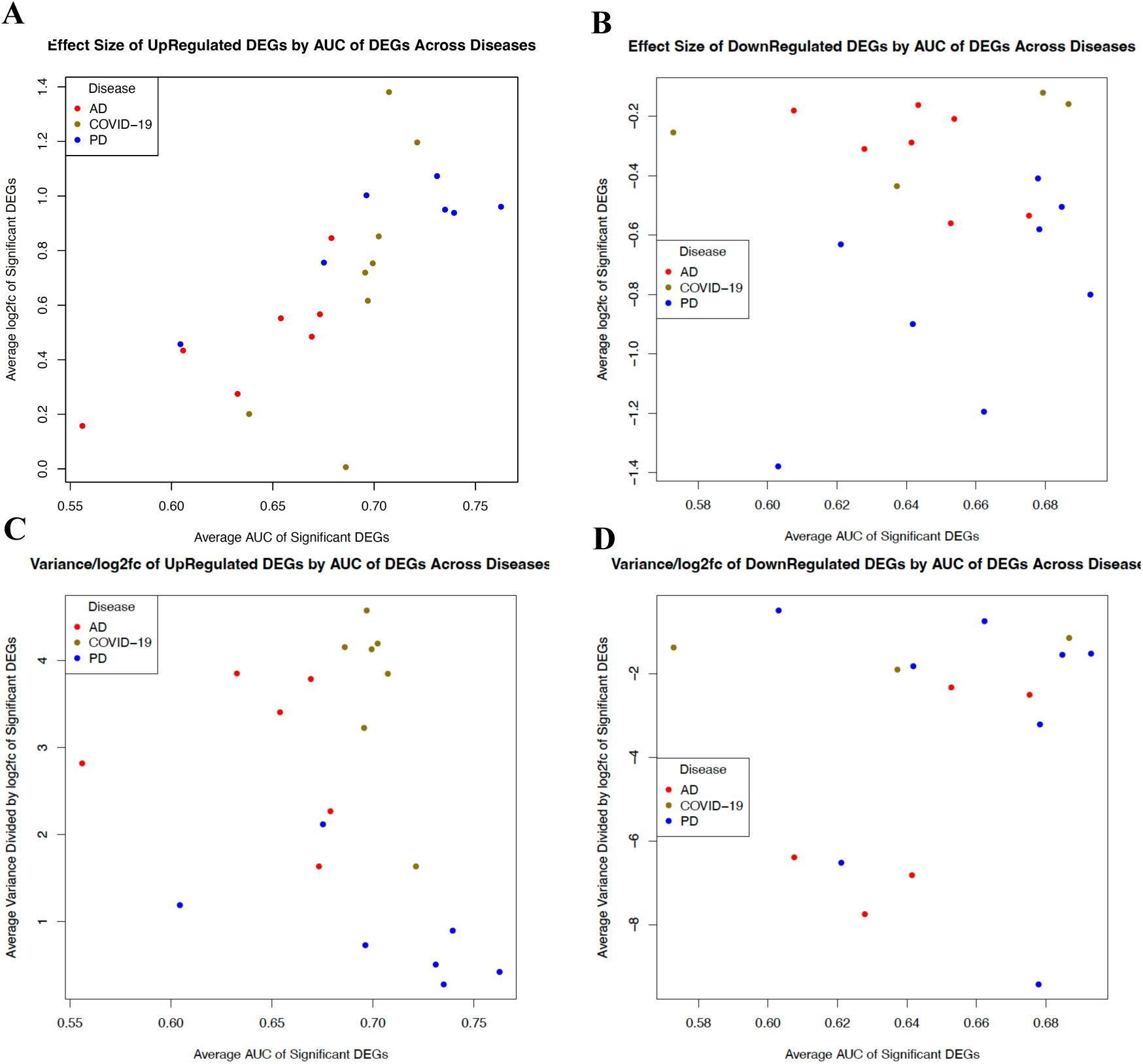
Average reproducibility of genes vs effect size and variance within each cell type for AD, PD, and COVID-19. The average AUC of significant DEGs in each cell type is plotted against their average log2fc for **A)** up-regulated and **B)** down-regulated genes. The average AUC of significant DEGs in each cell type is plotted against their average variance/log2fc for **C)** up-regulated and **D)** down-regulated genes AUCs for each DEG are calculated based on their ability to predict case-control status in all datasets.

**Supplementary Figure 10.**
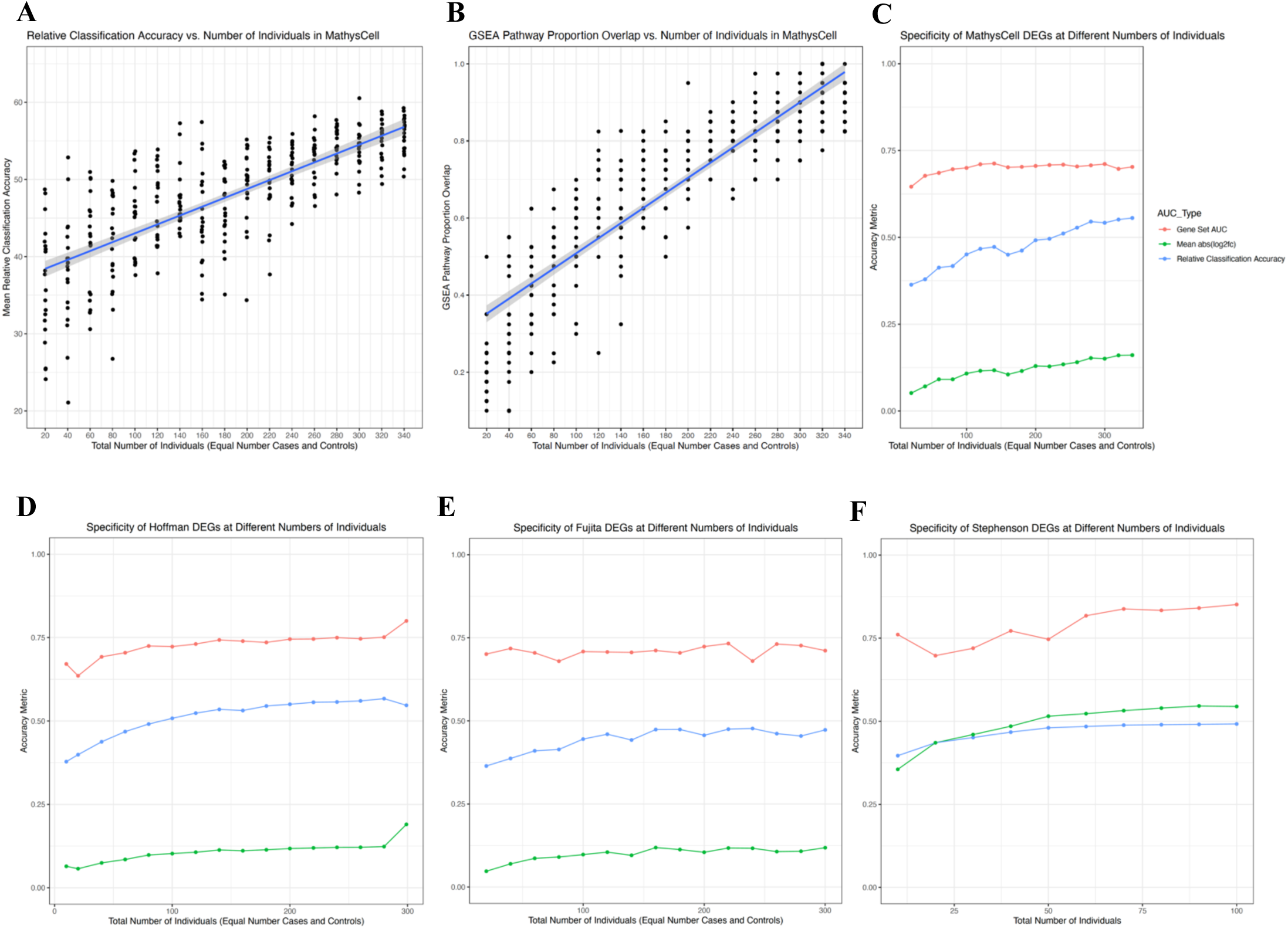
Reproducibility metrics after random down-sampling of large datasets. **A)** Relative Classification Accuracy at different numbers of individuals in MathysCell dataset. **B)** Proportion overlap of GSEA pathways at different number of individuals. Proportion overlap is the proportion of the top 20 GSEA pathways from a subsetted 340 individual MathysCell dataset that overlap with the top 100 GSEA pathways from differential expression after downsampling individuals from the MathysCell dataset (see Methods for more details). **C-F)** Average reproducibility metrics after down-sampling the MathysCell, Hoffman, Fujita, and Stephenson datasets. Gene Set AUC is the mean AUC when using the set of DEGs to predict diagnoses of other datasets. Relative Classification Accuracy is the normalized AUC of individual genes in their ability to distinguish diagnosis status in each dataset. Mean abs(log2fc) were from comparisons of cases vs controls. For all analyses here the DEG lists included the same number of top genes (based on the 814 SumRank genes with -log10(p-value)>3.65). For the Stephenson dataset (E), the points represent cases and controls in the following combinations: ((5,5), (10,10), (15,15), (20,20), (30,20), (40,20), (50,20), (70,20), and (80,20)). All points in B-E are plotted as the mean values after 20 random iterations.

**Supplementary Figure 11.**
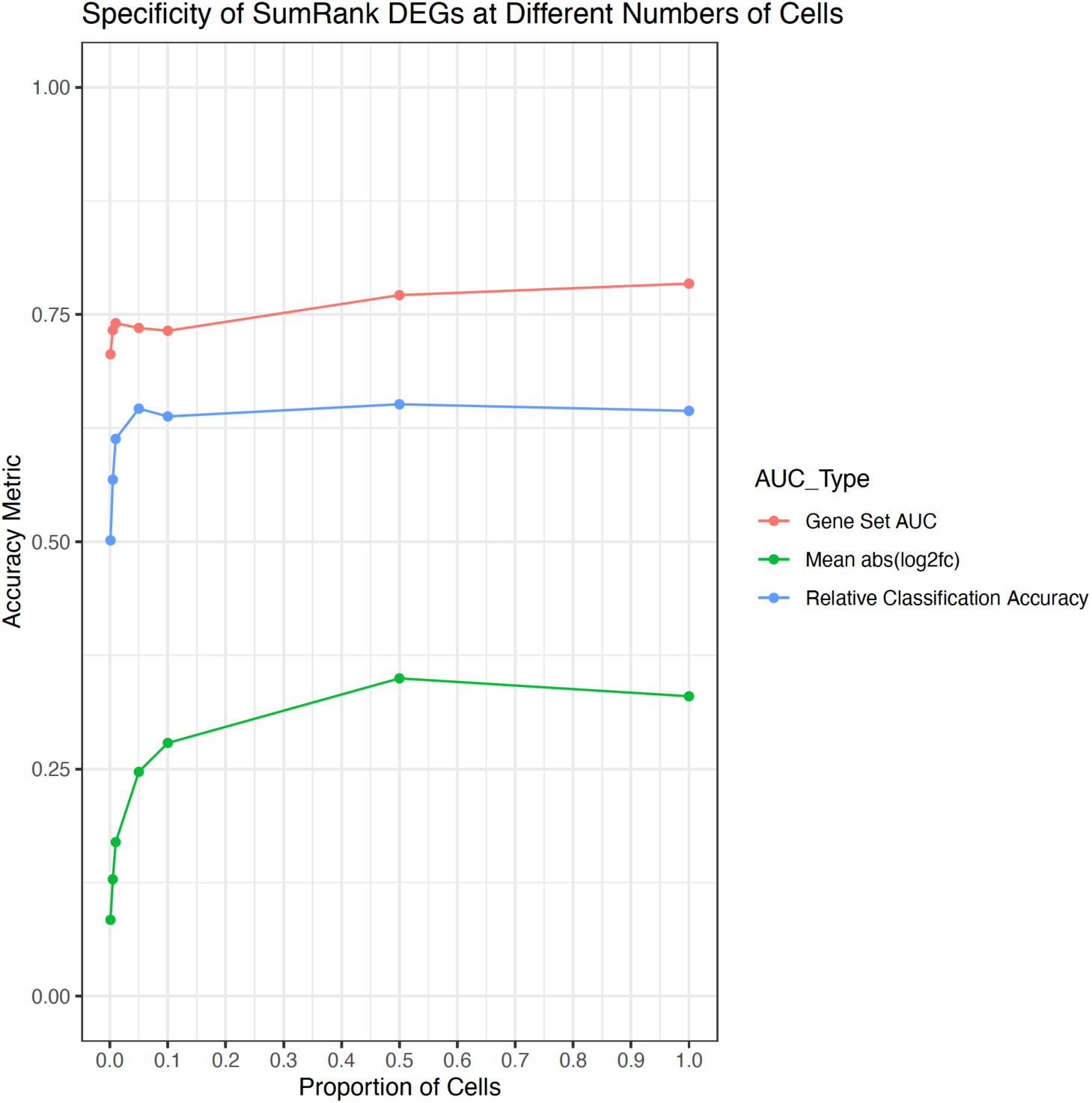
Reproducibility metrics of SumRank AD DEGs after random down-sampling of cells. Gene Set AUC is the mean AUC when using the set of DEGs to predict diagnoses of other datasets. Relative Classification Accuracy is the normalized AUC of individual genes in their ability to distinguish diagnosis status in each dataset. Mean abs(log2fc) were from comparisons of cases vs controls in each dataset. For all analyses here the DEG lists included the same number of top genes (based on the 814 SumRank genes with -log10(p-value)>3.65). The following down-sampling proportions were used: (0.001, 0.005, 0.001, 0.05, 0.1, 0.5).

**Supplementary Figure 12.**
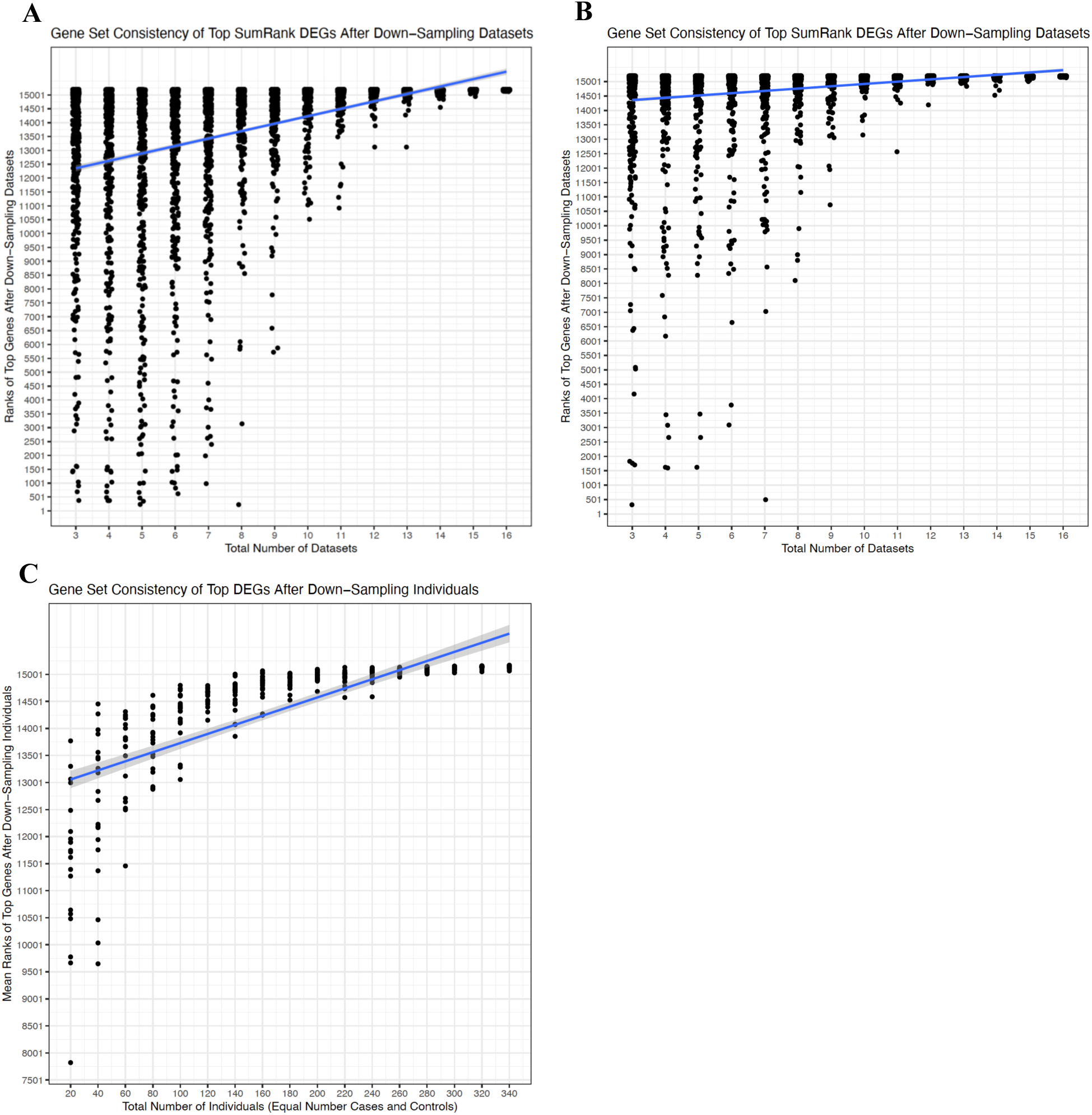
Gene set consistency after random down-sampling of datasets or cells. The top 25 genes from SumRank meta-analysis AD DEGs with all datasets were obtained and their ranks were assessed after performing SumRank after down-sampling datasets. **A)** AD datasets successively added from datasets with the lowest AUC to datasets with highest AUC (compare to Supplementary Table 15). **B)** AD datasets successively added from datasets with the highest AUC to datasets with lowest AUC (compare to Supplementary Table 16). In A-B, ranks are inversely defined with 15201 being the top rank and 1 being the lowest possible rank. Comparisons of all major cell types and both up- and down-regulated are shown here. **C)** The top 25 genes from a subsetted 340 individual MathysCell dataset were obtained and their ranks were assessed after down-sampling individuals from the MathysCell dataset. Each point represents a random sampling where the value is the mean rank of the 25 genes in each major cell type for both up- and down-regulated genes. See Methods for more details.

**Supplementary Figure 13.**
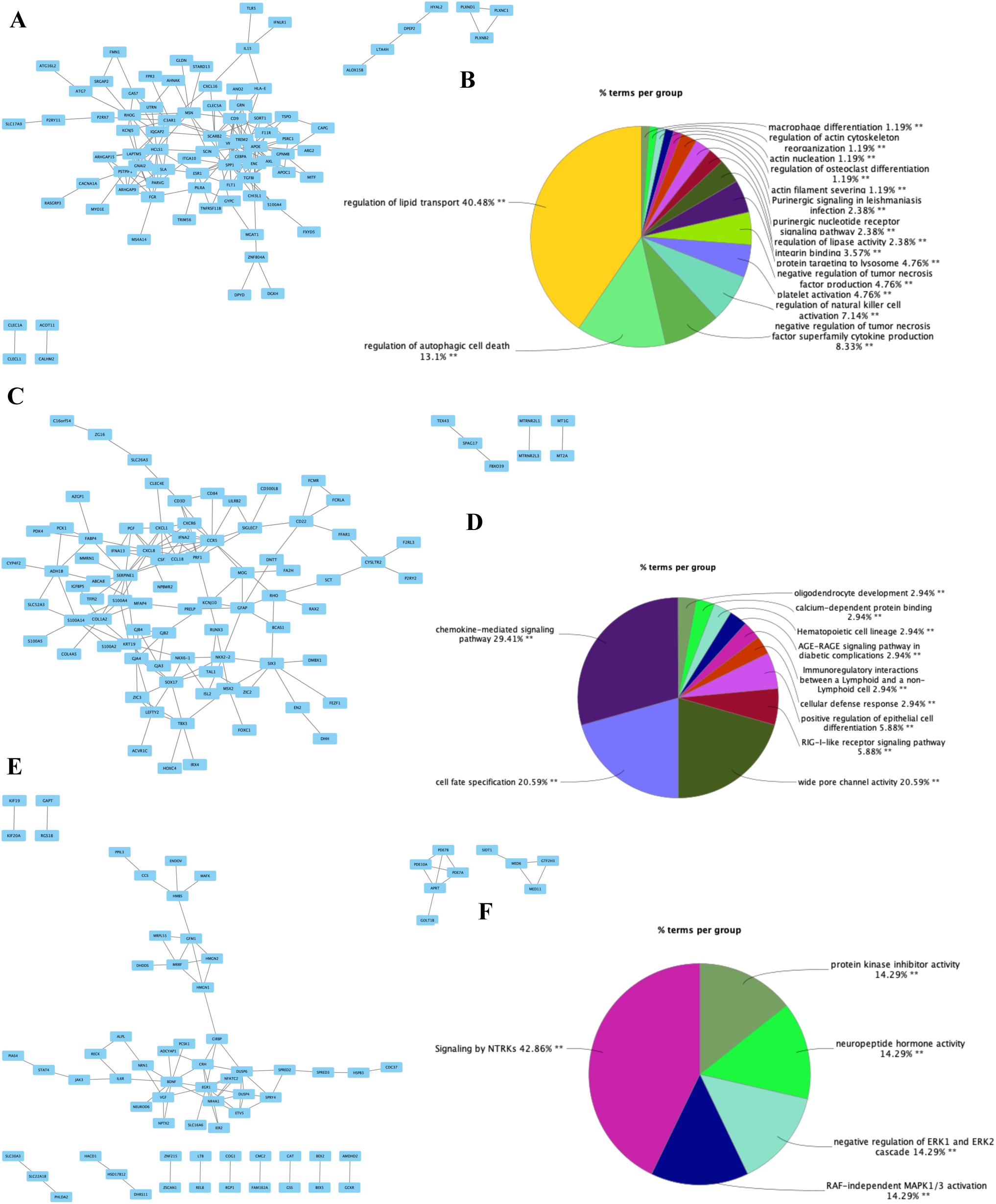
AD DEG protein-protein interaction (PPI) networks. **A)** Microglia up-regulated network formed by STRING^62^ database. **B)** The most densely connected network was analyzed by Cytoscape^115^ and enriched terms are shown in the pie chart. **C-D)** GABAergic neuron up-regulated network and enriched terms. **E-F)** Glutamatergic neuron down-regulated network and enriched terms. See Supplementary Data File 3 for the genes driving the enrichment terms.

**Supplementary Figure 14.**
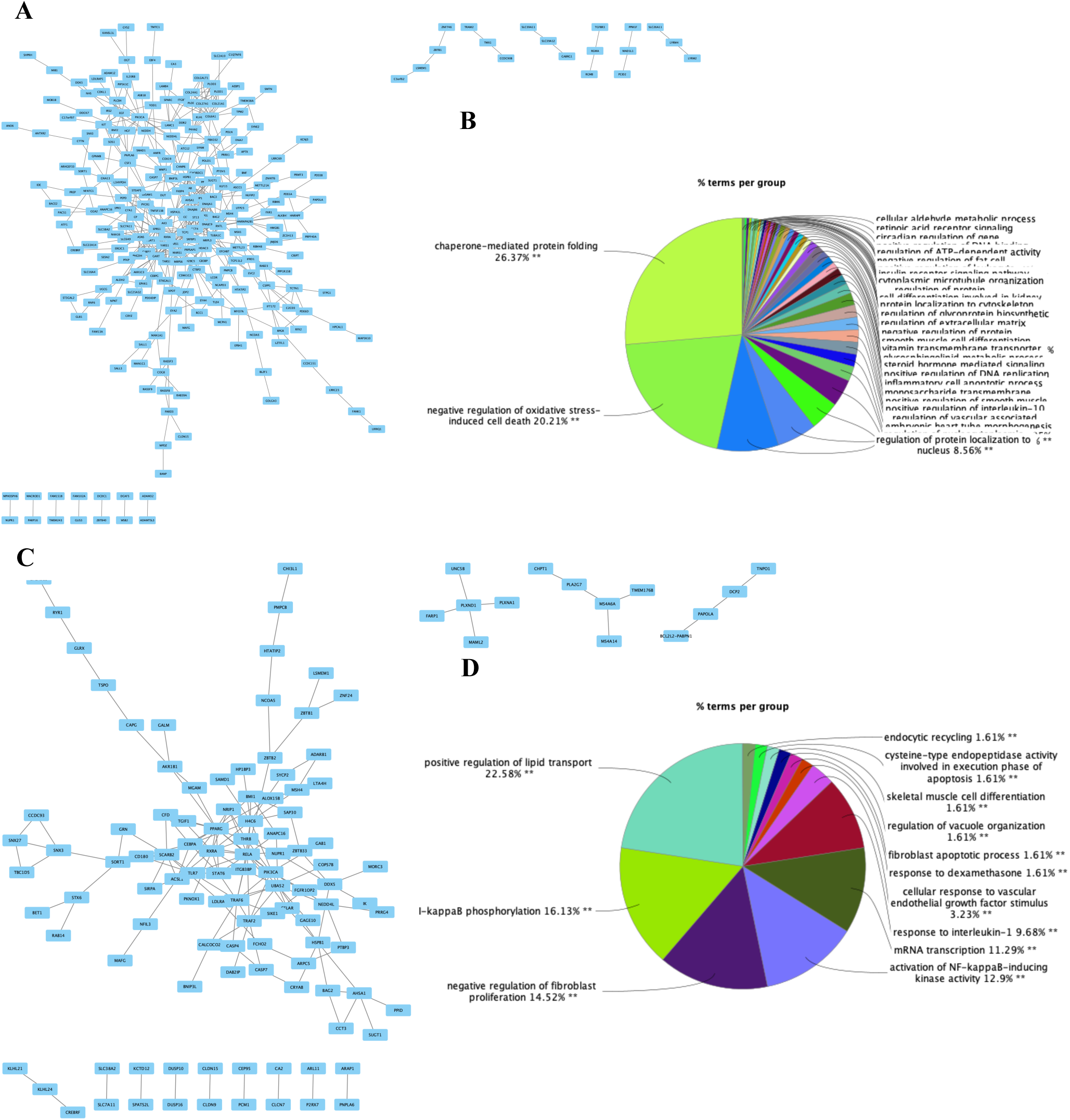
PD DEG protein-protein interaction (PPI) networks. **A)** Astrocyte up-regulated network formed by STRING^62^ database. **B)** The most densely connected network was analyzed by Cytoscape^115^ and enriched terms are shown in the pie chart. **C-D)** Microglia up-regulated network and enriched terms. See Supplementary Data File 4 for the genes driving the enrichment terms.

**Supplementary Table 21.** Genes significant in SumRank meta-analysis that are also significant in human genetic studies. . The -log10(p-value)s listed here are from the SumRank meta-analysis. See Methods for more details of specific human genetic studies used.

| Disease | Gene | Study Type | Cell Type | -log10(p-value) | Up- or Down-Regulated |
| --- | --- | --- | --- | --- | --- |
| AD | ABCA7 | WES | Astrocytes | 4.61 | Up |
| AD | APOE | WES | Microglia | 6.30 | Up |
| AD | PILRA | WES | Microglia | 3.80 | Up |
| AD | TREM2 | WES | Microglia | 4.43 | Up |
| AD | CR1 | GWAS | Oligodendrocytes | 3.67 | Up |
| AD | ABCA7 | GWAS | Astrocytes | 4.61 | Up |
| AD | INPP5D | GWAS | Astrocytes | 5.31 | Up |
| AD | EGFR | GWAS | Oligodendrocyte Precursor Cell | 5.43 | Up |
| AD | MAF | GWAS | Glutamatergic Neurons | 4.45 | Up |
| AD | APOE | GWAS | Microglia | 6.30 | Up |
| AD | GRN | GWAS | Microglia | 4.50 | Up |
| AD | SORT1 | GWAS | Microglia | 5.20 | Up |
| AD | TREM2 | GWAS | Microglia | 4.43 | Up |
| PD | ALG10 | GWAS | Oligodendrocytes | 3.45 | Down |
| PD | BIN3 | GWAS | Oligodendrocytes | 3.63 | Down |
| PD | CCT3 | GWAS | Oligodendrocytes | 3.82 | Down |
| PD | CCT3 | GWAS | Astrocytes | 5.01 | Down |
| PD | PIK3CA | GWAS | Astrocytes | 3.88 | Down |
| PD | CCT3 | GWAS | Oligodendrocyte Precursor Cell | 5.82 | Down |
| PD | MAPT | GWAS | Oligodendrocyte Precursor Cell | 5.80 | Down |
| PD | PIK3CA | GWAS | Oligodendrocyte Precursor Cell | 5.84 | Down |
| PD | CCT3 | GWAS | Microglia | 3.86 | Down |
| PD | PIK3CA | GWAS | Microglia | 3.96 | Down |
| PD | SCARB2 | GWAS | Microglia | 4.14 | Down |

**Supplementary Figure 15.**
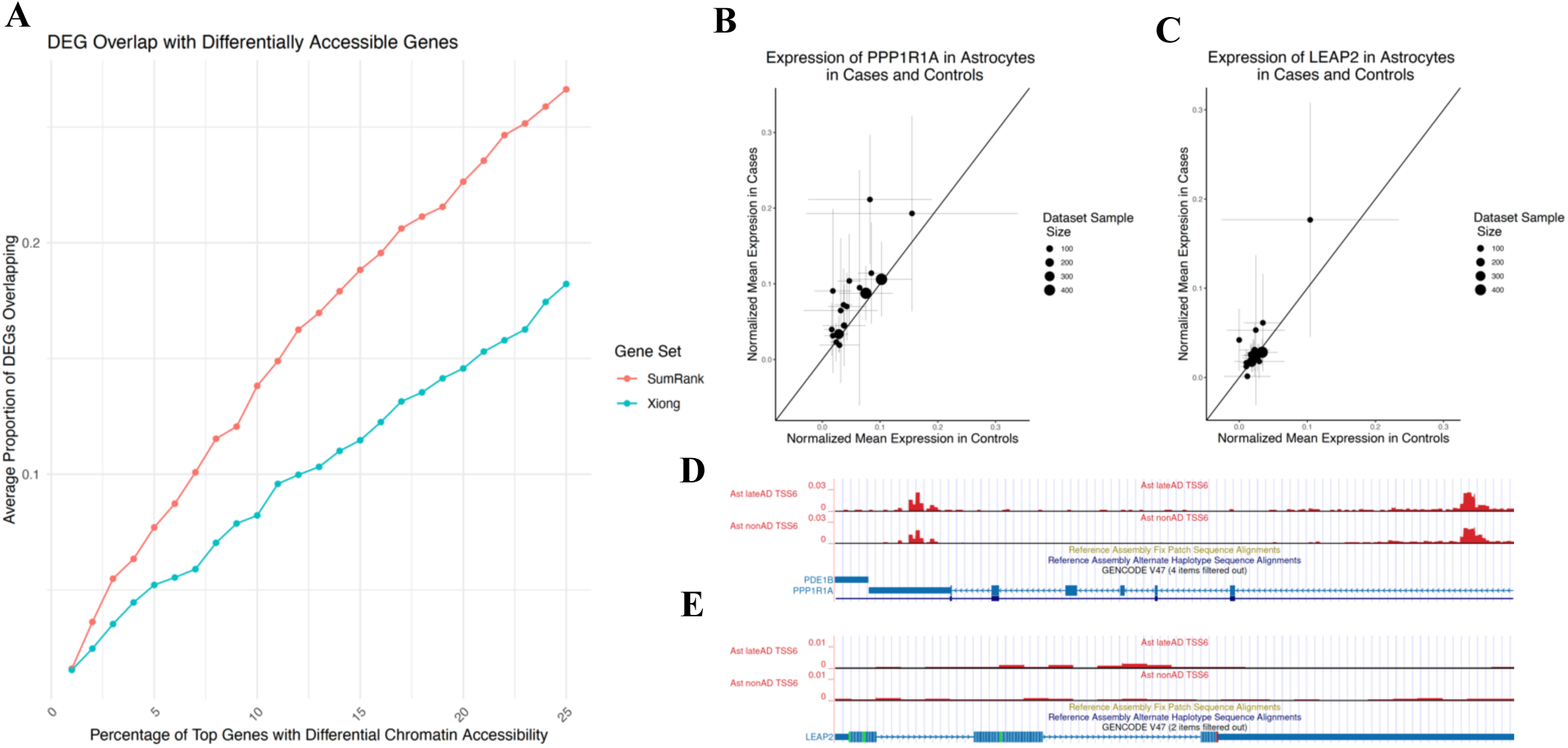
Overlap of meta-analysis DEGs with top genes associated with differential chromatin accessibility from snATAC-seq data. Differential chromatin accessibility peaks were obtained from Xiong et al.^52^ and genes were assigned to peaks based on proximity to the promoter region. Genes were ranked based on p-value for differential accessibility (subsetting to up- or down-regulated). **A)** The overlap of DEGs from SumRank and Xiong snRNA-seq was assessed at different percentages of top genes for all cell types (Methods). **B)** Representative example of a gene (*PPP1R1A*) up-regulated in most datasets in astrocytes (a SumRank DEG) that also shows differential chromatin accessibility. **C)** Representative example of a gene (*LEAP2*) up-regulated in only a few datasets in astrocytes (an inverse variance specific DEG) that does not show differential chromatin accessibility. snATAC-seq peaks visualized on UCSC genome browser for **D)** *PPP1R1A* and **E)** *LEAP2*.

**Supplementary Figure 16.**
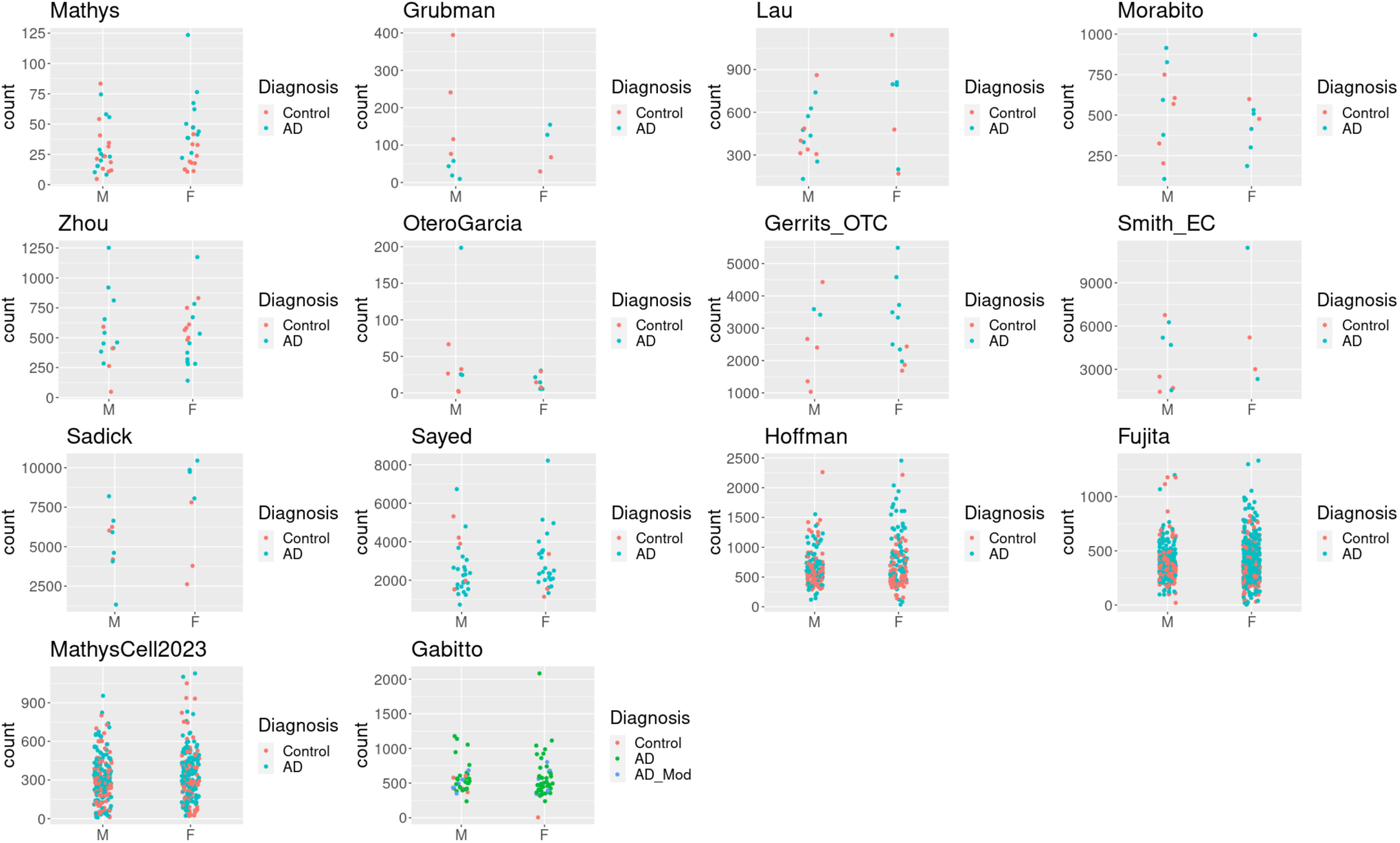
Expression of *ZFP36L1* gene in males and females in astrocytes across different datasets. Each point represents an individual. Analyses performed in DESeq2 (see Methods). M=male; F=female.

**Supplementary Figure 17.**
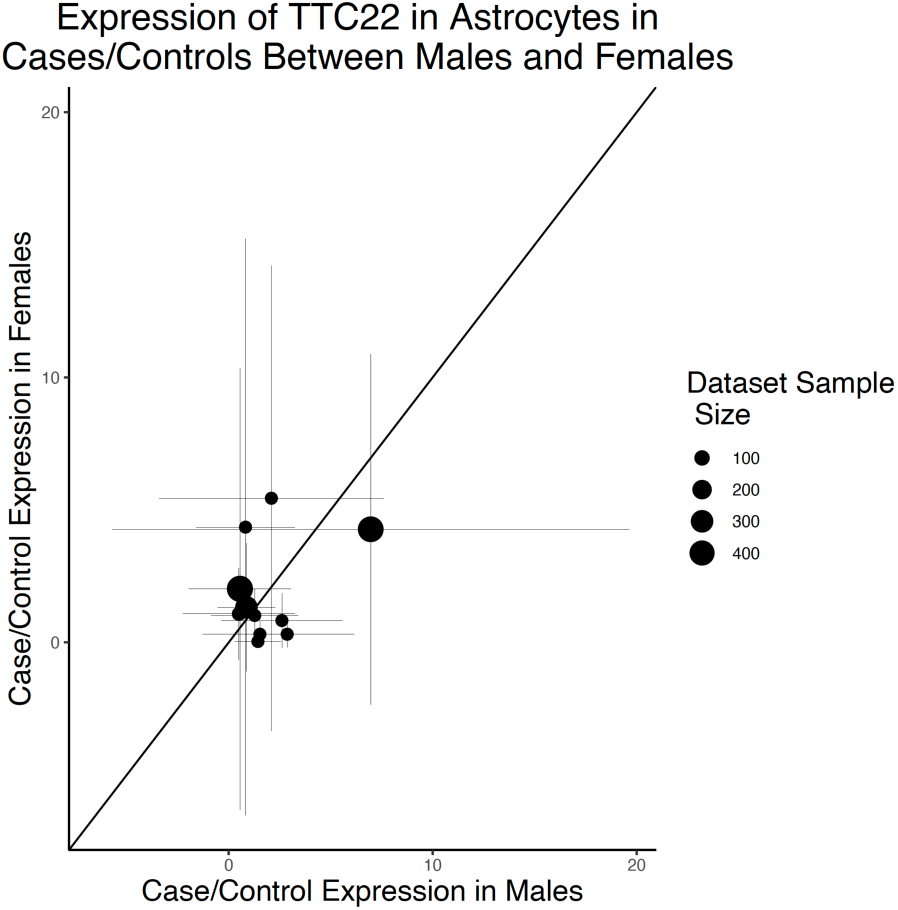
Male and female expression of TTC22 in different AD datasets. The ratios of mean expression of cases over mean expression of controls in females (y-axis) and males (x-axis). Error bars are standard deviations. Values above the line (intercept=0, slope=1) are up-regulated in females more than males, while values below the line are up-regulated in males more than females. This *TTC22* gene was the top gene with putative female-specific expression based on the merge Sex Interaction method with p_val_Bonferroni<5e-13.

**Supplementary Figure 18.**
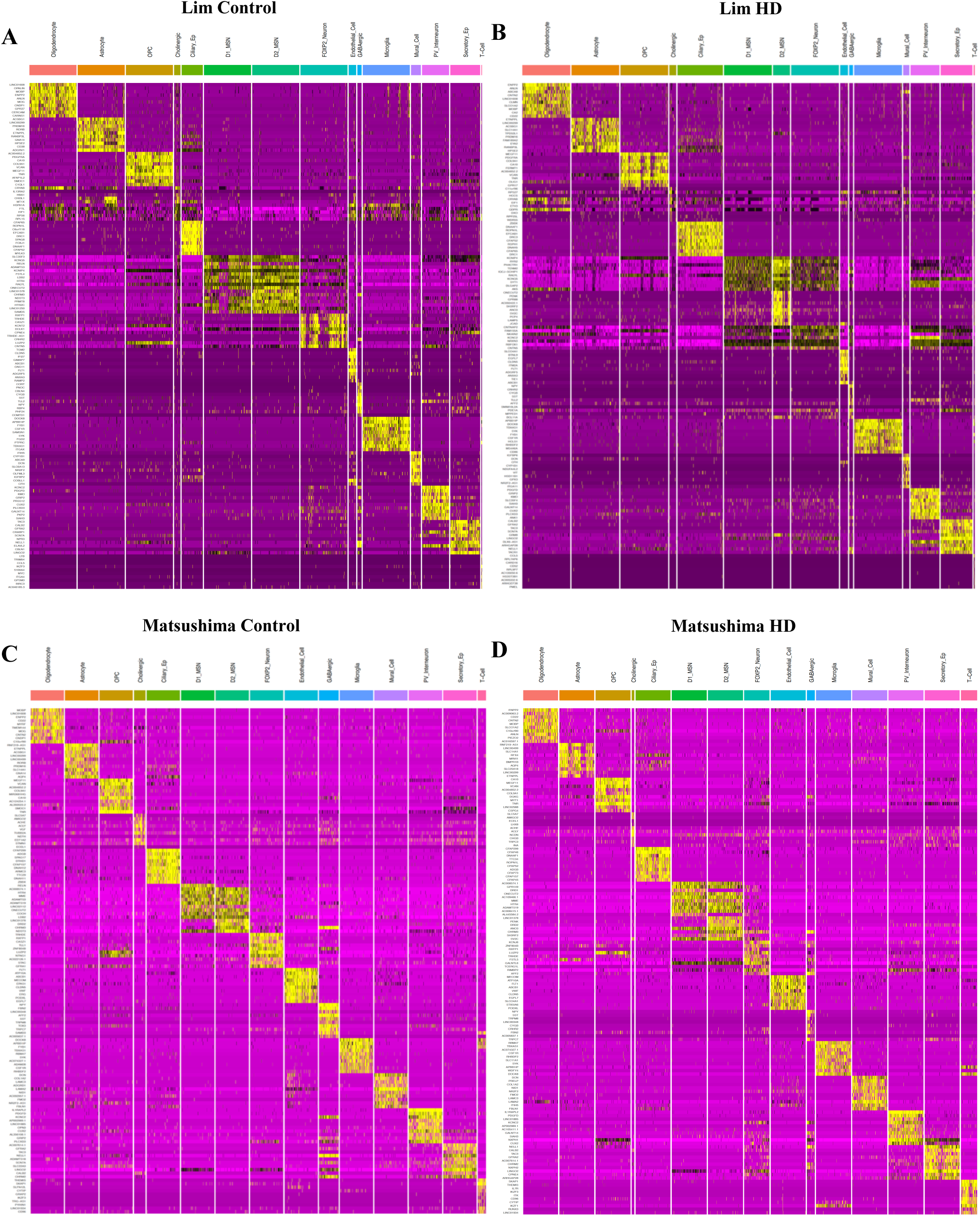
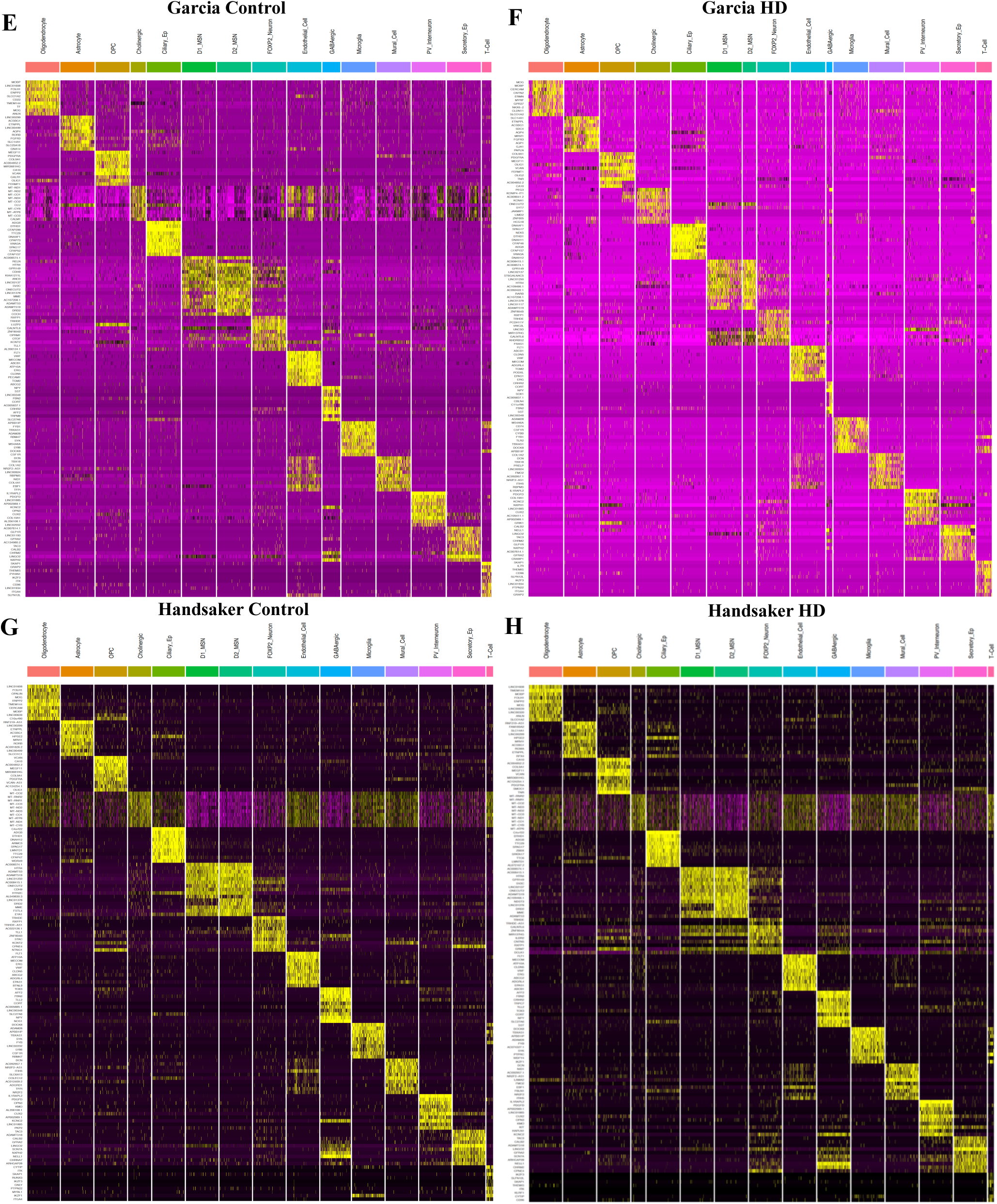
Cell-Type mapping of FoxP2 neurons in HD. Heatmaps of top differentially expressed genes between cells of each cell type relative to all other cells in the **A-B)** Lim, **C-D)** Matsushima, **E-F)** Garcia, and **G-H)** Handsaker datasets for control and HD, respectively.

**Supplementary Figure 19.**
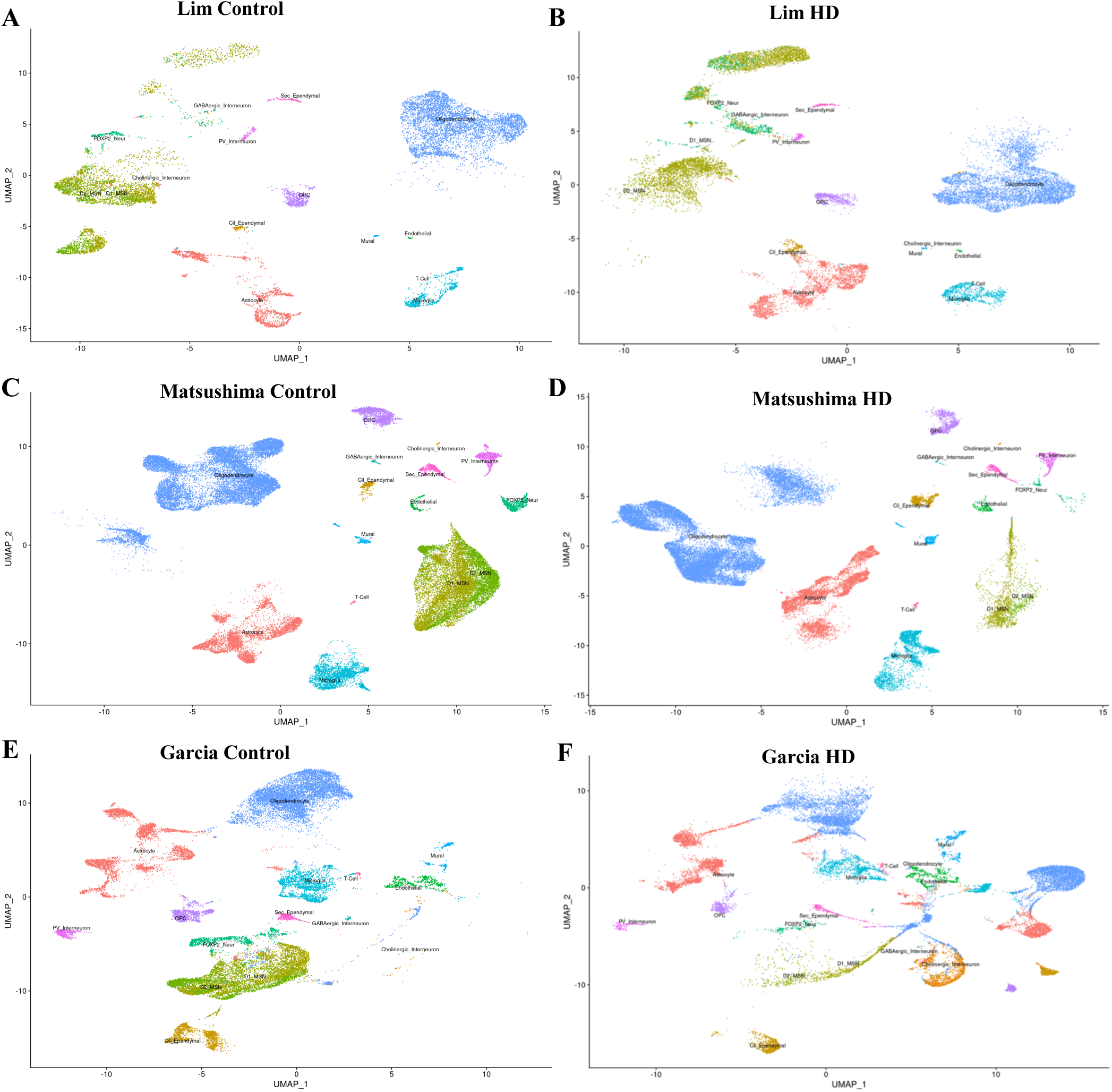
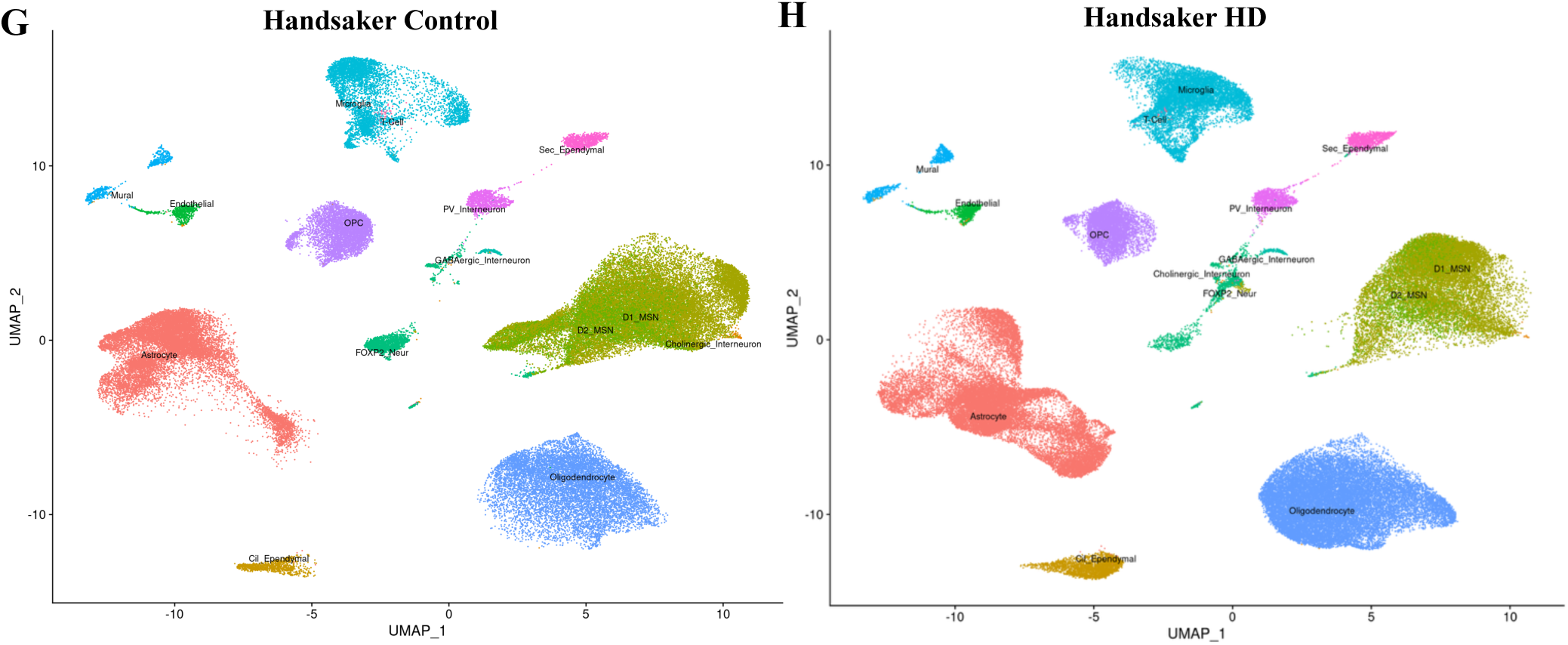
UMAPs for control and HD cells in the A-B) Lim, C-D) Matsushima, E-F) Garcia, and G-H) Handsaker datasets.

**Supplementary Figure 20.**
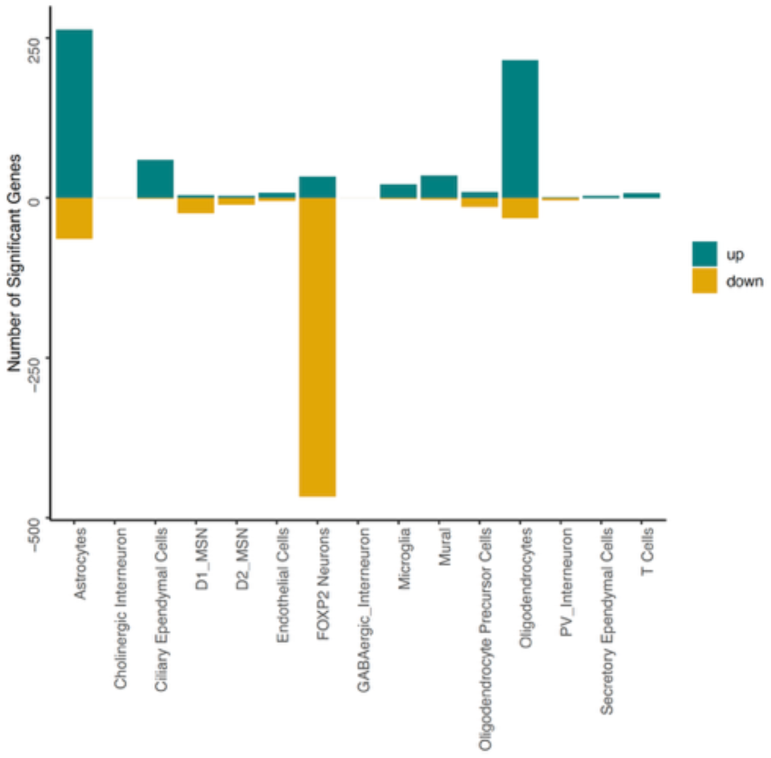
Number of up- and down-regulated genes in HD without Lim FOXP2 cells removed. Number of up- and down-regulated genes in HD with a cutoff of 0.05 from Benjamini-Hochberg corrected p-values.

**Supplementary Table 22.** Mean mapping scores for each cell type for each dataset’s control and HD individuals. Matsushima dataset is not included, because all datasets were mapped to that dataset.

| Cell Type | Lim<br>Control | Lim<br>HD | Garcia<br>Control | Garcia<br>HD | Handsaker<br>Control | Handsaker<br>HD |
| --- | --- | --- | --- | --- | --- | --- |
| Oligodendrocyte | 1.00 | 1.00 | 0.99 | 0.94 | 1.00 | 1.00 |
| Astrocyte | 1.00 | 0.99 | 0.99 | 0.98 | 1.00 | 1.00 |
| Oligodendrocyte<br>Precursor Cell | 1.00 | 0.99 | 1.00 | 0.98 | 1.00 | 1.00 |
| Cholinergic Interneuron | 0.56 | 0.53 | 0.71 | 0.72 | 0.59 | 0.54 |
| Ciliary Ependymal Cell | 0.97 | 0.96 | 0.99 | 0.99 | 0.99 | 0.99 |
| D1 MSN | 0.79 | 0.79 | 0.79 | 0.80 | 0.84 | 0.86 |
| D2 MSN | 0.83 | 0.68 | 0.81 | 0.66 | 0.81 | 0.76 |
| FOXP2 Neuron | 0.85 | 0.67 | 0.95 | 0.87 | 0.95 | 0.88 |
| Endothelial | 0.97 | 0.96 | 0.97 | 0.93 | 0.96 | 0.96 |
| GABAergic Interneuron | 0.89 | 0.63 | 0.96 | 0.72 | 0.97 | 0.95 |
| Microglia | 0.99 | 1.00 | 0.99 | 0.97 | 0.99 | 0.99 |
| Mural Cell | 0.95 | 0.98 | 0.98 | 0.95 | 0.98 | 0.98 |
| PV Interneuron | 0.93 | 0.99 | 0.99 | 0.95 | 0.99 | 0.99 |
| Secretory Ependymal Cell | 0.96 | 1.00 | 0.98 | 0.92 | 0.98 | 0.98 |
| T-Cell | 0.63 | 0.71 | 0.91 | 0.86 | 0.75 | 0.70 |

**Supplementary Table 23.** Mean nFeature for each cell type for each dataset’s control and HD individuals.

| Cell Type | Lim<br>Control | Lim<br>HD | Matsushima<br>Control | Matsushima<br>HD | Garcia<br>Control | Garcia<br>HD | Handsaker<br>Control | Handsaker<br>HD |
| --- | --- | --- | --- | --- | --- | --- | --- | --- |
| Oligodendrocyte | 2126 | 1829 | 1546 | 1531 | 1803 | 1618 | 1495 | 1462 |
| Astrocyte | 3698 | 2953 | 2831 | 2175 | 3618 | 2638 | 2493 | 2186 |
| Oligodendrocyte<br>Precursor Cell | 3036 | 2329 | 2368 | 2140 | 3159 | 2073 | 2192 | 2031 |
| Cholinergic Interneuron | 6025 | 1280 | 762 | 685 | 735 | 2292 | 752 | 985 |
| Ciliary Ependymal Cell | 4874 | 4484 | 2388 | 2366 | 3403 | 3208 | 2392 | 2495 |
| D1 MSN | 5241 | 940 | 4836 | 4576 | 5550 | 2947 | 4568 | 3856 |
| D2 MSN | 6664 | 4736 | 4698 | 4653 | 5393 | 4873 | 4881 | 4421 |
| FOXP2 Neuron | 4819 | NA | 3987 | 3856 | 4661 | 2607 | 4276 | 3231 |
| Endothelial | 4145 | 2359 | 2341 | 2215 | 4124 | 2345 | 1909 | 1733 |
| GABAergic Interneuron | 6934 | 2281 | 4640 | 3325 | 4757 | 4415 | 4923 | 4347 |
| Microglia | 2332 | 2029 | 1574 | 1555 | 2309 | 1600 | 1423 | 1520 |
| Mural Cell | 2723 | 2380 | 1786 | 1968 | 2197 | 1746 | 1644 | 1508 |
| PV Interneuron | 5994 | 4609 | 3919 | 3341 | 4308 | 2852 | 4621 | 3620 |
| Secretory Ependymal Cell | 5298 | 4253 | 3310 | 3511 | 3666 | 2767 | 3906 | 3230 |
| T-Cell | 1701 | 3006 | 1099 | 1393 | 1411 | 1287 | 1194 | 943 |

**Supplementary Table 24.** Mean nCount for each cell type for each dataset’s control and HD individuals.

| Cell Type | Lim<br>Control | Lim<br>HD | Matsushima<br>Control | Matsushima<br>HD | Garcia<br>Control | Garcia<br>HD | Handsaker<br>Control | Handsaker<br>HD |
| --- | --- | --- | --- | --- | --- | --- | --- | --- |
| Oligodendrocyte | 4720 | 4089 | 3032 | 3439 | 4056 | 3788 | 2879 | 2763 |
| Astrocyte | 10878 | 8437 | 6868 | 5862 | 10744 | 7838 | 5631 | 4734 |
| Oligodendrocyte<br>Precursor Cell | 7457 | 5866 | 5563 | 5597 | 8808 | 5543 | 4701 | 4352 |
| Cholinergic Interneuron | 30458 | 2487 | 1069 | 946 | 1000 | 4802 | 1378 | 1696 |
| Ciliary Ependymal Cell | 17048 | 14288 | 5337 | 5970 | 9593 | 11593 | 5196 | 5098 |
| D1 MSN | 28795 | 2738 | 19443 | 18925 | 28562 | 10931 | 16619 | 12680 |
| D2 MSN | 37503 | 28764 | 18800 | 20194 | 25518 | 23919 | 19427 | 16588 |
| FOXP2 Neuron | 24235 | NA | 14529 | 13846 | 19742 | 9449 | 15262 | 9506 |
| Endothelial | 17257 | 5697 | 5455 | 6006 | 16443 | 7987 | 3933 | 3292 |
| GABAergic Interneuron | 35416 | 10130 | 17175 | 10718 | 19583 | 15060 | 17301 | 14292 |
| Microglia | 5235 | 4532 | 2706 | 3051 | 5108 | 3361 | 2466 | 2613 |
| Mural Cell | 6991 | 5348 | 3324 | 4319 | 4667 | 4072 | 2977 | 2638 |
| PV Interneuron | 24842 | 16713 | 12650 | 11554 | 16016 | 9306 | 15105 | 10074 |
| Secretory Ependymal Cell | 18019 | 13237 | 8353 | 10362 | 10887 | 7327 | 10059 | 7609 |
| T-Cell | 3314 | 10005 | 1641 | 2561 | 2301 | 2351 | 2083 | 1405 |

**Supplementary Figure 21.**
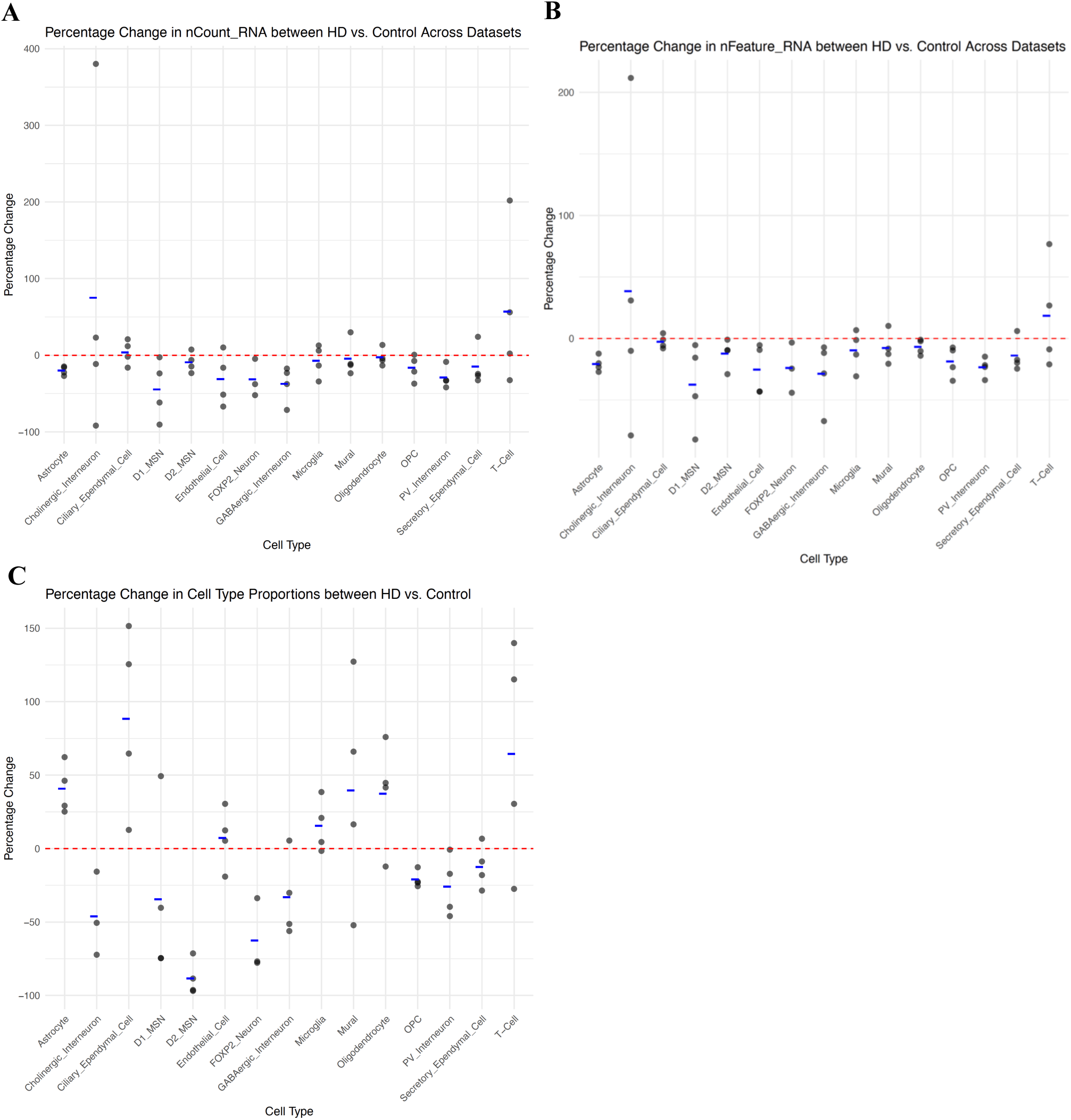
Scatterplot of percent change in A) mean nCount_RNA, B) nFeature_RNA, and C) cell type proportions for HD vs. Control in each data set for each cell type. Percentages above 0 represent an increase in HD relative to control. Blue line represents the mean for each cell type. Each point is a dataset. For cell type proportions, the cholinergic interneuron data from the Garcia dataset was removed due to it being a severe outlier (see Supplementary Table 25 for full numbers).

**Supplementary Table 25.** Percentage of each cell type for each dataset’s control and HD individuals.

| Cell Type | Lim<br>Control | Lim<br>HD | Matsushima<br>Control | Matsushima<br>HD | Garcia<br>Control | Garcia<br>HD | Handsaker<br>Control | Handsaker<br>HD |
| --- | --- | --- | --- | --- | --- | --- | --- | --- |
| Oligodendrocyte | 38.6 | 33.9 | 39.0 | 56.1 | 28.0 | 39.7 | 15.2 | 26.9 |
| Astrocyte | 12.6 | 16.3 | 11.6 | 18.8 | 15.7 | 22.9 | 20.1 | 25.3 |
| Oligodendrocyte<br>Precursor Cell | 4.4 | 3.2 | 4.3 | 3.3 | 4.6 | 3.5 | 7.5 | 6.6 |
| Cholinergic Interneuron | 0.2 | 0.2 | 0.2 | 0.0 | 0.3 | 7.2 | 0.2 | 0.1 |
| Ciliary Ependymal Cell | 0.7 | 1.2 | 1.0 | 2.4 | 5.1 | 5.8 | 1.5 | 3.3 |
| D1 MSN | 21.7 | 32.4 | 16.1 | 4.2 | 15.8 | 4.2 | 26.1 | 15.8 |
| D2 MSN | 10.9 | 0.4 | 13.4 | 1.5 | 12.3 | 0.4 | 13.3 | 3.7 |
| FOXP2 Neuron | 2.9 | NA | 2.0 | 0.5 | 4.1 | 0.9 | 3.1 | 2.0 |
| Endothelial | 0.3 | 0.2 | 0.7 | 0.8 | 2.2 | 2.9 | 0.9 | 1.0 |
| GABAergic Interneuron | 0.1 | 0.1 | 0.3 | 0.1 | 0.4 | 0.2 | 0.3 | 0.3 |
| Microglia | 5.2 | 5.1 | 6.9 | 8.3 | 7.2 | 7.5 | 7.8 | 10.7 |
| Mural Cell | 0.3 | 0.2 | 0.8 | 1.3 | 1.0 | 2.3 | 1.0 | 1.2 |
| PV Interneuron | 0.9 | 0.7 | 2.1 | 1.3 | 1.8 | 1.0 | 1.6 | 1.6 |
| Secretory Ependymal Cell | 1.0 | 0.8 | 1.4 | 1.0 | 1.3 | 1.2 | 1.3 | 1.4 |
| T-Cell | 0.0 | 0.0 | 0.1 | 0.3 | 0.2 | 0.4 | 0.1 | 0.0 |

## References

1 Schirmer, L. et al. Neuronal vulnerability and multilineage diversity in multiple sclerosis. Nature 573, 75–82 (2019).

2 Jäkel, S. et al. Altered human oligodendrocyte heterogeneity in multiple sclerosis. Nature 566, 543–547 (2019).

3 Kihara, Y. et al. Single-nucleus RNA-seq of normal-appearing brain regions in relapsing-remitting vs. secondary progressive multiple sclerosis: implications for the efficacy of fingolimod. Frontiers in Cellular Neuroscience 16, 918041 (2022).

4 Ruzicka, W. B. et al. Single-cell multi-cohort dissection of the schizophrenia transcriptome. Science 384, eadg5136 (2024).

5 Batiuk, M. Y. et al. Upper cortical layer–driven network impairment in schizophrenia. Science Advances 8, eabn8367 (2022).

6 Ling, E. et al. A concerted neuron–astrocyte program declines in ageing and schizophrenia. Nature 627, 604–611 (2024).

7 Nagy, C. et al. Single-nucleus transcriptomics of the prefrontal cortex in major depressive disorder implicates oligodendrocyte precursor cells and excitatory neurons. Nature neuroscience 23, 771–781 (2020).

8 Velmeshev, D. et al. Single-cell genomics identifies cell type–specific molecular changes in autism. Science 364, 685–689 (2019).

9 Gandal, M. J. et al. Broad transcriptomic dysregulation occurs across the cerebral cortex in ASD. Nature 611, 532–539 (2022).

10 Smajić, S. et al. Single-cell sequencing of human midbrain reveals glial activation and a Parkinson-specific neuronal state. Brain 145, 964–978 (2022).

11 Kamath, T. et al. Single-cell genomic profiling of human dopamine neurons identifies a population that selectively degenerates in Parkinson’s disease. Nature neuroscience 25, 588–595 (2022).

12 Martirosyan, A. et al. Unravelling cell type-specific responses to Parkinson’s Disease at single cell resolution. Molecular neurodegeneration 19, 1–24 (2024).

13 Lee, A. J. et al. Characterization of altered molecular mechanisms in Parkinson’s disease through cell type–resolved multiomics analyses. Science Advances 9, eabo2467 (2023).

14 Adams, L., Song, M. K., Yuen, S., Tanaka, Y. & Kim, Y.-S. A single-nuclei paired multiomic analysis of the human midbrain reveals age-and Parkinson’s disease– associated glial changes. Nature Aging 4, 364–378 (2024).

15 Wang, Q. et al. Molecular profiling of human substantia nigra identifies diverse neuron types associated with vulnerability in Parkinson’s disease. Science advances 10, eadi8287 (2024).

16 van den Oord, E. J., Xie, L. Y., Zhao, M., Aberg, K. A. & Clark, S. L. A single-nucleus transcriptomics study of alcohol use disorder in the nucleus accumbens. Addiction biology 28, e13250 (2023).

17 Brenner, E. et al. Single cell transcriptome profiling of the human alcohol-dependent brain. Human Molecular Genetics 29, 1144–1153 (2020).

18 Renthal, W. et al. Characterization of human mosaic Rett syndrome brain tissue by single-nucleus RNA sequencing. Nature neuroscience 21, 1670–1679 (2018).

19 Mitroi, D. N., Tian, M., Kawaguchi, R., Lowry, W. E. & Carmichael, S. T. Single-nucleus transcriptome analysis reveals disease-and regeneration-associated endothelial cells in white matter vascular dementia. Journal of Cellular and Molecular Medicine 26, 3183–3195 (2022).

20 Lee, H. et al. Cell type-specific transcriptomics reveals that mutant huntingtin leads to mitochondrial RNA release and neuronal innate immune activation. Neuron 107, 891–908. e898 (2020).

21 Matsushima, A. et al. Transcriptional vulnerabilities of striatal neurons in human and rodent models of Huntington’s disease. Nature communications 14, 282 (2023).

22 Al-Dalahmah, O. et al. Single-nucleus RNA-seq identifies Huntington disease astrocyte states. Acta neuropathologica communications 8, 1–21 (2020).

23 Lim, R. G. et al. Huntington disease oligodendrocyte maturation deficits revealed by single-nucleus RNAseq are rescued by thiamine-biotin supplementation. Nature Communications 13, 7791 (2022).

24 Fujita, M. et al. Cell subtype-specific effects of genetic variation in the Alzheimer’s disease brain. Nature Genetics, 1–10 (2024).

25 Su, Y. et al. Multi-omics resolves a sharp disease-state shift between mild and moderate COVID-19. Cell 183, 1479–1495. e1420 (2020).

26 Hoffman, G. E. et al. Efficient differential expression analysis of large-scale single cell transcriptomics data using dreamlet. bioRxiv, 2023.2003. 2017.533005 (2023).

27 Stephenson, E. et al. Single-cell multi-omics analysis of the immune response in COVID-19. Nature medicine 27, 904–916 (2021).

28 Squair, J. W. et al. Confronting false discoveries in single-cell differential expression. Nature communications 12, 1–15 (2021).

29 Murphy, A. E., Fancy, N. & Skene, N. Avoiding false discoveries in single-cell RNA-seq by revisiting the first Alzheimer’s disease dataset. Elife 12, RP90214 (2023).

30 Love, M. I., Huber, W. & Anders, S. Moderated estimation of fold change and dispersion for RNA-seq data with DESeq2. Genome biology 15, 1–21 (2014).

31 Cembrowski, M. S. Single-cell transcriptomics as a framework and roadmap for understanding the brain. Journal of neuroscience methods 326, 108353 (2019).

32 Wendt, F. R., Pathak, G. A., Tylee, D. S., Goswami, A. & Polimanti, R. Heterogeneity and polygenicity in psychiatric disorders: a genome-wide perspective. Chronic Stress 4, 2470547020924844 (2020).

33 Marigorta, U. M., Rodríguez, J. A., Gibson, G. & Navarro, A. Replicability and prediction: lessons and challenges from GWAS. Trends in Genetics 34, 504–517 (2018).

34 Willer, C. J., Li, Y. & Abecasis, G. R. METAL: fast and efficient meta-analysis of genomewide association scans. Bioinformatics 26, 2190–2191 (2010).

35 Mägi, R. & Morris, A. P. GWAMA: software for genome-wide association meta-analysis. BMC bioinformatics 11, 1–6 (2010).

36 Evangelou, E. & Ioannidis, J. P. Meta-analysis methods for genome-wide association studies and beyond. Nature Reviews Genetics 14, 379–389 (2013).

37 Bakken, T. E. et al. Comparative cellular analysis of motor cortex in human, marmoset and mouse. Nature 598, 111–119 (2021).

38 Hao, Y. et al. Integrated analysis of multimodal single-cell data. Cell 184, 3573–3587. e3529 (2021).

39 Junttila, S., Smolander, J. & Elo, L. L. Benchmarking methods for detecting differential states between conditions from multi-subject single-cell RNA-seq data. Briefings in bioinformatics 23, bbac286 (2022).

40 Murdock, M. H. & Tsai, L.-H. Insights into Alzheimer’s disease from single-cell genomic approaches. Nature Neuroscience, 1–15 (2023).

41 Andreatta, M. & Carmona, S. J. UCell: Robust and scalable single-cell gene signature scoring. Computational and structural biotechnology journal 19, 3796–3798 (2021).

42 Schwarzer, G., Carpenter, J. R. & Rücker, G. Meta-analysis with R. Vol. 4784 (Springer, 2015).

43 Kim, M. C. et al. Method of moments framework for differential expression analysis of single-cell RNA sequencing data. Cell 187, 6393–6410. e6316 (2024).

44 Conway, J. R., Lex, A. & Gehlenborg, N. UpSetR: an R package for the visualization of intersecting sets and their properties. Bioinformatics 33, 2938–2940 (2017).

45 Mathys, H. et al. Single-cell transcriptomic analysis of Alzheimer’s disease. Nature 570, 332–337 (2019).

46 Gabitto, M. I. et al. Integrated multimodal cell atlas of Alzheimer’s disease. bioRxiv, 2023.2005. 2008.539485 (2023).

47 Hao, Y. et al. Dictionary learning for integrative, multimodal and scalable single-cell analysis. Nature Biotechnology, 1–12 (2023).

48 Fujita, M. et al. Cell-subtype specific effects of genetic variation in the aging and Alzheimer cortex. bioRxiv, 2022.2011. 2007.515446 (2022).

49 Mathys, H. et al. Single-cell atlas reveals correlates of high cognitive function, dementia, and resilience to Alzheimer’s disease pathology. Cell 186, 4365–4385. e4327 (2023).

50 Subramanian, A. et al. Gene set enrichment analysis: a knowledge-based approach for interpreting genome-wide expression profiles. Proceedings of the National Academy of Sciences 102, 15545–15550 (2005).

51 Mathys, H. et al. Single-cell multiregion dissection of Alzheimer’s disease. Nature 632, 858–868 (2024).

52 Xiong, X. et al. Epigenomic dissection of Alzheimer’s disease pinpoints causal variants and reveals epigenome erosion. Cell 186, 4422–4437. e4421 (2023).

53 Wu, T. et al. clusterProfiler 4.0: A universal enrichment tool for interpreting omics data. The Innovation 2, 100141 (2021).

54 Jiang, L. et al. Systematic reconstruction of molecular pathway signatures using scalable single-cell perturbation screens. bioRxiv, 2024.2001. 2029.576933 (2024).

55 Kim, J. J. et al. Multi-ancestry genome-wide association meta-analysis of Parkinson’s disease. Nature genetics 56, 27–36 (2024).

56 Cattaneo, A. et al. The expression of VGF is reduced in leukocytes of depressed patients and it is restored by effective antidepressant treatment. Neuropsychopharmacology 35, 1423–1428 (2010).

57 Giusto, E. et al. Prospective role of PAK6 and 14-3-3γ as biomarkers for Parkinson’s disease. Journal of Parkinson’s Disease, 1–12 (2024).

58 Pogoda, A., Chmielewska, N., Maciejak, P. & Szyndler, J. Transcriptional dysregulation in Huntington’s disease: the role in pathogenesis and potency for pharmacological targeting. Current medicinal chemistry 28, 2783–2806 (2021).

59 Hachigian, L. J. et al. Control of Huntington’s disease-associated phenotypes by the striatum-enriched transcription factor Foxp2. Cell reports 21, 2688–2695 (2017).

60 Rodríguez-Urgellés, E. et al. Postnatal Foxp2 regulates early psychiatric-like phenotypes and associated molecular alterations in the R6/1 transgenic mouse model of Huntington’s disease. Neurobiology of disease 173, 105854 (2022).

61 Miao, J. et al. Microglia in Alzheimer’s disease: Pathogenesis, mechanisms, and therapeutic potentials. Frontiers in aging neuroscience 15, 1201982 (2023).

62 Szklarczyk, D. et al. The STRING database in 2023: protein–protein association networks and functional enrichment analyses for any sequenced genome of interest. Nucleic acids research 51, D638–D646 (2023).

63 Bellenguez, C. et al. New insights into the genetic etiology of Alzheimer’s disease and related dementias. Nature genetics 54, 412–436 (2022).

64 Wightman, D. P. et al. A genome-wide association study with 1,126,563 individuals identifies new risk loci for Alzheimer’s disease. Nature genetics 53, 1276–1282 (2021).

65 De Rojas, I. et al. Common variants in Alzheimer’s disease and risk stratification by polygenic risk scores. Nature communications 12, 3417 (2021).

66 Xi, M. et al. Therapeutic potential of phosphodiesterase inhibitors for cognitive amelioration in Alzheimer’s disease. European Journal of Medicinal Chemistry 232, 114170 (2022).

67 Sikora, J. et al. Quetiapine and novel PDE10A inhibitors potentiate the anti-BuChE activity of donepezil. Journal of Enzyme Inhibition and Medicinal Chemistry 35, 1743–1750 (2020).

68 Kageyama, R., Ohtsuka, T. & Kobayashi, T. Roles of Hes genes in neural development. Development, growth & differentiation 50, S97–S103 (2008).

69 Bai, G. et al. Epigenetic dysregulation of hairy and enhancer of split 4 (HES4) is associated with striatal degeneration in postmortem Huntington brains. Human molecular genetics 24, 1441–1456 (2015).

70 Bozdagi, O. et al. The neurotrophin-inducible gene Vgf regulates hippocampal function and behavior through a brain-derived neurotrophic factor-dependent mechanism. Journal of Neuroscience 28, 9857–9869 (2008).

71 Ali, M. & Bracko, O. VEGF paradoxically reduces cerebral blood flow in Alzheimer’s disease mice. Neuroscience Insights 17, 26331055221109254 (2022).

72 De Schepper, S. et al. Perivascular cells induce microglial phagocytic states and synaptic engulfment via SPP1 in mouse models of Alzheimer’s disease. Nature Neuroscience 26, 406–415 (2023).

73 Gurses, M. S., Ural, M. N., Gulec, M. A., Akyol, O. & Akyol, S. Pathophysiological function of ADAMTS enzymes on molecular mechanism of Alzheimer’s disease. Aging and disease 7, 479 (2016).

74 Nandi, A., Yan, L.-J., Jana, C. K. & Das, N. Role of catalase in oxidative stress-and age- associated degenerative diseases. Oxidative medicine and cellular longevity 2019, 9613090 (2019).

75 Nell, H. J. et al. Targeted antioxidant, catalase–SKL, reduces beta-amyloid toxicity in the rat brain. Brain Pathology 27, 86–94 (2017).

76 Forner, S. et al. Systematic phenotyping and characterization of the 5xFAD mouse model of Alzheimer’s disease. Scientific data 8, 270 (2021).

77 Siddik, M. A. B. et al. Branched-chain amino acids are linked with Alzheimer’s disease-related pathology and cognitive deficits. Cells 11, 3523 (2022).

78 Nong, X. et al. The mechanism of branched-chain amino acid transferases in different diseases: Research progress and future prospects. Frontiers in Oncology 12, 988290 (2022).

79 Bis, J. C. et al. Whole exome sequencing study identifies novel rare and common Alzheimer’s-Associated variants involved in immune response and transcriptional regulation. Molecular psychiatry 25, 1859–1875 (2020).

80 Holstege, H. et al. Exome sequencing identifies rare damaging variants in ATP8B4 and ABCA1 as risk factors for Alzheimer’s disease. Nature Genetics, 1–9 (2022).

81 Prokopenko, D. et al. Whole-genome sequencing reveals new Alzheimer’s disease– associated rare variants in loci related to synaptic function and neuronal development. Alzheimer’s & Dementia 17, 1509–1527 (2021).

82 Guo, L., Zhong, M. B., Zhang, L., Zhang, B. & Cai, D. Sex differences in Alzheimer’s disease: Insights from the multiomics landscape. Biological psychiatry 91, 61–71 (2022).

83 Zhao, S., Ye, B., Chi, H., Cheng, C. & Liu, J. Identification of peripheral blood immune infiltration signatures and construction of monocyte-associated signatures in ovarian cancer and Alzheimer’s disease using single-cell sequencing. Heliyon 9 (2023).

84 Patel, H., Dobson, R. J. & Newhouse, S. J. A meta-analysis of Alzheimer’s disease brain transcriptomic data. Journal of Alzheimer’s Disease 68, 1635–1656 (2019).

85 Tian, Y. et al. Identification of diagnostic signatures associated with immune infiltration in Alzheimer’s disease by integrating bioinformatic analysis and machine-learning strategies. Frontiers in Aging Neuroscience 14, 919614 (2022).

86 Walters, S. et al. Associations of sex, race, and apolipoprotein e alleles with multiple domains of cognition among older adults. JAMA neurology 80, 929–939 (2023).

87 Grove, J. et al. Identification of common genetic risk variants for autism spectrum disorder. Nature genetics 51, 431–444 (2019).

88 Langfelder, P. & Horvath, S. WGCNA: an R package for weighted correlation network analysis. BMC bioinformatics 9, 1–13 (2008).

89 Johnson, E. C. et al. Large-scale deep multi-layer analysis of Alzheimer’s disease brain reveals strong proteomic disease-related changes not observed at the RNA level. Nature neuroscience 25, 213–225 (2022).

90 Loh, P.-R., Kichaev, G., Gazal, S., Schoech, A. P. & Price, A. L. Mixed-model association for biobank-scale datasets. Nature genetics 50, 906–908 (2018).

91 Uffelmann, E. et al. Genome-wide association studies. Nature Reviews Methods Primers 1, 59 (2021).

92 Li, Y. et al. Analyzing bivariate cross-trait genetic architecture in GWAS summary statistics with the BIGA cloud computing platform. bioRxiv, 2023.2004. 2028.538585 (2023).

93 Plenge, R. M., Scolnick, E. M. & Altshuler, D. Validating therapeutic targets through human genetics. Nature reviews Drug discovery 12, 581–594 (2013).

94 Otero-Garcia, M. et al. Molecular signatures underlying neurofibrillary tangle susceptibility in Alzheimer’s disease. Neuron 110, 2929–2948. e2928 (2022).

95 Leng, K. et al. Molecular characterization of selectively vulnerable neurons in Alzheimer’s disease. Nature neuroscience 24, 276–287 (2021).

96 Zhou, Y. et al. Human and mouse single-nucleus transcriptomics reveal TREM2-dependent and TREM2-independent cellular responses in Alzheimer’s disease. Nature medicine 26, 131–142 (2020).

97 Grubman, A. et al. A single-cell atlas of entorhinal cortex from individuals with Alzheimer’s disease reveals cell-type-specific gene expression regulation. Nature neuroscience 22, 2087–2097 (2019).

98 Morabito, S. et al. Single-nucleus chromatin accessibility and transcriptomic characterization of Alzheimer’s disease. Nature genetics 53, 1143–1155 (2021).

99 Lau, S.-F., Cao, H., Fu, A. K. & Ip, N. Y. Single-nucleus transcriptome analysis reveals dysregulation of angiogenic endothelial cells and neuroprotective glia in Alzheimer’s disease. Proceedings of the National Academy of Sciences 117, 25800–25809 (2020).

100 Yang, A. C. et al. A human brain vascular atlas reveals diverse mediators of Alzheimer’s risk. Nature 603, 885–892 (2022).

101 Sayed, F. A. et al. AD-linked R47H-TREM2 mutation induces disease-enhancing microglial states via AKT hyperactivation. Science translational medicine 13, eabe3947 (2021).

102 Gerrits, E. et al. Distinct amyloid-β and tau-associated microglia profiles in Alzheimer’s disease. Acta neuropathologica 141, 681–696 (2021).

103 Smith, A. M. et al. Diverse human astrocyte and microglial transcriptional responses to Alzheimer’s pathology. Acta neuropathologica 143, 75–91 (2022).

104 Sadick, J. S. et al. Astrocytes and oligodendrocytes undergo subtype-specific transcriptional changes in Alzheimer’s disease. Neuron 110, 1788–1805. e1710 (2022).

105 Garcia, F. J. et al. Single-cell dissection of the human brain vasculature. Nature 603, 893–899 (2022).

106 Handsaker, R. E. et al. Long somatic DNA-repeat expansion drives neurodegeneration in Huntington’s disease. Cell 188, 623–639. e619 (2025).

107 Barker, S. J. et al. MEF2 is a key regulator of cognitive potential and confers resilience to neurodegeneration. Science Translational Medicine 13, eabd7695 (2021).

108 Tian, Y. et al. Single-cell immunology of SARS-CoV-2 infection. Nature biotechnology 40, 30–41 (2022).

109 Smith, G. Limma: linear models for microarray data. Bioinformatics and Computational Biology Solutions using R and Bioconductor. *Springer*, New York, 397–420 (2005).

110 Quijano Xacur, O. A. The unifed distribution. Journal of Statistical Distributions and Applications 6, 1–12 (2019).

111 Robin, X. et al. pROC: an open-source package for R and S+ to analyze and compare ROC curves. BMC bioinformatics 12, 1–8 (2011).

112 Lawrence, M. et al. Software for computing and annotating genomic ranges. PLoS computational biology 9, e1003118 (2013).

113 Morgan, M. & Shepherd, L. AnnotationHub: Client to access AnnotationHub resources. R package version 2, 2017 (2017).

114 Perez, G. et al. The UCSC Genome Browser database: 2025 update. Nucleic Acids Research 53, D1243–D1249 (2025).

115 Shannon, P. et al. Cytoscape: a software environment for integrated models of biomolecular interaction networks. Genome research 13, 2498–2504 (2003).

116 Crowell, H. L. et al. Muscat detects subpopulation-specific state transitions from multi-sample multi-condition single-cell transcriptomics data. Nature communications 11, 6077 (2020).

